# Targeting the BAG-1 family of co-chaperones in lethal prostate cancer

**DOI:** 10.1101/2022.10.17.512378

**Authors:** Antje Neeb, Ines Figueiredo, Denisa Bogdan, Laura Cato, Jutta Stober, Juan M. Jimenez-Vacas, Victor Gourain, Irene I. Lee, Rebecca Seeger, Claudia Muhle-Goll, Bora Gurel, Jonathan Welti, Daniel Nava Rodrigues, Jan Rekowski, Xintao Qiu, Yija Jiang, Patrizio Di Micco, Borja Mateos, Stasė Bielskutė, Ruth Riisnaes, Ana Ferreira, Susana Miranda, Mateus Crespo, Lorenzo Buroni, Jian Ning, Stefan Bräse, Nicole Jung, Simone Gräßle, Daniel Metzger, Amanda Swain, Xavier Salvatella, Stephen R. Plymate, Bissan Al-Lazikani, Henry Long, Wei Yuan, Myles Brown, Andrew C. B. Cato, Johann S. de Bono, Adam Sharp

## Abstract

Therapies that abrogate persistent androgen receptor (AR) signaling in castration resistant prostate cancer (CRPC) remain an unmet clinical need. The N-terminal domain (NTD) of the AR drives transcriptional activity in CRPC but is intrinsically disordered and remains a challenging therapeutic target. Therefore, inhibiting critical co-chaperones, such as BAG-1L, is an attractive alternative strategy. We performed druggability analyses demonstrating the BAG domain to be a challenging drug target. Thio-2, a tool compound, has been reported to bind the BAG domain of BAG-1L and inhibit BAG-1L-mediated AR transactivation. However, despite these data, the mechanism of action of Thio-2 is poorly understood and the BAG domain which is present in all BAG-1 isoforms has not been validated as a therapeutic target. Herein, we demonstrate growth inhibiting activity of Thio-2 in CRPC cell lines and patient derived models with decreased AR genomic binding and AR signaling independent of BAG-1 isoform function. Furthermore, genomic abrogation of BAG-1 isoforms did not recapitulate the described Thio-2 phenotype, and NMR studies suggest that Thio-2 may bind the AR NTD, uncovering a potential alternative mechanism of action, although in the context of low compound solubility. Furthermore, BAG-1 isoform knockout mice are viable and fertile, in contrast to previous studies, and when crossed with prostate cancer mouse models, BAG-1 deletion does not significantly impact prostate cancer development and growth. Overall, these data demonstrate that Thio-2 inhibits AR signaling and growth in CRPC independent of BAG-1 isoforms, and unlike previous studies of the activated AR, therapeutic targeting of the BAG domain requires further validation before being considered a therapeutic strategy for the treatment of CRPC.

## Introduction

Prostate cancer is the most commonly diagnosed non-cutaneous malignancy in men and is a leading cause of male mortality; patients with advanced disease have a poor prognosis with a 5-year overall survival of 31% (1). The androgen receptor (AR) remains the major therapeutic target for both advanced castration sensitive prostate cancer (CSPC) and castration resistant prostate cancer (CRPC) (2, 3). Primary and secondary resistance to therapies targeting the AR signaling axis remains inevitable, driven, in part, by ongoing AR signaling through AR amplification, AR mutations, and constitutively active AR splice variants (3–5). The development of novel therapies that block persistent AR signaling is an unmet clinical need.

Activity of the inhibited AR, and AR splice variant that emerge during the development of CRPC, has been reported to be driven by the constitutively active AR N-terminal domain (NTD), one of the largest intrinsically disordered polypeptides and challenging therapeutic target (6). One attractive therapeutic strategy is to target molecular co-chaperones, such as BAG-1 (BCL-2-associated athanogene-1), reported to be critical for AR signaling. BAG-1 interacts with a wide range of molecular targets to regulate multiple cellular pathways (including apoptosis, proliferation, metastasis, and nuclear hormone receptor transactivation) important for the development and progression of cancer (7–9). Three major isoforms, BAG-1L (50kDa), BAG-1M (46kDa) and BAG-1S (36kDa), exist in humans and are generated through alternative initiation of translation from a single mRNA (10). Consistent with this, BAG-1L has a unique N-terminus which contains a nuclear localization sequence and is predominantly localized within the nucleus, supporting its interaction with the activated AR, whereas the other isoforms (BAG-1M and BAG-1S) are found in both the nucleus and cytoplasm (7–9). All BAG-1 isoforms share a common C-terminus, which contains the highly conserved BAG domain, critical for the interaction between BAG-1 isoforms and the heat shock chaperones, HSC70/HSP70 (11–13). Importantly, the BAG-1:HSC70/HSP70 interaction is critical for BAG-1 function; therapies targeting this interaction are an attractive strategy to overcome BAG-1 activity in cancer (7–9, 14–21).

BAG-1L plays a critical role in transactivation of the AR, and nuclear BAG-1 protein expression correlates with important clinical characteristics (14–17, 22–24). Through its C-terminal BAG domain, BAG-1L binds to the AR NTD, leading to receptor transactivation (14–17). Consistent with this, loss of BAG-1L abrogates AR signaling and reduces prostate cancer growth (14). In addition, expression of nuclear BAG-1 correlates with worse outcome from AR targeting therapies in patients with CRPC (14). Moreover, mutagenesis studies demonstrated that specific amino acid residues within the BAG domain of BAG-1L are critical for the BAG-1L:AR interaction and AR transactivation (14). Finally, Thio-2, a tool compound that has been predicted to bind the BAG domain within BAG-1 isoforms through *in-silico* docking experiments, inhibited BAG-1L mediated AR transactivation (14, 25). More recently, due to concerns regarding Thio-2 specificity, A4B17 has been developed (26). Similar to Thio-2, A4B17 disrupts the BAG-1L:AR NTD interaction and suppresses AR target gene expression demonstrating efficacy in pre-clinical prostate cancer models (27). Taken together, these data support targeting the BAG domain of BAG-1L as an attractive therapeutic strategy to overcome persistent AR signaling in CRPC.

Despite these promising data, it is important to note that the impact of Thio-2 on AR signaling and prostate cancer growth in CRPC has not been shown to be mediated through the BAG domain. In addition, therapies that target the BAG domain would block the function of all three BAG-1 isoforms (S, M and L), and this has not been validated as a therapeutic strategy in CRPC with nearly all studies focusing on the function of BAG-1L and the activated AR. These studies are critically important and required to further determine whether the BAG domain of the BAG-1 isoforms should be considered for drug discovery and development efforts in CRPC. Herein, we confirm that Thio-2 inhibits AR signaling and growth in CRPC models through a BAG-1 isoform-independent mechanism. In addition, genomic abrogation of BAG-1 isoforms in CRPC models, BAG-1 isoform mouse knockout models, and quantification of BAG-1L expression in prostate cancer tissue, suggest that BAG-1 isoforms may play a limited role in regulating AR signaling in CRPC. Finally, Thio-2 may function through an interaction with the AR NTD in a BAG-1 independent manner, but its poor solubility suggests it will be a challenging starting point for drug development.

## Materials and Methods

### In-vitro patient derived xenograft organoid proliferation studies

CP50, CP89 and CP142 patient derived xenografts (PDXs) have been previously described (28–30). PDX-organoids (PDX-Os) were cultured, and proliferation measured as previously described (28, 31). Briefly PDX tumours were harvested in PDX harvesting solution (adDMEM/F12 containing 10 μM ROCK inhibitor Y27632 (Selleck Chemicals), penicillin/streptomycin, 10 mM Hepes and GlutaMAX 100× diluted all purchased form Thermofisher), cut into small pieces (< 3mm^3^) and single cell suspensions were generated by mechanical separation (40 μm Corning cell strainer, Sigma-Aldrich). Pellets were washed once on ice-cold PBS/5 mM EDTA/1x GlutaMax/10 μM Y27632, and red blood cells were removed using red blood cell lysis buffer (0.8% NH4Cl in 0.1 mM EDTA in water, buffered with KHCO3 to pH of 7.2 - 7.6, incubated 1-minute on ice) followed by another wash with ice cold PBS/5 mM EDTA/1x GlutaMax/10 μM Y27632. Single cell suspensions were either frozen for later use in BioCat BambankerTM freezing medium (Fisher Scientific) supplemented with 10 μM Y27632, or directly resuspended in ice-cold organoid growth medium (as published by Drost and colleagues with the following alterations: The p38 inhibitor SB202190 was replaced by the addition of 5 nM of NRG1) and subsequently diluted in one volume of phenol red-free, growth factor reduced, Corning MatrigelTM (Fisher Scientific) (28, 32). Organoid domes (5-50 μl) were plated as previously described by Drost and colleagues and topped up with warm medium after solidification (32). Cultures were observed over 3-7 days until visible organoid formation could be observed and then re-seeded for actual experiment. Immunohistochemistry was performed to confirm the presence of BAG-1 and AR in the developing organoids. For drug treatment, organoids were harvested in organoid harvesting solution (Amsbio), washed with medium and re-plated in the same way as described above and incubated in organoid growth medium for 5 days following drug treatment. CellTiter-Glo® 3D Cell Viability Assay (Promega) was used to assay growth of the organoids according to the manufacturer’s instruction and luminescence was measured using Synergy HTX (BioTek).

### Immunohistochemistry

For the Institute of Cancer Research/Royal Marsden Hospital (ICR/RMH) immunohistochemistry (IHC) studies, androgen receptor full length (AR-FL) IHC was performed as previously described (5). BAG-1L (human, rabbit monoclonal, RM310, RevMAb), panBAG-1 (human, rabbit monoclonal, RM356, RevMAb), panBAG-1 (mouse, goat polyclonal, AF815, R&D systems), and AR-FL (mouse/human, rabbit monoclonal, EPR1535(2), abcam) were all validated and optimized for IHC in this study. Specific details on the IHC assays developed are detailed in **Supplementary Table 1**. For the University of Washington/Fred Hutchinson Cancer Research Center (UW/FHCRC) BAG-1L IHC, antigen retrieval was performed in 10 mM citrate buffer (pH 6.0) in a pressure cooker for 30 minutes. Endogenous peroxidase and avidin/biotin were blocked respectively (Vector Laboratories). Sections were then blocked with 5% normal goat–horse–chicken serum, incubated with primary anti-BAG-1L (human, rabbit monoclonal, RM310, RevMAb) antibody (1:500), incubated with biotinylated secondary antibody (Vector Laboratories), followed by ABC reagent (Vector Laboratories) and stable DAB (Invitrogen). All sections were lightly counterstained with hematoxylin and mounted with Cytoseal XYL (Richard-Allan Scientific). Rabbit IgG (Vector Laboratories) were used as negative controls at the same concentration as the primary antibody.

### IHC quantification

For both ICR/RMH and UW/FHCRC studies, BAG-1L, mouse pan BAG-1 and mouse AR-FL IHC, nuclear and cytoplasmic quantification for each sample was determined by a pathologist blinded to clinical and molecular data using modified H-scores ([% of negative staining x 0] + [% of weak staining x 1] + [% of moderate staining x 2] + [% of strong staining x 3]), to determine the overall percentage of positivity across the entire stained samples, yielding a range from 0 to 300 (33).

### Western blotting

Cell line, PDX and PDX-O were lysed with RIPA buffer (Pierce™; ThermoFisher) supplemented with protease inhibitor cocktail (Roche; Sigma-Aldrich) and PhosStop phosphatase inhibitor mix (Roche; Sigma-Aldrich). PDX lysate was obtained by mechanical homogenization using a Qiagen TissueLyser homogenizer according to the manufacturer’s instructions. Protein extracts (25 μg) were sonicated, heated for 5 minutes at 95C and separated on 4-12% NuPAGE® Bis-Tris gel (Invitrogen) by electrophoresis and subsequently transferred onto Immobilon-P™ PVDF membranes of 0.45 μm pore size (Millipore®; Merck). Details of primary antibodies used are provided in **Supplementary Table 2**. Chemiluminescence was detected on the Chemidoc™ Touch imaging system (Bio-Rad).

### RNA extraction

RNA from knockout mice was obtained by mechanical homogenization in PeqGold, RNApure solution (VWR), following the manufacturer’s instruction or, reconstituted with RNeasy RLT buffer, passed through a Qiashredder tube (Qiagen), and further processed with RNeasy Plus Mini Kit as per manufacturer’s instructions. Cell line, PDX and PDX-O RNA was extracted using the RNeasy Plus Mini Kit (Qiagen) as per manufacturer’s instructions.

### Quantitative reverse transcription PCR (qRT-PCR)

cDNA was synthesized using the Revertaid First Strand cDNA Synthesis kit (ThermoFisher). qRT-PCR was carried out using a ViiA 7 Real-Time PCR System (Life Technologies) using the TaqMan Universal PCR Master Mix (Applied Biosystems). TaqMan probes (ThermoFisher) used are listed in **Supplementary Table 3**. Fold change in mRNA expression levels were calculated by the comparative Ct method, using the formula 2-(-(ΔΔCt). Cell line and PDX-O samples were normalized against the average of four (GAPDH, B2M, HRPT1 and RPLP0) housekeeping genes and knockout mouse samples were normalized to mouse GAPDH.

### In-vivo Thio-2 toxicity studies

Non tumor bearing NSG male mice were treated with vehicle (5% DMSO in 10% (w/v) HBC (2-hydroxypropyl-β-cyclodextrin) in 0.9% saline) or 15mg/kg Thio-2 by once daily intraperitoneal injection for 5 days with daily weights. Following 5 days treatment mice were sacrificed and organ (heart, kidney, testes, seminal vesicles, and prostate) weights were determined.

### In-vivo PDX studies

CP50 PDX fragments were grafted subcutaneously into NSG male mice and drug treatment commenced with vehicle (5% DMSO in 10% (w/v) HBC (2-hydroxypropyl-β-cyclodextrin) in 0.9% saline) or 15 mg/kg Thio-2 by once daily intraperitoneal injection when tumors reached a size of 300 to 400 mm^3^. Mice were treated daily for 14 days and tumor measurements were taken every 2-3 days (grouped by 5-day intervals). After 14 days treatment mice were sacrificed, and plasma and tumors were collected for pharmacodynamic analyses.

### In-vivo PDX serum PSA analyses

Serum was separated by 5min centrifugation at 9000rpm from blood collected from mice by cardiac puncture under general terminal anesthesia after blood clotting was allowed to take place for 15min. Serum PSA was analyzed in 1:100 diluted serum using the human PSA SimpleStepTM ELISA kit (Abcam) following the manufacturer’s instructions.

### RNA-sequencing and analysis (cell line)

For unstimulated experiments, LNCaP cells were grown in full media (10% fetal bovine serum) prior to treatment with vehicle (DMSO 0.1 %) or 50 μM Thio-2 for 17 hours. For stimulated experiments, LNCaP cells were grown in starved media (10% charcoal stripped serum) for 72 hours prior to treatment with vehicle (DMSO 0.1 %) or 5 μM Thio-2. Following 1 hour pre-treatment with vehicle or 5 μM Thio-2; cells were treated with vehicle (Ethanol 0.1 %) or 10 nM dihydrotestosterone (DHT) for 16 h thereafter (17 hours total treatment). Following treatments, cells were harvested and lysed, and RNA was extracted using Qiagen RNAeasyPlus RNA extraction kit Mini as per manufacturer’s instruction. RNA quality was analyzed using the Agilent Tapestation RNA ScreenTape. 500 ng of total RNA from each sample was first used in the NEBNext rRNA Depletion Kit followed by the NEBNext Ultra II Directional RNA Library Prep Kit, according to the manufacturer’s instructions. Library quality was confirmed using the Agilent Tapestation High Sensitivity DNA ScreenTape. The libraries were quantified and normalized by qPCR using the KAPA Library Quantification Kit (Roche). Library clustering was performed on a cBot with Illumina HiSeq PE Cluster Kit v3. The libraries were sequenced as paired-end 101 base pair reads on an Illumina HiSeq 2500 with an Illumina HiSeq SBS Kit v3. Base calling and quality scoring were performed using Real-Time Analyses (version 1.18.64) and FASTQ file generation and de-multiplexing using CASAVA. Paired end raw reads in FASTQ format were aligned to the reference human genome (GRCh37/hg19) using RNA sequencing spliced read mapper TopHat (v2.1.0), with default settings. The library and mapping quality were assessed using Picard tools (http://broadinstitute.github.io/picard).

Differential gene expression was calculated using Cuffdiff (Cufflinks v2.2.1), with default settings. The expressed genes (median gene expression level (FPKM) in either control and Thio-2 treated samples > 0; number of genes = 25635) were ranked from high to low using the fold change (log_2_), and subsequently used for pathway analysis. Pathway analysis was performed using the Gene Set Enrichment Analysis (GSEA) Pre-Ranked algorithm from GSEA software (v4.1.0). GSEA Pre-Ranked results were obtained using the H collection of Hallmark gene sets (MsigDB v7.0), with default parameters.

### Chromatin immunoprecipitation (ChIP)-sequencing and analysis

LNCaP cells were grown in starved media (10% charcoal stripped serum) for 72 hours prior to treatment with vehicle (DMSO 0.1 %) or 5 μM Thio-2. Following 1 hour pretreatment with vehicle or 5 μM Thio-2; cells were treated with vehicle (Ethanol 0.1 %) or 10 nM dihydrotestosterone (DHT) for 16 h thereafter (17 hours total treatment). Following treatment, AR ChIP-sequencing (ChIP-seq) was carried out as previously described using the anti-AR antibody (clone N-20; Santa Cruz) (14). ChIP-seq libraries were generated using the ThruPLEX DNA-seq kit (Rubicon Genomics) and were sequenced on the Illumina NextSeq 500 platform at the Molecular Biology Core Facility (Dana-Farber Cancer Institute). All samples were processed through the computational pipeline developed at the Dana-Farber Cancer Institute Center for Functional Cancer Epigenetics (CFCE) using primarily open-source programs (34, 35). Sequence tags were aligned with Burrows-Wheeler Aligner (BWA) to build hg19 and uniquely mapped, non-redundant reads were retained. These reads were used to generate binding sites with Model-Based Analysis of ChIP-Seq 2 (MACS v2.1.1.20160309), with a q-value (FDR) threshold of 0.01 (36, 37). A read per million (RPM) normalized BedGraph signal track file generated by MACS2 is further converted to a BigWig file with bedGraphToBigWig (38). Deeptools is used for the plots heatmap (39).

### canSAR platform

Computational analysis of the druggability of the interfaces for 44 3-dimensional (3D) HSC70-BAG domain structures was carried out using canSAR’s machine learning algorithm. Briefly, the algorithm identifies up to 10 cavities on a 3D-structure and measures ~30 geometric and physicochemical properties for each of these cavities to determine ligandability. The tools and methodologies used are available at our online canSAR platform (40–42). Since proteins are mobile, and this mobility affects the formation of druggable cavities, we performed Monte Carlo simulations to explore limited movements of each structure. The simulations were performed using the CONCOORD method (43). Yamber2, Van der Waals and CONCOORD default bond/angle were set as parameters. A total of 449 alternative structures were shaped (at least 10 structures for each of the 44 original structures) and all cavities identified were assessed for ligandability by canSAR algorithm as described above.

### Cell lines

All cell lines used in this study were grown in recommended media at 37°C in 5% CO_2_ and are detailed in **Supplementary Table 4**. All cell lines were tested for mycoplasma using the VenorGem One Step PCR Kit (Cambio) and STR-profiled using the Cell authentication service by Eurofins Medigenomix.

### Small interfering (si) RNA

Cells were transiently transfected with siRNA as indicated. All siRNA were ON-TARGETplus pools (Dharmacon; Horizon), listed in **Supplementary Table 5**. The siRNA was used along with 0.4% mRNAiMax transfection reagent (ThermoFisher) as per manufacturer’s instructions and incubated with cells as indicated.

### Development of BAG-1 CRISPR knockout 22Rv1 and LNCaP95 cells

22Rv1 and LNCaP95 BAG-1 CRISPR knockout cells were developed following the manufacturers protocol. Briefly, 100000 cells were transfected with BAG-1 sgRNA (6μM; Synthego) and Cas9 2NLS (0.67μM; Synthego) using the 4D Nucleofector System (Lonza Bioscience). After 48 hours, transfection efficiency was assessed in cells transfected with pmaxGFP (0.4μg; Synthego) using fluorescence microscopy, and sgRNA/Cas9 transfected cells were plated in 96-well plates (one cell per well) for clonal expansion. Visual monitoring of single-cell-derived clones was performed daily. The clones that were clearly derived from single cells were screened for BAG-1 protein levels by western blot and selected for study based on BAG-1 protein knockdown efficiency. Details of the BAG-1 sgRNAs used are listed in **Supplementary Table 6**. Cells transfected with Cas9 2NLS complexed with no sgRNA were used as control.

### siRNA and CRISPR in-vitro cell proliferation

Cell proliferation was measured in BAG-1 CRISPR knockout cells or siRNA treated cell in response to vehicle (0.1% DMSO) and Thio-2 (5μM and 50μM) using CellTiter-Glo (Promega) according to the manufacturer’s instructions. Briefly, 3000 cells/well were plated in 96-well plates. For siRNA treated cells, 24 hours after siRNA transfection, cells were seeded and subsequently (24 hours later) treated with either vehicle or Thio-2 in medium. For CRISPR clones, siRNA transfection was omitted. CellTiter-Glo® Cell Viability Assay (Promega) was used to assay growth according to the manufacturer’s instruction on day 0 or after 6 days of treatment and luminescence was measured using Synergy HTX (BioTek).

### Androgen receptor N-terminus and Thio-2 binding

NMR spectra were recorded at 278 K on either a Bruker 800 MHz Avance NEO or a 600 MHz Bruker Avance III spectrometer, equipped with TCl cryoprobes. Intensities and chemical shift perturbations (CSP) were obtained from ^1^H, ^15^N correlation experiments and calculated using the following equation:

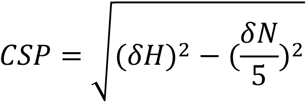

All spectra were referenced using DSS.

NMR spectra were obtained for 25 μM AR-NTD constructs NTD_1-518_ and NTD_330-447_ (Tau-5*) in the present and absence of 250 μM Thio2 and EPI-001. Samples were prepared in phosphate buffer (20 mM sodium phosphate, pH 7.4, 1 mM TCEP, 0.05 % NaN_3_), containing 10 % D_2_O, 10 μM DSS and 0.5 % DMSO-d_6_.

Experiments with ^15^N-labelled AR-NTD constructs NTD_1-518_ and NTD_330-447_ at 25 μM were mixed with a 10 molar excess equivalents (250 μM) of Thio-2 or EPI-001 (positive control) and measured at 5°C.

### Thio-2 solubility

Thio-2 solubility was measured by comparing Thio-2 aromatic signals (region 6.5-8 ppm) to DSS signal (internal reference, at 0 ppm) in 1D ^1^H spectra. Samples containing variable concentrations of Thio-2 were prepared in NMR buffer, containing 10 % D_2_O, 10 μM DSS and 0.5 or 2 % DMSO-d_6_. Samples were measured on 600 MHz spectrometer at 278, 298 and 310 K.

Integration of Thio-2 ^1^H aromatic signals (region 6.5-8 ppm) and the internal reference (DSS) ^1^H signal (at 0 ppm) were used for quantification. Samples containing 5 μM Thio-2, 10 μM DSS, and variable amounts of DMSO-d_6_ (buffer 20 mM sodium phosphate (pH 7.4), 1mM TCEP, 10% D_2_O, 0.05% NaN_3_) were recorded on 600 MHz Bruker Avance spectrometer equipped with a cryoprobe.

### BAG-1 exon 1 and exon 2 knockout mice

Studies with BAG-1 knockout mice were performed at the Karlsruhe Institute of Technology (KIT), Germany according to European and German statutory regulations and approved by the Regierungspräsidium Karlsruhe, Germany. BAG-1 exon 1 deleted knockout mice were kindly provided by Michael Sendtner, Institute for Clinical Neurobiology, University of Wuerzburg, Germany (44). BAG-1 exon 2 deleted knockout mice (Bag1tm1a(EUCOMM)Hmgu) were provided by the Infrafrontier European Mouse Mutant Archive (EMMA). Animals were bred using conventional breeding methods, body weight was measured weekly. At the age of three months mice were culled by heart puncture. Serum was isolated and testosterone content was analyzed by Biocontrol (Bioscientia Healthcare GmbH, Ingelheim, Germany). Subsequently, organs were taken, weighed, and fixed for immunohistochemistry or snap frozen for protein and RNA preparation.

### Micro-array analysis (exon 1 BAG-1 knockout mice)

Prostates from BAG-1 exon 1 deleted heterozygous and wild-type control mice castrated for 12 weeks were minced and subjected to total RNA extraction using TRIzol (Invitrogen) and the RNAeasy Mini purification kit (Qiagen). Biological triplicate RNAs were hybridized to a human U133 Plus 2.0 expression array (Affymetrix) at the Dana-Farber Cancer Institute Microarray Core Facility. Gene expression data were normalized and log-scaled using the RMA algorithm and the RefSeq probe definition (45, 46).

### RNA-sequencing and analysis (BAG-1 knockout mice)

From 1 μg of total RNA we pulled down polyadenylated RNAs with poly-dT magnetic beads. We then prepared sequencing libraries with the TrueSeq Stranded mRNA kit (Illumina) following manufacturer protocol. These libraries were sequenced in paired-end mode (2×50 cycles) with a Hiseq1500 sequencer (Illumina). Raw sequencing data were demultiplexed with Bcl2fastq (version 2.17.1.14, Illumina). Paired end raw reads in FASTQ format were aligned to the reference mouse genome (mm9) using RNA sequencing spliced read mapper TopHat (v2.0.7), with default settings. Differential gene expression and individual gene and transcript expression in units of FPKM (fragments per kilobase of transcript per million mapped reads) were calculated using Cuffdiff (Cufflinks v2.2.1), with default settings. The expressed genes (median expression in either control or treatment samples > 0; n = 17459) were ranked from high to low using the fold change (log_2_), and subsequently used for pathway analysis. Pathway analysis was performed using the GSEA Pre-Ranked algorithm from GSEA software (v4.1.0). GSEA Pre-Ranked results were obtained using the H collection of hallmark gene sets and the C2 collection of curated gene sets (MSigDB v7.1), with default parameters. H and C2 collections were previously mapped to mouse orthologs using the HGNC Comparison of Orthology Prediction tool (https://www.genenames.org/tools/hcop/).

### Development of BAG-1 knockout in PTEN conditional knockout and TRAMP transgenic mouse models

BAG-1 exon 2 deleted knockout mice (Bag1tm1a(EUCOMM)Hmgu) were cross-bred with the inducible PSA-Cre^ERT2^/PTEN^fl/fl^ knockout mouse as well as with the transgenic adenocarcinoma mouse prostate (TRAMP) model of prostate cancer (47, 48). PSA-Cre^ERT2^/PTEN^fl/fl^ mice with and without BAG-1 deletion were treated at 8 weeks of age with a 5-day course of Tamoxifen injection (100μl of 1mg Tamoxifen in ethanol/sesame oil 1:10 mixture, once daily, intraperitoneal). Mice were sacrificed at 14 months of age, the prostate were taken out, weighed, and analysed for tumour stage (Hematoxylin and Eosin stain, H&E), and BAG-1 and AR IHC. TRAMP mice with BAG-1 deletion and WT littermate controls were sacrificed at 6 months and analysed as the PSA-Cre^ERT2^/PTEN^fl/fl^ knockout mice.

### Development of BAG-1L specific transcription activator-like effector nuclease (TALEN) knockout and BAG-1 small hairpin (sh) RNA LNCaP cells

BAG-1 L specific TALEN knockout and TALEN control LNCaP cells were generated as previously described (14). BAG-1 shRNA and control shRNA LNCaP cells were generated using predesigned BAG1 MISSION shRNA lentiviral transduction particles in pLKO.1; clones NM_004323.2-506s1c1 (clone 506) and NM_004323.2-666s1c1 (clone 666) or pLKO.1 non-silencing control (clone control C2) respectively (Mission^®^; Sigma-Aldrich). Briefly, 1000 LNCaP cells were seeded per 96well, allowed to adhere overnight, and transduced with 1 x 10^4^ TU of viral particles. After 48 hours, the medium was exchanged, and positive mass cultures selected using 1 μg/ml puromycin. shRNA mediated knockdown of BAG-1 was confirmed by western blot and qRT-PCR.

### Immunoprecipitation

22Rv1 cells were plated and left for 48 hours. Cells were resuspended in HMKEN buffer (10 mM HEPES, pH 7.2, 5 mM MgCl_2_, 142 mM KCl, 2 mM EGTA, 0.2% (v/v) Nonidet P40, 1:100 protease inhibitor cocktail (Sigma Chemical)) by trituration through a 21-gauge needle, lysed on ice for 30 minutes, and clarified by centrifugation (13,000 rpm for 30 minutes). One-thirtieth (50 μl) of the lysate was retained as a whole-cell lysate. The remaining sample was precleared by use of protein A/G magnetic Dynabeads^TM^ (ThermoFisher) for 30 minutes at 4°C. Dynabeads™ were removed by using a DynaMag-2™ magnet. Lysate (600 μl) was incubated with 5 μg BAG-1L specific antibody rabbit monoclonal antibody (clone RM310; RevMAb Biosciences) at 4 °C for 16 hours to analyze the specificity of this antibody for its target. A further lysate (600 μl) was incubated with 5 μg rabbit immunoglobulins (Vector Laboratories) to control for nonspecific interactions. The immune complexes were incubated with protein A/G magnetic Dynabeads™ for 4 to 6 hours and removed using a magnet. The beads were washed five times by use of HMKEN buffer, resuspended Nupage™ LDS gel electrophoresis sample buffer supplemented with Nupage™ reducing agent (both ThermoFisher), and heated at 95 °C for 5 minutes. Western blotting was performed as described above.

### Institute of Cancer Research/The Royal Marsden Hospital and University of Washington/Fred Hutchinson Cancer Research Center tissue samples

The ICR/RMH patient IHC cohort consisted of forty-three castration-sensitive prostate cancer (CSPC) and sixty-seven castration-resistant prostate cancer (CRPC) tissue biopsies from men with CRPC treated at The RMH. The UW/FHCRC patient IHC cohort consisted of a tissue microarray that included thirty radical prostatectomies and a tissue microarray including thirty metastases from the University of Washington Medical Center Rapid Autopsy Program (49). Human biological samples were sourced ethically, and their research use was in accordance with the terms of the informed consent provided. All tissue blocks were freshly sectioned and were only considered for IHC analyses if adequate material was present.

### Study approvals

All patients treated at The RMH had provided written informed consent and were enrolled in institutional protocols approved by the Royal Marsden NHS Foundation Trust Hospital (London, United Kingdom) ethics review committee (reference 04/Q0801/60). All procedures involving human subjects at the University of Washington (Seattle, Washington, USA) and Fred Hutchinson Cancer Research Center (Seattle, Washington, USA) were approved by the Institutional Review Board at those institutions. All mouse work was carried out in accordance with the Institute of Cancer Research guidelines, including approval by the ICR Animal Welfare and Ethical Review Body, and with the UK Animals (Scientific Procedures) Act 1986, and/or in accordance with the German national and KIT institutional guidelines, including approval by the KIT Animal Welfare and Ethical Review Body, and the Regierungspräsidium Karlsruhe, Germany.

### Statistical analyses

Unpaired Student t-tests were used to determine the difference between growth of PDX-Os, PDXs and prostate cancer cell lines (with siRNA control/BAG-1 and CRISPR control/BAG-1) treated with vehicle or Thio-2. Unpaired Student t-tests were used to determine the difference between mRNA expression of PDX-Os, PDX and prostate cancer cell lines (with siRNA control/BAG-1 and CRISPR control/BAG-1) treated with vehicle or Thio-2. Unpaired Student t-tests were used to determine the difference between weights of non-tumor bearing mice and organs treated with vehicle or Thio-2. Unpaired Student t-tests were used to determine the difference between serum PSA of PDXs treated with and without Thio-2. Kruskal-Wallis test was used to determine the difference between ligandable properties of BAG-1, BCL2 and druggable protein kinase ATP site. Unpaired Student t-tests were used to determine the difference between mRNA expression of BAG-1 exon 2 deleted knockout and wildtype mice. Overall survival of BAG-1 exon 2 deleted knockout and wildtype mice were estimated using the Kaplan–Meier method, and respective hazard ratios were obtained by Cox regression. Unpaired Student t-tests were used to determine differences between characteristics of BAG-1 exon 2 deleted knockout and wildtype mice, including prostate weights following crossing with TRAMP and PTEN knockout models. Chi-squared tests were used to determine the differences between prostate histologies in BAG-1 exon 2 deleted knockout and wildtype mice following crossing with TRAMP and PTEN knockout models. Unpaired Student t-tests were used to determine the difference between BAG-1 isoforms, BAG-1L and AR-FL protein expression in clinical patient cohorts and BAG-1 exon 2 deleted knockout and wildtype mice following crossing with TRAMP and PTEN knockout models. Fisher’s exact tests were used to determine the difference in 12-week PSA response in response to AR-targeted therapy by BAG-1 L protein expression. The time to CRPC and overall survival from diagnosis, and time to PSA progression, radiological/clinical progression and overall survival on AR-targeted therapy, were estimated using the Kaplan-Meier method, and respective hazard ratios were obtained by Cox regression. Statistical analyses were performed with GraphPad Prism Version 7 (GraphPad Software). All experimental replicates and statistical analyses performed are detailed in figure legends. Statistical significance was pre-specified at P ≤ 0.05. No adjustment for multiple testing has been made.

## Results

### Thio-2 inhibits androgen receptor signaling and growth of patient derived models of castration resistant prostate cancer

The tool compound Thio-2 has been postulated to bind the BAG domain of the BAG-1 isoforms and suppress BAG-1L enhanced AR transactivation, however, those studies were primarily performed with AR stimulation (14, 25). We therefore explored the impact of Thio-2 on AR signaling and growth of patient derived models of castration resistant prostate cancer (CRPC). We utilized three of our patient derived xenograft (PDX) models, CP50, CP89 and CP142, all of which were developed from lymph node biopsies of patients with CRPC (**Supplementary Figure 1A**) (28–30). PDX-organoids (PDX-O) were derived from these individual PDX models to support interrogation of Thio-2 *in-vitro*. Having validated a panBAG-1 antibody for immunohistochemistry (IHC), we demonstrated AR-FL and BAG-1 expression across all PDX and their related PDX-O models (**Supplementary Figure 1B and 2**). Thio-2 inhibited the growth of PDX-Os from CP50, CP89 and CP142 in a dose-dependent manner (**Figure 1A, Supplementary Figure 3A-B**). In the CP50 PDX-O model, Thio-2 did not significantly impact AR-FL and BAG-1 protein expression (by western blot and IHC), or consistently down-regulate AR target genes (PSA, TMPRSS2 and FKBP5), although PSA was suppressed at the highest concentration studied (**Figure 1B-D**). We next investigated Thio-2 *in-vivo* and first explored whether any toxicity was associated with Thio-2 by treating non tumor-bearing mice with 15 mg/kg once daily (OD) intraperitoneal (IP), which significantly impacted heart weight (P < 0.01, Student t-test), and although not significant, reduced other parameters including kidney, testes, seminal vesicles, prostate, and body weight (**Supplementary Figure 4A-C**). We next explored the impact of 15 mg/kg OD IP Thio-2 on AR signaling and growth in the tumor bearing CP50 PDX, to determine if any therapeutic impact was observed (**Supplementary Figure 4D**). Thio-2 significantly (P < 0.01, Student t-test) decreased the growth of CP50 PDX compared to vehicle (**Supplementary Figure 4E**). In addition, Thio-2 treatment reduced serum and tumor PSA protein levels, although this was not consistent with tumor PSA mRNA expression (**Supplementary Figure 4F-H**). In addition, other AR target genes (TMPRSS2 and FKBP5) were downregulated (**Supplementary Figure 4G-H**). Taken together, these *in-vitro* and *in-vivo* data demonstrate Thio-2 demonstrating anti-tumor activity and pharmacodynamic modulation of AR signaling in patient derived models of CRPC.

**Figure 1:**
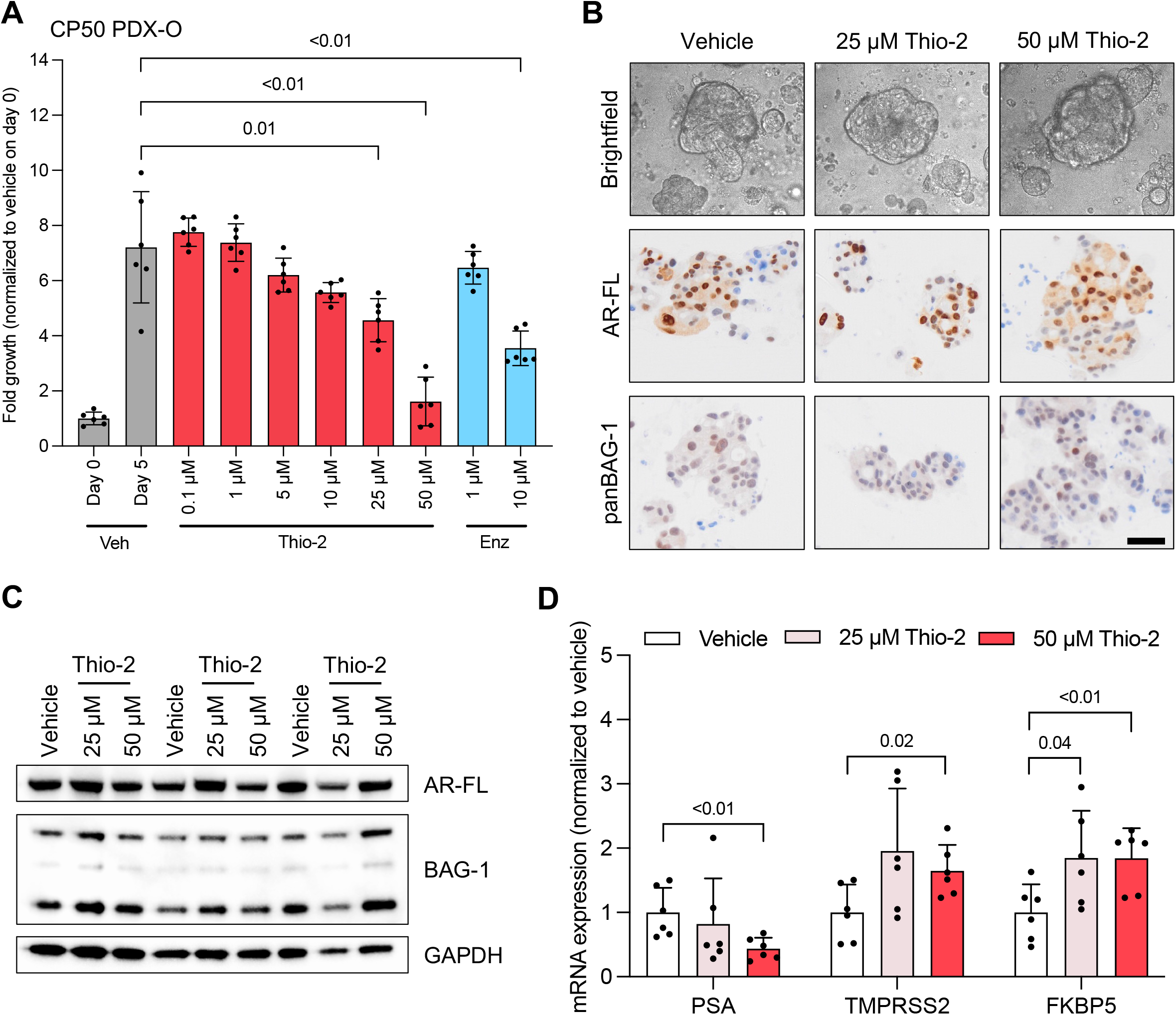
Thio-2 inhibits the growth of castration resistant prostate cancer patient derived xenograft organoids with associated suppression of androgen receptor target genes. **(A)** CP50 patient derived xenograft organoids (PDX-Os) were treated with vehicle (DMSO 0.1 %) or various concentrations (0.1, 1, 5, 10, 25 and 50 μM) of Thio-2 or Enzalutamide (Enz, 1 and 10 μM) and growth determined after 5 days by CellTiter-Glo® 3D Cell Viability Assay. Mean fold change in growth (compared to day 0) with standard deviation from a single experiment with six replicates is shown. P values were calculated for each condition compared to vehicle at 5 days using unpaired Student t-test. P values ≤ 0.05 are shown. **(B)** CP50 PDX-O treated with vehicle (DMSO 0.1 %) or various concentrations (25 and 50 μM) of Thio-2 for 17 hours are shown as live brightfield microscopy images of *in-vitro* cultured PDX-O and representative micrographs of androgen receptor (AR-FL) and pan BAG-1 (panBAG-1) detection by immunohistochemistry of formalin-fixed paraffin-embedded PDX-O are shown. Scale bar: 50 μm. **(C)** CP50 PDX-O were treated with vehicle (DMSO 0.1 %) or various concentrations (25 and 50 μM) of Thio-2 for 17 hours. The effect of each condition on AR-FL, BAG-1 and GAPDH protein expression was determined. Single western blot with triplicates is shown. **(D)** CP50 PDX-O were with vehicle (DMSO 0.1%) or various concentrations (25 and 50 μM) of Thio-2 for 17 hours. The effect of each condition on PSA, TMPRSS2 and FKBP5 mRNA expression was determined. Mean mRNA expression (normalized to average of GAPDH/B2M/HRPT1/RPLP0 and vehicle treatment; defined as 1) with standard deviation from a single experiment with six replicates is shown. P values were calculated for each condition compared to vehicle using unpaired Student t-test. P values ≤ 0.05 are shown.

### Thio-2 inhibits the induction of dihydrotestosterone responsive genes by disrupting binding of the androgen receptor to androgen response elements in LNCaP prostate cancer cells

To further investigate the mechanism of action of Thio-2 and its impact on cellular pathways in prostate cancer, RNA-sequencing (RNA-seq) was performed on LNCaP cells grown in full media treated with 50 μM Thio-2 or vehicle (DMSO 0.1 %) for 17 hours (**Figure 2A**). To investigate pathways associated with the observed gene expression changes, Gene Set Enrichment Analysis (GSEA) was performed using the Hallmarks gene set from the Molecular Signatures Database (MsigDB) (50). Sixteen pathways were found to be significantly enriched after Thio-2 treatment, mainly associated with immune and inflammatory responses (**Figure 2B-C**), but also AR response with the normalized enrichment score (NES) being −2.43, and a false discovery rate (FDR) of < 0.001. Additionally, other key pathways implicated in prostate cancer biology such as MYC targets V1 (NES −2.38, FDR < 0.001), MYC targets V2 (NES −2.14, FDR < 0.001) and DNA repair (NES −1.59, FDR < 0.001) were identified (**Figure 2B-G; Supplementary Table 7**). These data confirm that Thio-2, albeit at relatively high concentrations, inhibits AR signaling and regulates important gene networks implicated in prostate cancer biology. Next, we investigated whether lower concentrations (5 μM) of Thio-2 are also sufficient to inhibit AR signaling and genome-wide AR binding in response to dihydrotestosterone (DHT). To investigate this, RNA-seq was performed using LNCaP cells depleted of hormones for 72 hours and then treated with 5 μM Thio-2 (or DMSO 0.1 %) for 1 hour, prior to stimulation with 10 nM DHT (or ethanol) for 16 hours (**Figure 3A**). The expression of 471 genes significantly (P ≤ 0.05, absolute log_2_ fold change > 1) changed in response to DHT (**Figure 3B-C**). Treatment with 5 μM Thio-2 led to a reduction in gene expression changes, with only 151 (32%) of those 471 DHT-regulated genes remaining altered following DHT treatment (**Figure 3B and D**). Consistent with the relatively small number of genes whose expression changed in response to 5 μM Thio-2, only a small number of Hallmark gene sets were found to be altered (**Figure 3E and F**).

**Figure 2:**
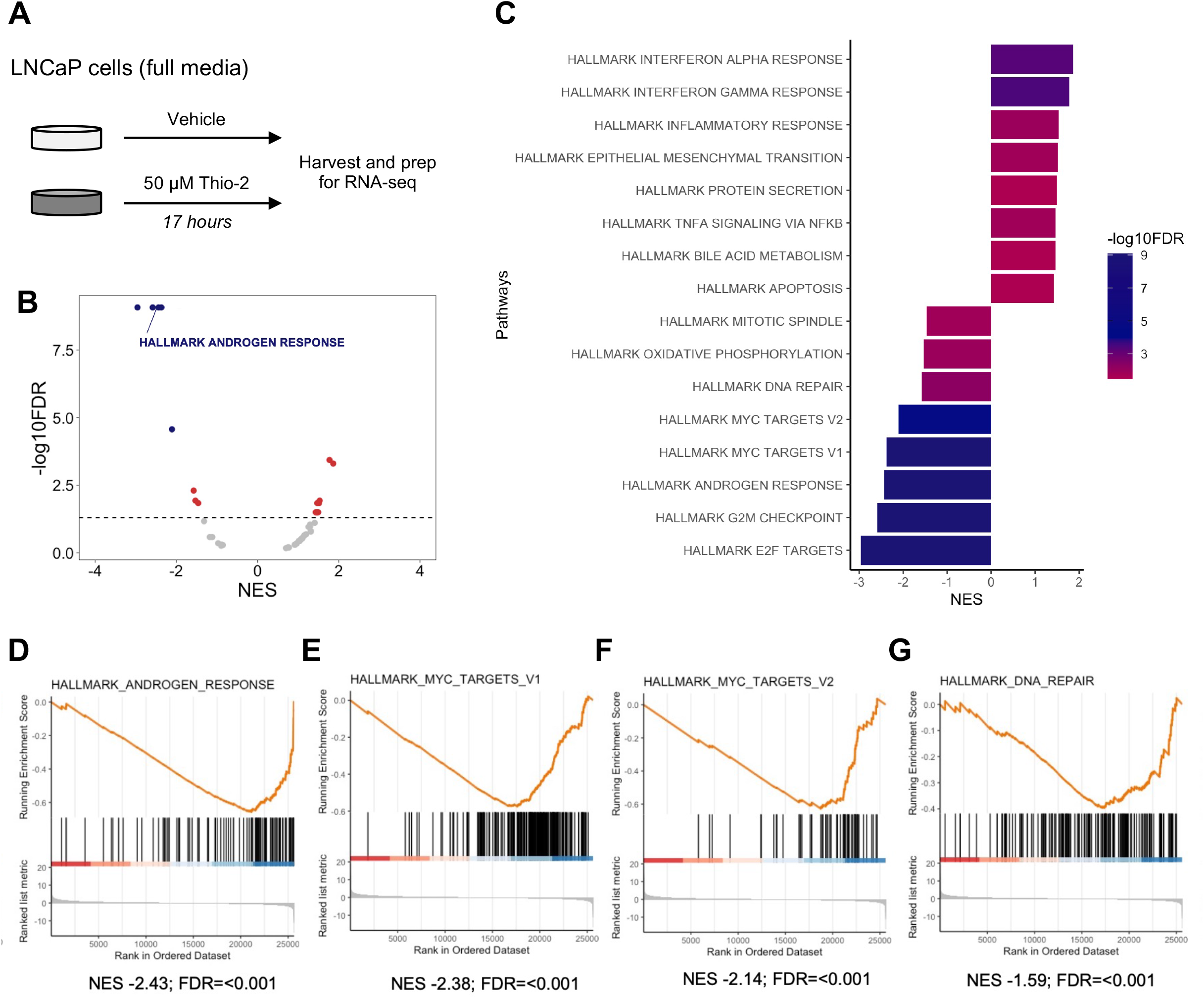
Thio-2 downregulates androgen receptor response and other pathways important in prostate cancer biology in LNCaP prostate cancer cells. **(A)** Schematic of RNA-sequencing (RNA-seq) experimental setup. LNCaP cells were grown in full media (10% fetal bovine serum) prior to treatment with vehicle (DMSO 0.1 %) or 50 μM Thio-2 for 17 hours. RNA-sequencing was performed on a single experiment in triplicate. **(B)** Analysis of RNA-seq with Gene Set Enrichment Analysis (GESA) test shows Thio-2 treatment associates with census Hallmark pathways. Normalized Enrichment Scores (NES) and False Discovery Rates (FDR) are shown. Dotted line indicates significant threshold (FDR 0.05). Colored dots denote significant Hallmark pathways enriched and decreased with Thio-2 treatment. **(C)** Hallmark pathways significantly enriched and de-enriched with Thio-2 treatment are shown. **(D-G)** Leading edge plots for AR response (D), MYC targets V1 (E), MYC targets V2 (F) and DNA repair (G). NES and FDR are indicated below the graphs.

**Figure 3:**
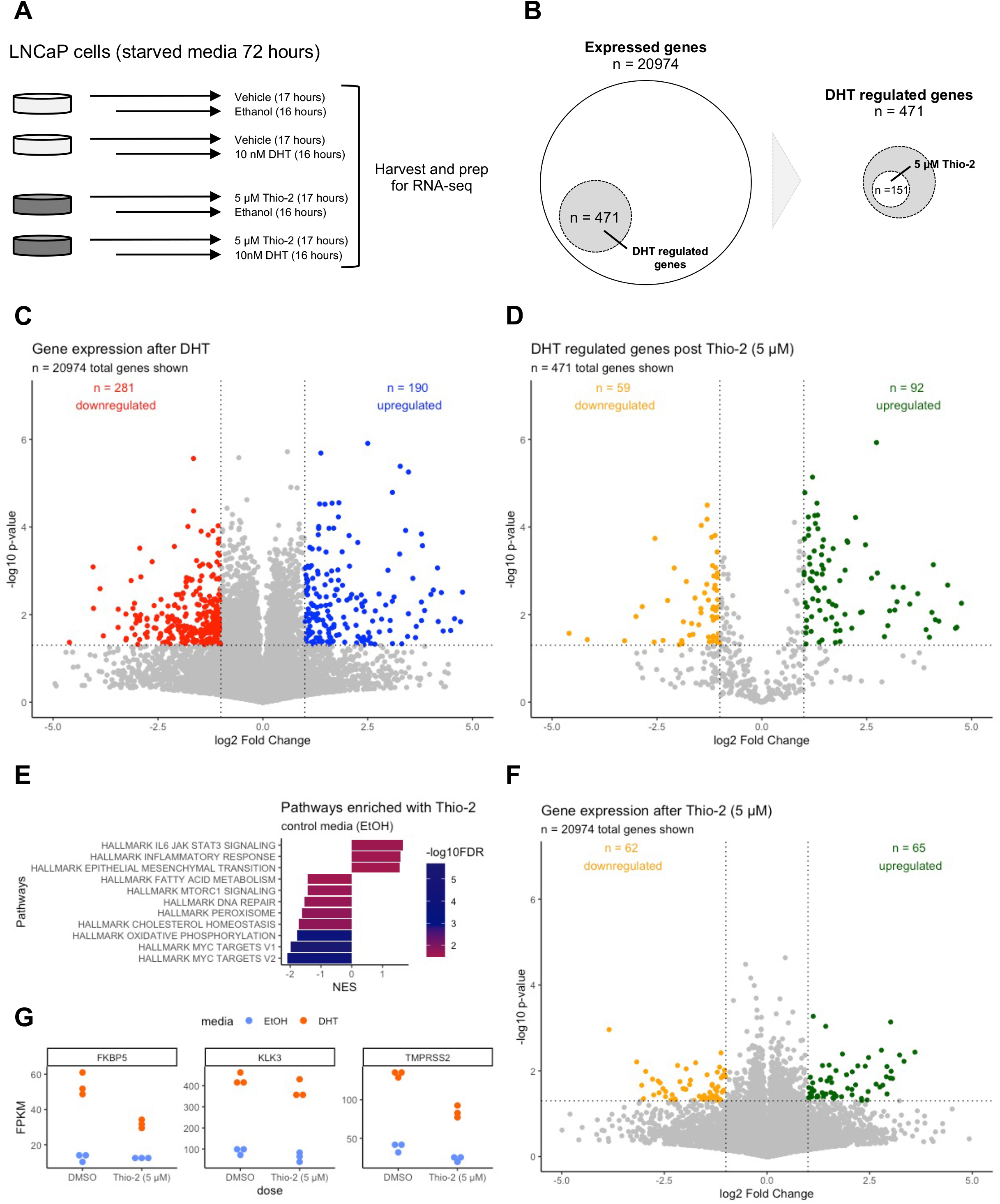
Thio-2 suppresses regulation of androgen receptor responsive genes by dihydrotestosterone in the prostate cancer cell line LNCaP. **(A)** Schematic of RNA-sequencing (RNAseq) experimental setup. LNCaP cells were grown in starved media (10% charcoal stripped serum) for 72 hours prior to treatment with vehicle (DMSO 0.1 %) or 5 μM Thio-2. Following 1 hour pre-treatment with vehicle or 5 μM Thio-2; cells were treated with vehicle (Ethanol 0.1 %) or 10 nM dihydrotestosterone (DHT) for 16 hours (17 hours total treatment). RNAseq was performed on a single experiment in triplicate. **(B-D)** DHT regulated genes (n = 471) were identified by quantifying mRNA expression in starved (vehicle; Ethanol 0.1 %) media and DHT induced media (P value ≤ 0.05, absolute log_2_ fold change > 1) (B, C). Out of 471 DHT regulated genes, 151 remain differentially expressed after Thio-2 treatment at 5 μM (B, D). Venn and volcano plots are shown. Horizontal dotted line indicates the significance threshold (P = 0.05). Vertical dotted line indicates the fold change threshold (absolute log_2_ fold change > 1). **(E-F)** Transcriptome analysis with the GSEA algorithm shows significant enrichment in key Hallmark pathways following 5 μM Thio-2 treatment in starved media (E) and volcano plot showing differentially expressed genes following 5 μM Thio-2 in starved media (P ≤ 0.05, absolute log_2_ fold change > 1) (F). Horizontal dotted line indicates the significance threshold (P ≤ 0.05). Vertical dotted line indicates the fold change threshold (absolute log_2_ fold change > 1). **(G)** Absolute mRNA expression of androgen receptor regulated genes following 5 μM Thio-2 treatment in starved (Ethanol 0.1 %; blue) media and DHT induced (DHT; orange) media is shown.

Furthermore, when focused on individually, commonly studied AR-regulated genes FKBP5, KLK3 and TMPRSS2 demonstrate the same Thio-2 regulation (**Figure 3G**). These data suggest that Thio-2 acts, in part, by preventing expression of AR-regulated genes. To investigate this further, we performed AR chromatin immunoprecipitation under the same conditions as the previously presented RNA-seq to determine the impact on 5 μM Thio-2 on genome-wide AR binding (**Figure 2A**). Thio-2 moderately reduced genome-wide binding of both the unstimulated and stimulated AR, suggesting that Thio-2 may, in part, reduce AR signaling through destabilization of AR at its DNA binding sites (**Supplementary Figure 5**).

### Druggability analyses of the BAG domain demonstrate that the BAG-1 isoforms represent a challenging drug target

We had previously reported that BAG-1 presents a large groove, suitable for peptide or peptide-mimetic modulators, but inconsistent with a small molecule inhibitor. Nonetheless, we wanted to interrogate whether Thio-2 elicits its action through a BAG-1 isoform-mediated mechanism as it has been suggested it may bind the BAG domain of the BAG-1 isoforms through *in-silico* docking experiments (14, 25). Using our updated canSAR analysis, the 44 3-dimensional (3D) HSC70-BAG domain structures continue to reveal a lack of a classical ‘ligandable’ cavity within the BAG domain (**Supplementary Table 8**). This updated analysis, continues to show the same large groove that would be inconsistent with a small molecule, but rather more suited to a peptide inhibitor. (**Figure 4A**) (14, 40, 41, 51, 52). Our machine learning approaches assess all identified cavities for key geometric and physicochemical parameters which contribute to the druggability of a pocket. These include a measure of the enclosure of the entire cavity (in this case groove); the volume of the most enclosed pocket within this cavity; the ratio of non-polar to polar chemical groups within the pocket; the inverse Andrew’s Energy; and the number of accessible and buried vertices, as well as the ration of hydrophobic atoms within the pocket. We compared all these parameters to a typical ‘druggable pocket’ parameters such as would be observed for ATP binding sites of protein kinases and protein-protein interactions (such as BCL2). We find that the properties of the BAG-1 groove fall outside the distributions expected for druggable cavities (all P < 0.01, Kruskal-Wallis test) (**Figure 4B**) (14, 42, 53). We next wanted to explore whether a cryptic druggable cavity might emerge should we probe the structural fluctuations of the protein. To this end, we performed Monte Carlo simulations (43). These stimulations generated 449 models with 4489 cavities (up to 10 cavities per model) from the original 44 3D HSC70-BAG domain structures (**Figure 4C**). Interestingly, despite allowance for structural fluctuations, the cavity of interest (80% of amino acid residues within the original pocket; 3FZLB) within all these models remains challenging for small molecule inhibition (**Figure 4C**). Furthermore, of the remaining cavities identified, only seven (of 4489) cavities have been identified as ligandable, and these may represent artifact as they are only identified in a limited number of models (six of 449) derived (**Figure 4C**). These data identify the BAG domain of the BAG-1 isoforms to be a challenging drug target for small molecule inhibition that may require alternative approaches such as peptidomimetics suggesting that Thio-2 may not function through a BAG-1 mediated mechanism.

**Figure 4:**
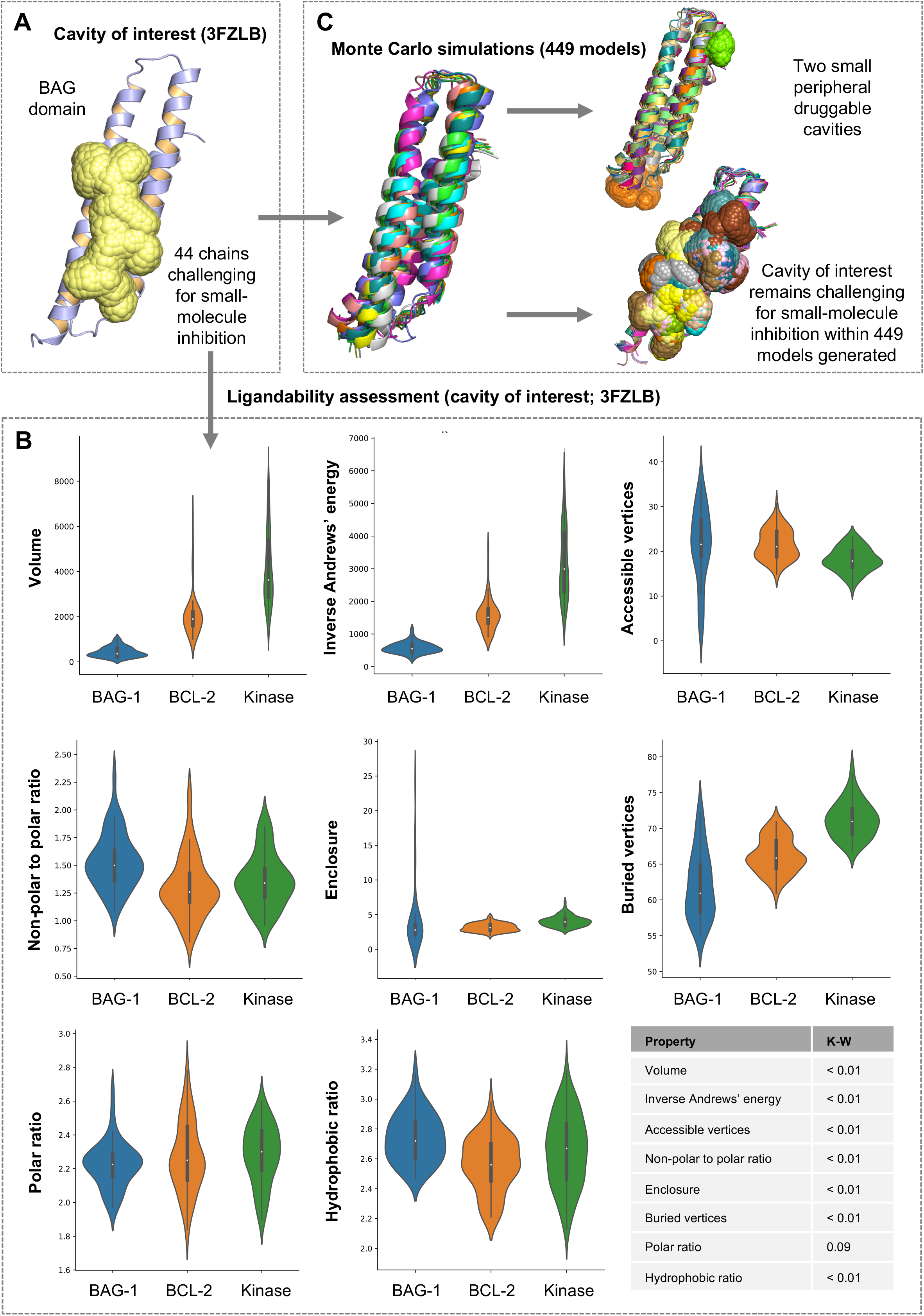
Druggability assessment of the BAG domain of BAG-1. **(A)** Visualization of the BAG domain cavity of interest identified by canSAR using the 44 3-dimensional (3D) HSC70-BAG domain structures available mapped onto the representative structure (PDB ID 3FZLB). The cavity of interest as volume surface (in yellow) is shown on the BAG domain (violet) of BAG-1. **(B)** Key geometric and physicochemical parameters for the cavity of interest within the BAG domain (blue), a druggable protein-protein interaction (BCL-2; orange) and the druggable kinase ATP site (green) are shown as violin plots. P values were calculated using the Kruskal-Wallis (K-W) test. **(C)** Monte Carlo simulations identified 449 models with 4489 cavities for the original 44 3D HSC70-BAG domain structures. The cavity of interest remains challenging for small molecule inhibition within all models despite allowances for structural fluctuations.

### Thio-2 inhibits growth and androgen receptor signaling in prostate cancer cell lines through a BAG-1 isoform independent mechanism

We next interrogated whether Thio-2 functions through a BAG-1 isoform-dependent mechanism, and whether abrogating the function of all BAG-1 isoforms is an attractive therapeutic strategy in CRPC. To support these studies, we utilized siRNA knockdown and CRISPR knockout of all three BAG-1 isoforms (S, M and L) in prostate cancer cell line models. Interestingly, BAG-1 isoform siRNA knockdown led to a small but significant increase in growth of LNCaP (P = 0.02, Student t-test) and 22Rv1 (P < 0.01, Student t-test), but not in LNCaP95 cells (**Figure 5A and D, Supplementary Figure 6A**). In addition, BAG-1 isoform siRNA knockdown had no effect on AR-FL, AR-V7 or PSA protein expression in all three cell lines (**Figure 5B and E, Supplementary Figure 6B**). Furthermore, BAG-1 isoform siRNA knockdown did not consistently suppress downstream AR target genes (PSA, TMPRSS2 and FKBP5), with significant downregulation of only TMRPSS2 in LNCaP (P < 0.01, Student t-test) and LNCaP95 (P < 0.01, Student t-test) cells, and PSA (P < 0.01, Student t-test), TMPRSS2 (P < 0.01, Student t-test) and FKBP5 (P = 0.02, Student t-test) in 22Rv1 cells respectively (**Figure 5C and F, Supplementary Figure 6C**). However, Thio-2 significantly inhibited the growth of LNCaP, 22Rv1 and LNCaP95 in a dose-dependent manner, irrespective of BAG-1 isoform siRNA knockdown status, suggesting that its growth inhibitory effects may not be mediated entirely through BAG-1 isoforms (**Figure 5A and D, Supplementary Figure 6A**). Furthermore, and in contrast to BAG-1 knockdown, higher concentrations (50 μM) of Thio-2 slightly decreased AR-FL, AR-V7 and PSA protein levels in all three cell lines independent of BAG-1 isoform siRNA knockdown status (**Figure 5B and E, Supplementary Figure 6B**). Moreover, unlike BAG-1 isoform siRNA knockdown alone, Thio-2 seemed to suppress all AR target genes more consistently across all cell lines tested independent of BAG-1 isoform siRNA knockdown status (**Figure 5C and F, Supplementary Figure 6C**). Taken together, these data suggest that BAG-1 isoform function may have little impact on growth and AR signaling in CRPC models, and that Thio-2 inhibition of growth and AR signaling may, in part, be independent of BAG-1 isoform function.

**Figure 5:**
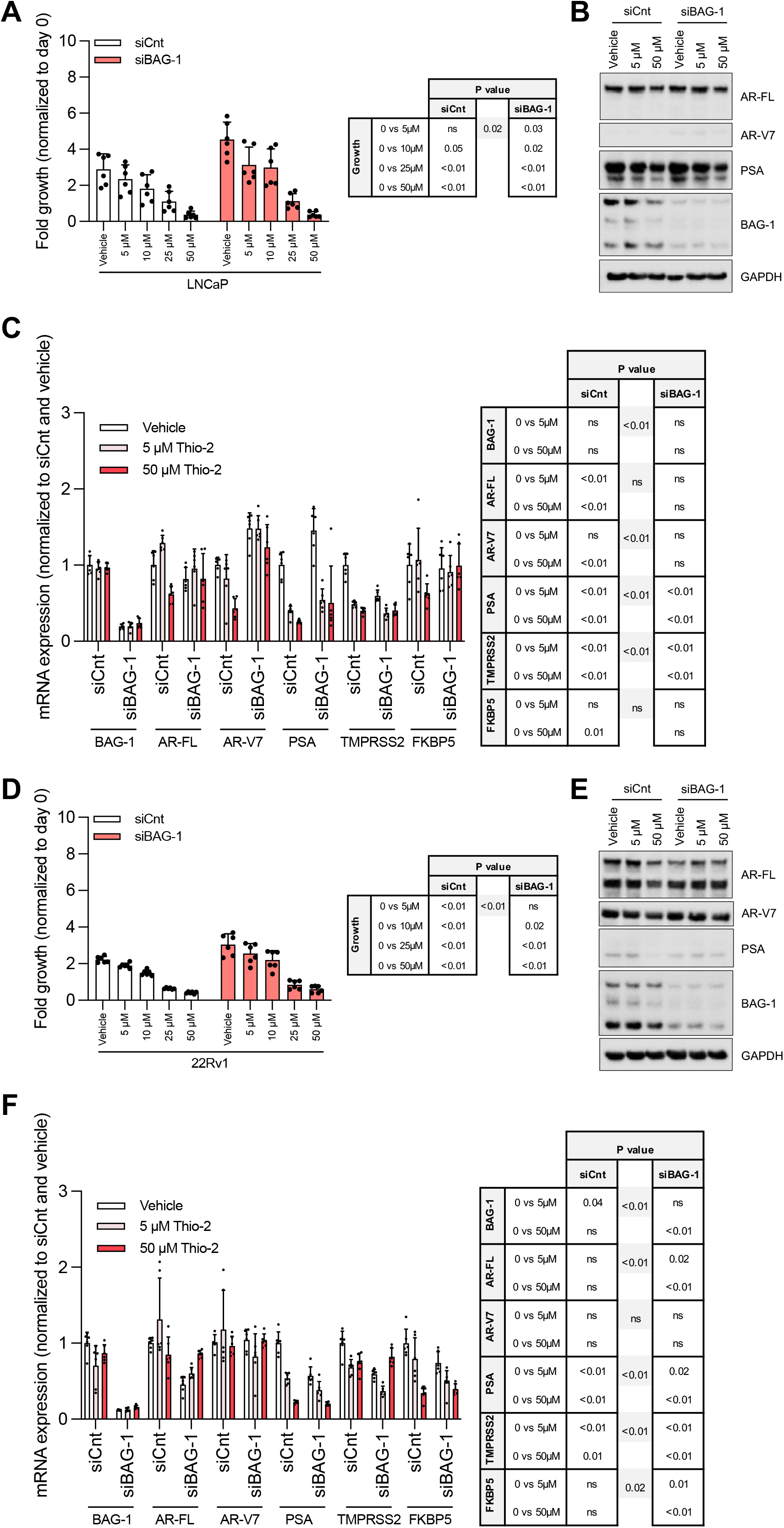
Inhibition of growth and androgen receptor signaling by Thio-2 in the prostate cancer cell lines LNCaP and 22Rv1 is not exclusively dependent on BAG-1. **(A and D)** LNCaP (A) and 22Rv1 (D) prostate cancer cells were transfected with 50nM of either control (siCnt; clear bars) or BAG-1 (siBAG-1; red bars) siRNA for 72 hours prior to treatment with vehicle (DMSO 0.1 %) or various concentrations (5, 10, 25 and 50 μM) of Thio-2 and growth was determined after 6 days by CellTiter-Glo® Luminescent Cell Viability Assay. Mean fold change in growth (compared to day 0) with standard deviation from a single experiment with six replicates is shown. P values were calculated for each condition compared to vehicle in siCnt and siBAG-1 cells, and between vehicle treated siCnt and siBAG-1 cells (grey shading), using unpaired Student t-test. P values ≤ 0.05 are shown and P values > 0.05 are shown as nonsignificant (ns). **(B and E)** LNCaP (B) and 22Rv1 (E) prostate cancer cells were transfected with 50 nM of either siCnt or siBAG-1 siRNA for 55 hours prior to treatment with vehicle (DMSO 0.1 %) or various concentrations (5 and 50 μM) of Thio-2 for 17 hours (total 72 hours) and AR-FL, AR-V7, PSA, BAG-1 and GAPDH protein expression was determined. Single western blot is shown. **(C and F)** LNCaP (C) and 22Rv1 (F) prostate cancer cells were transfected with 50 nM of either siCnt or siBAG-1 siRNA for 55 hours prior to treatment with vehicle (DMSO 0.1 %) or various concentrations (5 and 50 μM) of Thio-2 for 17 hours (total 72 hours) and BAG-1, AR-FL, AR-V7, PSA, TMPRSS2 and FKBP5 mRNA expression was determined. Mean mRNA expression (normalized to average of GAPDH/B2M/HRPT1/RPLP0 and siCnt/vehicle; defined as 1) with standard deviation from a single experiment with six replicates is shown. P values were calculated for each condition compared to vehicle in siCnt and siBAG-1 cells, and between vehicle treated siCnt and siBAG-1 cells (grey shading), using unpaired Student t-test. P values ≤ 0.05 are shown and P values > 0.05 are shown as non-significant (ns).

To further validate, and interrogate these findings, we next developed 22Rv1 and LNCaP95 BAG-1 isoform CRISPR knockout clones exhibiting loss of all three BAG-1 isoforms. In agreement with earlier findings, BAG-1 isoform CRISPR knockout using three different guides in 22Rv1 again led to a significant (all P < 0.01, Student t-test) increase in growth (**Figure 6A**). In contrast to BAG-1 isoform siRNA knockdown which had no effect, BAG-1 isoform CRISPR knockout using multiple guides in LNCaP95 led to a significant (P = 0.04 and 0.03, Student t-test) decrease in growth (**Supplementary Figure 7A**). Similar to BAG-1 isoform siRNA knockdown, BAG-1 isoform CRISPR knockout did not consistently impact AR-FL or AR-V7 protein levels in these cell lines (**Figure 6B, Supplementary Figure 7B**). In addition, BAG-1 isoform CRISPR knockout did not significantly suppress any AR target genes, with several of them significantly increasing in response to BAG-1 knockout (**Figure 6C, Supplementary Figure 7C**). Thio-2 treatment in BAG-1 isoform CRISPR knockout clones significantly inhibited the growth of 22Rv1 and LNCaP95, irrespective of BAG-1 isoform CRISPR knockout status (**Figure 7A, Supplementary Figure 7A**). Higher concentrations (50 μM) of Thio-2 decreased AR-FL and AR-V7 protein levels, although not consistently, in both cell lines independent of BAG-1 isoform CRISPR knockout status (**Figure 6B, Supplementary Figure 7B**). Finally, consistent with BAG-1 isoform siRNA knockdown studies, Thio-2 seemed to suppress AR target genes more consistently across both cell lines independent of BAG-1 isoform CRISPR knockout status (**Figure 6C, Supplementary Figure 7C**). Taken together, these studies confirm that BAG-1 isoform function is not critical for persistent AR signaling and growth in CRPC, and that the AR inhibitory and growth reduction effects of Thio-2 may be mediated through a BAG-1 isoform independent mechanism.

**Figure 6:**
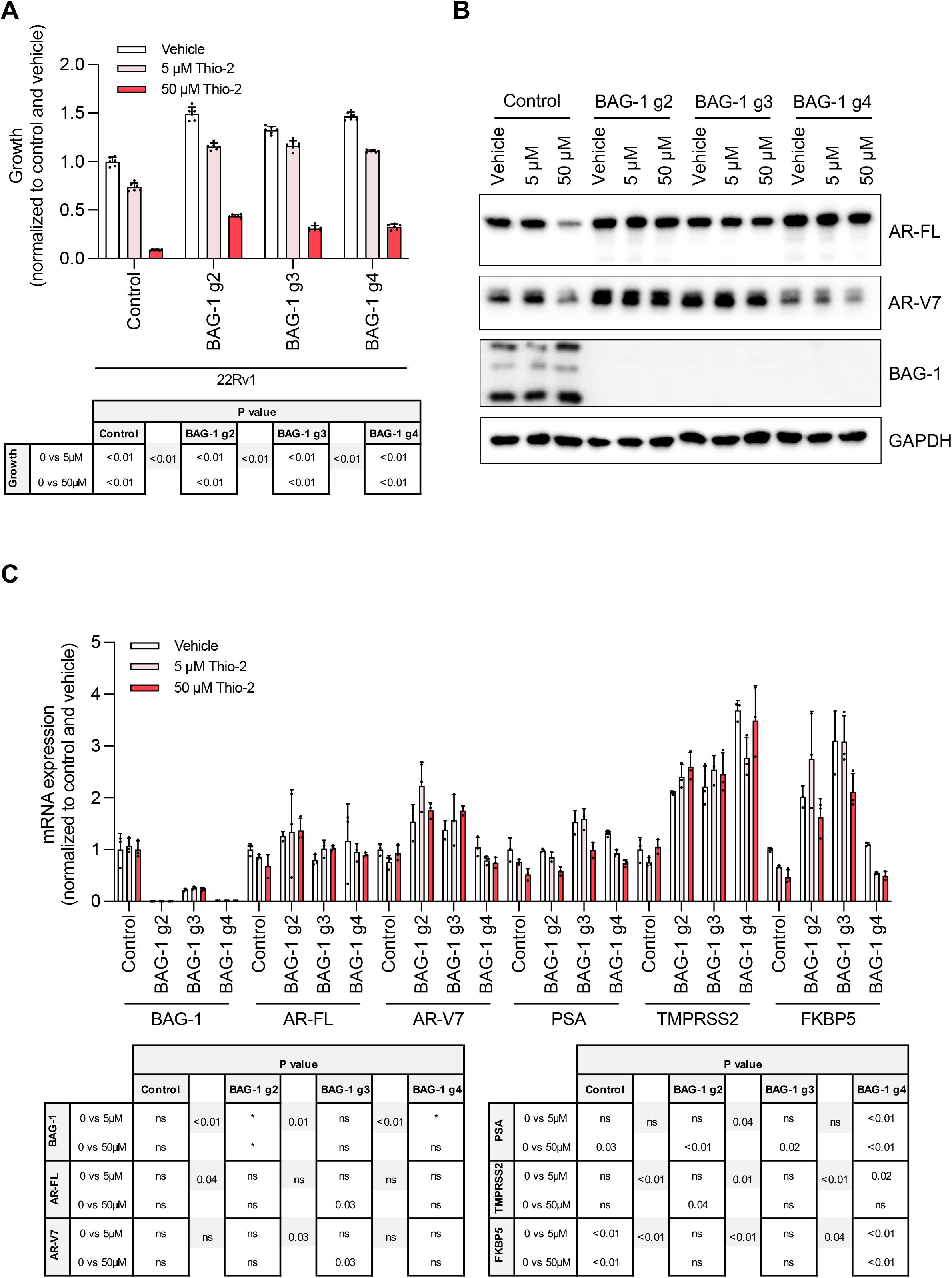
Inhibition of growth and androgen receptor signaling by Thio-2 in the prostate cancer cell line 22Rv1 is not exclusively dependent on BAG-1. **(A)** BAG-1 knockout CRISPR/Cas9 clones were developed in 22Rv1 prostate cancer cells. Control (Cas9) and three BAG-1 knockout (guide 2, g2; guide 3, g3; guide 4, g4) were used for transfection and single cell derived 22Rv1 clones were selected. Clones were treated with vehicle (DMSO 0.1 %) or various concentrations (5 and 50 μM) of Thio-2 and growth was determined after 6 days by CellTiter-Glo® Luminescent Cell Viability Assay. Mean growth (normalized to vehicle treated control clone; defined as 1) with standard deviation from a single experiment with six replicates is shown. P values were calculated for each condition compared to vehicle for each individual guide, and between vehicle treated control and BAG-1 guides (grey shading), using unpaired Student t-test. P values ≤ 0.05 are shown and P values > 0.05 are shown as non-significant (ns). **(B)** 22Rv1 clones were treated with vehicle (DMSO 0.1 %) or various concentrations (5 and 50 μM) of Thio-2 for 17 hours and AR-FL, AR-V7, BAG-1 and GAPDH protein expression was determined. Single western blot is shown. **(C)** 22Rv1 clones were treated with vehicle (DMSO 0.1 %) or various concentrations (5 and 50 μM) of Thio-2 for 17 hours and BAG-1, AR-FL, AR-V7, PSA, TMPRSS2 and FKBP5 mRNA expression was determined. Mean mRNA expression (normalized to average of GAPDH/B2M/HRPT1/RPLP0 and vehicle treated control clone; defined as 1) with standard deviation from a single experiment with three replicates is shown. P values were calculated for each condition compared to vehicle for each individual guide, and between vehicle treated control and BAG-1 guides (grey shading), using unpaired Student t-test. P values ≤ 0.05 are shown and P values > 0.05 are shown as non-significant (ns). *non-positive variance, P values not defined.

**Figure 7:**
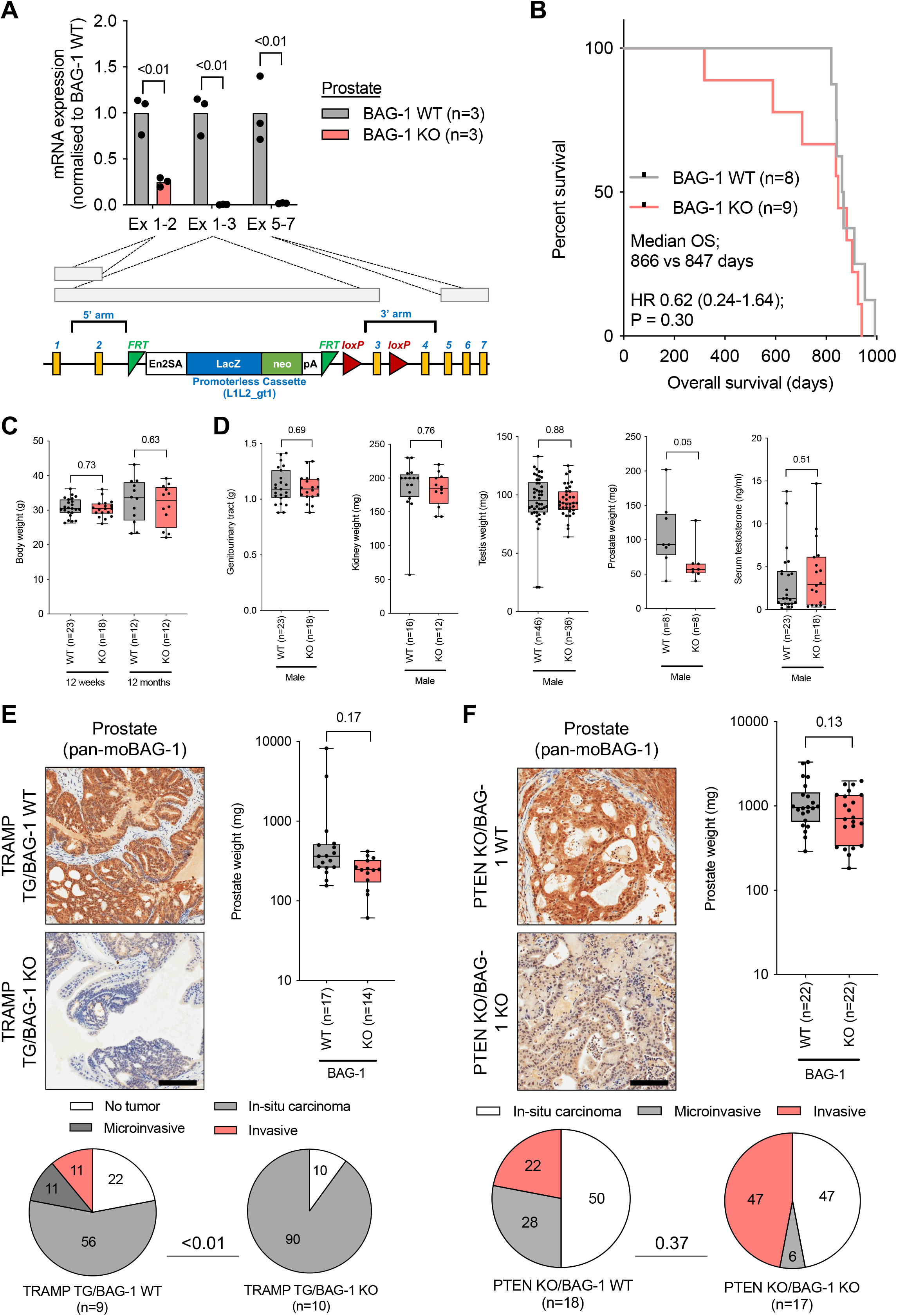
BAG-1 knockout male mice are viable and BAG-1 knockout has limited impact on prostate cancer formation in PTEN conditional knockout and TRAMP transgenic mouse models. **(A)** BAG-1 knockout KO (BAG-1 KO) mouse strain Bag1tm1a(EUCOMM)Hmgu was developed by the European conditional mouse mutagenesis (EUCOMM) program by insertion of an artificial exon containing the coding sequence of beta-Geo, a fusion protein of beta-Galactosidase (LacZ) and neomycin (neo) followed by a stop codon and polyadenylation (pA) sequence between exon 2 and exon 3 flanked by Flp recognition sites disrupts the expression of the WT gene, replacing the endogenous BAG-1 expression by a fusion protein of the BAG-1 N-terminal sequence encoded in exon 1 and 2 and beta-Geo. Mouse prostates from BAG-1 knockout KO (BAG-1 KO) and BAG-1 wildtype (BAG-1 WT) male mice were analyzed for BAG-1 mRNA (quantitative reverse transcriptase-polymerase chain reaction; qRT-PCR) levels. BAG-1 exon 1 to 2 (Ex 1-2), exon 1 to 3 (Ex 1-3) and exon 5 to 7 (Ex 5-7) mRNA was quantified for BAG-1 KO (red symbols; n=3) and BAG-1 WT (gray symbols; n=3) mice. mRNA expression was calculated relative to mouse GAPDH and normalized to BAG-1 WT. Mean levels from three prostates are shown. P values were calculated for BAG-1 KO compared with BAG-1 WT mice using unpaired Student t-test. **(B)** Kaplan-Meier curves of overall survival (OS) of BAG-1 KO (red line; n=9) and BAG-1 WT (gray line; n=8) male mice from birth. Median OS, hazard ratio (HR) with 95% confidence intervals and P values for univariate Cox survival model are shown. **(C)** The body weight of male BAG-1 KO (red bars) and BAG-1 WT (gray bars) male mice at 12 weeks and 12 months was determined. Median body weight with interquartile range, and smallest and largest value, is shown. P values were calculated for BAG-1 KO compared with BAG-1 WT mice using unpaired Student t-test. **(D)** The weight of the genitourinary tract, kidney, testis, prostate, and serum levels of testosterone, from male BAG-1 KO (red bar) and BAG-1 WT (gray bar) male mice at age 3 months and older was determined. Median weight or serum testosterone levels with interquartile range, and smallest and largest value, is shown. P values were calculated for BAG-1 KO compared with BAG-1 WT mice using unpaired Student t-test. **(E)** BAG-1 KO and BAG-1 WT mice were crossed with TRAMP transgenic (TG) mice. Prostates from TRAMP TG/BAG-1 KO and TRAMP TG/BAG-1 WT male mice were analyzed for mouse (mo) BAG-1 protein (immunohistochemistry; IHC) levels. Representative micrographs of BAG-1 detection in mouse prostates by pan-BAG-1 antibody IHC are shown. Scale bar, 100 μm. The weight of the prostates from TRAMP TG/BAG-1 KO (red bar) and TRAMP TG/BAG-1 WT (gray bar) mice at 6 months of age was determined. Median prostate weight (on the log10 scale) with interquartile range, and smallest and largest value, is shown. P value was calculated for TRAMP TG/BAG-1 KO compared with TRAMP TG/BAG-1 WT mice using unpaired Student t-test. Histology review of prostates from TRAMP TG/BAG-1 KO and TRAMP TG/BAG-1 WT mice with P value for chi-squared test for trend is shown. **(F)** BAG-1 KO and BAG-1 WT mice were crossed with inducible PTEN KO mice. PTEN KO was induced by tamoxifen injection at 8 weeks, prostates from PTEN KO/BAG-1 KO and PTEN KO/BAG-1 WT male mice were analyzed for moBAG-1 protein (IHC) levels. Representative micrographs of BAG-1 detection in mouse prostates by pan-BAG-1 antibody IHC are shown. Scale bar, 100 μm. The weight of the prostates from PTEN KO/BAG-1 KO (red bars) and PTEN KO/BAG-1 WT (gray bars) mice at 14 months of age was determined. Median prostate weight (on the log10 scale) with interquartile range, and smallest and largest value, is shown. P value was calculated for PTEN KO/BAG-1 KO compared with PTEN KO/BAG-1 WT mice using unpaired Student t-test. Histology review of prostates from PTEN KO/BAG-1 KO and PTEN KO/BAG-1 WT mice with P value for Chi-squared test for trend is shown.

### Thio-2 binds the AR N-terminus, but demonstrates low solubility which may limit its therapeutic developability

Having demonstrated that Thio-2 inhibits AR signaling in CRPC models independent of BAG-1 isoforms, we further investigated whether this maybe through direct interaction with the AR N-terminus. Previous studies have demonstrated that Thio-2 inhibits AR N-terminal transactivation independent of the longer BAG-1 isoform, BAG-1L (14). Similarly, small molecules such as EPI-001, have been reported to bind the AR N-terminal disordered region (54, 55). This binding can be identified by intensity changes in the NMR protein ^1^H-^15^N correlation spectra for the full-length AR N-terminus (residues 1-558) and chemical shift perturbations in the partially helical regions of a shorter transactivation unit 5 construct (residues 330-447) (**Supplementary Figure 8A-B**). Thio-2 experiments showed modest changes in the same AR regions, suggesting that it may bind the AR through similar binding mechanism (**Supplementary Figure 8C-D**). Despite these intriguing findings, interrogation of Thio-2 solubility estimated it to be 2.5 μM (in 0.5 % DMSO at 37 °C), 1 μM (in 0.5 % DMSO at 25 °C) and 4 μM (in 2% DMSO at 37 °C), suggesting that above low micromolar concentrations the biological phenotypes observed may be related to limitations in solubility (**Supplementary Figure 9**). The solubility limitations of Thio-2 mean that this tentative binding to the AR N-terminus, and the biological phenotype observed at higher concentrations, need to be interpreted with caution.

### BAG-1 does not play a critical role in normal mouse physiological development

To further investigate BAG-1 as a potential drug target, we next moved on to study the impact of BAG-1 loss in BAG-1-deleted mouse knockout models. Targeted deletion of exon 1 and 2 of the BAG-1 gene has shown BAG-1 homozygous deletion to be embryonically lethal (44). Analyzing BAG-1 heterozygous mice that are viable, we demonstrated that BAG-1 mRNA is indeed reduced but surprisingly CHMP5 (which is located on the opposite strand of chromosome 9 to BAG-1) is also downregulated in this model (**Supplementary Figure 10A**). Importantly, CHMP5 deletion has been previously shown to be embryonically lethal; we therefore hypothesized that this apparent double gene (BAG-1/CHMP5) knockout may explain the embryonically lethal phenotype previously reported (56). Considering this, to explore the potential toxicity associated with therapeutic targeting of BAG-1, and the potential impact that CHMP5 co-deletion has on the BAG-1 deletion phenotype, we explored an alternative knockout strategy. We utilized a BAG-1 specific knockout-first mouse strain Bag1tm1a(EUCOMM)Hmgu (referred to as BAG-1 knockout from here on out), developed by the European Conditional Mouse Mutagenesis (EUCOMM) Program to study the impact of losing just BAG-1 (**Figure 7A**). In these mice, the insertion of an artificial exon containing the coding sequence of beta-Geo, a fusion protein of LacZ and neomycin, followed by a stop codon and polyadenylation sequence between exon 2 and exon 3 disrupts the expression of the BAG-1 gene. The cassette is flanked by Flp recognition sites thus providing the option to convert the allele to an inducible knockout cassette, an option that we did not use in this project. The insertion replaces the BAG-1 protein by a fusion protein of the BAG-1 N-terminal sequence encoded by exon 1 and 2 and beta-Geo (**Figure 7A**).

As these BAG-1 knockout (KO) mice are viable and fertile, BAG-1 deletion was confirmed at the mRNA and protein level in prostates isolated from knockout mice and littermate controls (**Supplementary Figure 10B-D**, **Figure 7A**). In order to investigate the impact of Bag-1 deletion on gene expression and signaling pathways, RNA sequencing was performed on BAG-1 KO and wild type (WT) mouse prostates demonstrating significant (P < 0.01, Student t-test) reduction in BAG-1 mRNA with no significant change in CHMP5 or other BAG family members, confirming the knockout is BAG-1 specific and there was no compensatory upregulation of other known BAG family members (**Supplementary Figure 10E**). Despite BAG-1 being a multifunctional protein, there was no significant enrichment in functional pathways in BAG-1 KO compared to WT mouse prostates (**Supplementary Figure 10F**). Consistent with this and in contrast to the embryonic lethal phenotype associated with CHMP5 single and CHMP5/BAG-1 double knockouts, BAG-1 deletion did not impact overall survival (**Figure 7B**, **Supplementary Figure 11A**). However, although most characteristics were not significantly different with BAG-1 deletion, significant differences observed included decreased prostatic weight (P = 0.05, Student t-test), increased duration of pregnancy (P = 0.01, Student t-test), decreased litter size (P = 0.04, Student t-test), increased day 1 neonatal weight (P = 0.02, Student t-test), and decreased neonate survival rate on day 2 (P = 0.04, Student t-test) when comparing BAG-1 KO and WT mice and their litter mates (**Figure 7C-D**, **Supplementary Figure 11B-D**). Consistent with these physiological findings, histological analysis of all major organs demonstrated no difference between BAG-1 KO and WT mice, and there was decreased BAG-1 protein expression in BAG-1 KO mice (**Supplementary Figure 12 and 13**). These data, in contrast to previous studies, suggest that BAG-1 KO mice demonstrate normal physiological development with no severe impact on survival or AR signaling in the mouse prostate.

### BAG-1 knockout has limited impact on prostate cancer formation in PTEN conditional knockout and TRAMP transgenic mouse models

Having demonstrated this non-lethal phenotype from BAG-1 only deletion, we next investigated whether BAG-1 isoforms play a critical role in prostate cancer growth, and crossed BAG-1 KO mice with the TRAMP transgenic and inducible PSA-Cre^ERT2^/PTEN^fl/fl^ prostate tumor models (57). In TRAMP transgenic mice, BAG-1 KO did not significantly impact prostatic weight (**Figure 7E**). In addition, we interrogate the prostate histology which demonstrated BAG-1 KO to be associated with significantly (P < 0.01, Chi-squared test) less invasive cancer (**Figure 7E**). Furthermore, following validation of a mouse AR antibody, we demonstrated there were no significant changes in AR protein expression or localization associated with BAG-1 KO (**Supplementary Figure 14, Supplementary Figure 15A-C**). Next, we crossed these BAG-1 KO mice with the inducible PTEN KO prostate tumor model PSA-Cre^ERT2^/PTEN^fl/fl^. In this model, BAG-1 KO did not significantly impact prostatic weight (**Figure 7F**). In addition, histopathology review demonstrated that BAG-1 KO was not associated with more invasive cancer formation (**Figure 7F**). Furthermore, there was no significant changes in AR protein expression or localization associated with BAG-1 KO (**Supplementary Figure 16A-C**). These data suggest that BAG-1 isoforms do not play a key role in prostate cancer growth in these specific mouse models.

### BAG-1L protein expression does not predict clinical course of advanced prostate cancer

We next investigated the clinical significance of the nuclear isoform BAG-1L, which has been reported to bind and activate the AR and is therefore the most likely modulator of AR within the BAG-1 family of proteins. To further support these studies, a recombinant rabbit monoclonal antibody (clone RM310) against the unique N-terminus of BAG-1L was developed. Antibody validation was performed by western blot analysis of VCaP and 22Rv1 cells, and demonstrating a strong BAG-1L band at 50 kDa that disappeared in our previously described transcription activator-like effector nuclease (TALEN) BAG-1L knockout LNCaP cells, being also significantly reduced in our BAG-1 shRNA treated LNCaP cells and BAG-1 siRNA treated HeLa cells (**Supplementary Figure 17A**) (14). In addition, and unlike panBAG-1 antibodies, RM310 was specific for BAG-1L and did not recognize either BAG-1 or BAG-1M (**Supplementary Figure 17A**). Consistent with this, specificity of RM310 for BAG-1L was confirmed by immunoprecipitation using 22Rv1 cells which demonstrated a single band at 50 kDa (**Supplementary Figure 17B**). Following confirmation RM310 specifically recognizes BAG-1L, we optimized RM310 for IHC and demonstrated negative or markedly reduced BAG-1L staining in BAG-1L specific TALEN knockout LNCaP cells, BAG-1 shRNA treated LNCaP cells and BAG-1 siRNA treated HeLa cells compared to control cells (**Supplementary Figure 17C**). In addition, BAG-1L is predominantly localized within the nucleus, consistent with its unique nuclear localization sequence within its N-terminus (**Supplementary Figure 17C**). Having confirmed RM310 BAG-1L specificity, we performed IHC on formalin-fixed, paraffin-embedded PC patient tissue biopsies within our study cohorts, demonstrating strong and predominantly nuclear staining (**Supplementary Figure 18A-B and 19A**).

We next used this validated antibody to investigate BAG-1L protein expression in same-patient, matched biopsies, as 43 patients progressed from castration sensitive prostate cancer (CSPC) to CRPC (ICR/RMH patient IHC cohort) (**Supplementary Figure 18A, Supplementary Table 9**). In this cohort, nuclear BAG-1L expression did not significantly (P = 0.15, Student t-test) increase as patients progressed from CSPC (median H score, interquartile range [IQR]; 50, 14-90) to CRPC (80, 10-95) (**Supplementary Figure 19B**). In addition, nuclear BAG-1L expression was significantly different (P = 0.04, Student t-test) in CRPC biopsies (n = 67, H score 80, IQR 17-95) taken before abiraterone acetate (AA) or enzalutamide (E) therapy (H score 80, IQR 52.5-102.5) and after AA and/or E therapy (H score 70, 9.75-9.5), although these differences were not marked (**Supplementary Figure 19C**). In contrast to this, nuclear BAG-1L expression in a second independent clinical cohort (UW/FHCRC patient IHC cohort), was slightly higher (P = 0.05, Student t-test) when comparing primary localized prostate cancer (H score 0, IQR 0-50) and (unmatched) metastatic CRPC (H score 40, IQR 3-60) (**Supplementary Figure 18B and 19D, Supplementary Table 10**). We next investigated the impact of nuclear BAG-1L expression at diagnosis on patient outcome. Patients with lower nuclear BAG-1L expression (H score ≤ 50, n = 22) at CSPC did not show significantly different median time to CRPC (21.1 vs 20.3 months, HR 1.28, 95% CI 0.70-2.34, P = 0.40), or median overall survival (74.8 vs 74.5 months, HR 0.96, 95% CI 0.51-1.79, P = 0.89), when compared to patients with higher BAG-1L protein expression (H score > 50, n = 21) (**Supplementary Figure 18A and 19E-F**). To investigate the impact of nuclear BAG-1L expression on response to current AR targeting therapies we determined the response of the ICR/RMH patient IHC cohort to AA or E following chemotherapy in nuclear BAG-1L low (H score ≤ 70; n = 26) and high (H score > 70; n = 24) expressing CRPC biopsies (**Supplementary Figure 18A and 20A-D, Supplementary Table 11**). Patients with lower nuclear BAG-1L expression did not have significantly different 12-week PSA response rate (35 vs 38%, P > 0.99, Fisher’s exact test), median time to PSA progression (2.7 vs 3.0 months, HR 0.90, 95% CI 0.50-1.57, P = 0.68), median time to radiological/clinical progression (5.9 vs 4.3 months, HR 0.68, 95% CI 0.38-1.19, P = 0.15), or median overall survival (18.5 vs 14.8 months, HR 0.73, 95% CI 0.41-1.30, P = 0.27), when compared to patients with higher BAG-1L protein expression (**Supplementary Figure 20A-D**). Taken together, these data indicate that BAG-1L expression at diagnosis or at time of CRPC biopsy does not associate with clinical outcome in men with advanced prostate cancer.

## Discussion

Here we show for the first time that Thio-2 inhibits AR signaling and growth in CRPC patient derived models. In addition, we demonstrate that Thio-2 may bind the AR N-terminus and destabilize AR-DNA interactions, providing new insights into its mechanism of action. In contrast, genetic abrogation of all BAG-1 isoforms did not inhibit AR signaling and growth in CRPC models, and the Thio-2 related phenotype was maintained in CRPC models with all BAG-1 isoforms knockdown/out, suggesting its activity is mainly independent of the BAG domain, consistent with our computer modelling that demonstrates the BAG domain to be a challenging drug target. Furthermore, we demonstrate that panBAG-1 knockout is not critical for normal mouse physiological development, although this did decrease prostatic weight, with all BAG-1 isoform knockout not impacting cancer initiation or progression in the TRAMP and PTEN loss mouse models. Finally, a novel IHC assay, using a BAG-1L specific antibody, identified BAG-1L protein expression to be high in advanced prostate cancer, although expression levels did not associate with clinical outcome. These important data suggest that therapies, such as Thio-2, that target persistent AR signaling are an attractive therapeutic strategy for CRPC, but underline that targeting the BAG domain of all BAG-1 isoforms requires further investigation to be considered an attractive therapeutic strategy to abrogate persistent AR signaling in CRPC.

It is important to consider these data, and the limitations of the study, in the context of historical studies. Thio-2, a novel compound derived from Thioflavin S, has been reported to inhibit AR signaling and CSPC model growth (14, 25). Thio-2 has been predicted to bind the BAG domain of BAG-1 isoforms though *in-silico* docking experiments, and studies in melanoma, breast and prostate cancer cell lines, have suggested a reduction in binding of BAG-1 to its binding partners (such as HSC/P70, BRAF and AR), to inhibit AR, MEK and AKT signaling (14, 25, 58). These studies support the hypothesis that Thio-2 functions though the BAG domain, however, our current studies demonstrate that Thio-2 may bind the AR N-terminus through a similar mechanism to EPI-001, and destabilizes genome-wide AR binding, suggesting a previously undescribed alternative mechanism of action (55). This is further supported by our studies that show that Thio-2 suppresses AR signaling in CRPC models with all BAG-1 isoform knockdown/out, suggesting the pharmacological activity is independent of the BAG domain. Although the identification of a potential interaction with the AR N-terminus and suppression of AR signaling is promising it is conceivable that at high concentration the low solubility of Thio-2 may result in cellular stress and lead to changes in transcription and/or translation which manifest as indirect downregulation of AR signaling. This is an important consideration for Thio-2, or any novel compound thought to directly impact AR and/or AR signaling. It is now critically important that Thio-2 derivatives, such as A4B17, that are reported to disrupt the BAG-1L:AR NTD interaction, suppress AR signaling, and inhibit prostate cancer model growth are subjected to a rigorous interrogation to validate these findings (26, 27).

Our studies showed little impact of knockdown/out of all BAG-1 isoforms on AR signaling and prostate cancer cell growth. Unlike other studies that focused on CSPC models, our studies have focused on basal AR signaling and CRPC models, suggesting that BAG-1L may play a more important role in regulating the stimulated AR (14–17). One potential limitation of these studies, specific to our LNCaP95 CRISPR clones, is the challenges of a heterogenous cell population giving rise to biological differences independent of the genomic manipulation performed (59). Interestingly, studies in breast cancer cell line models have demonstrated differential response in growth to BAG-1 knockdown and it maybe that the biological background is important for sensitivity to BAG-1 abrogation which will be important to understand in the context of prostate cancer (58, 60). This finding is important as drugs targeting the BAG domain would presumably impact all three isoforms, and not resemble the impact of BAG-1L specific knockouts or mutations previously described (14–17). There might remain a requirement for BAG-1S and BAG-1M mediated interactions to observe the phenotype associated with BAG-1L knockdown alone, although this seems unlikely. Taken together, the clear differences between BAG-1L specific knockdown and all BAG-1 isoform knockdown/out will need to be further investigated if targeting the BAG domain of BAG-1L, and other BAG-1 isoforms, is to be considered for therapeutic development.

In contrast to previous studies, BAG-1 knockout had limited impact on normal mouse physiological development. Interestingly, male mice had smaller prostates, and pregnant female mice had increased duration of pregnancy, decreased litter size, increased neonatal weight on day 1 and decreased neonatal survival on day 2, which may point to a role in hormonal regulation. Although there were differences in prostate size, there were no significant alternations in molecular pathways when mRNA transcripts were compared between BAG-1 knockout and wildtype mouse prostates. Previous studies reported BAG-1 knockout to be embryonically lethal, likely due to BAG-1 and CHMP5 co-deletion, with CHMP5 loss being the driver of the observed phenotype (44, 56). This is consistent with the development of BAG-1 knockout embryonic stem cells that maintained pluripotency and the ability to differentiate (61). These data suggest that therapies targeting BAG-1 may be associated with limited treatment-related toxicity since BAG-1 knockout had limited impact on normal mouse physiological development. BAG-1 knockout in the TRAMP and PTEN loss models of prostate cancer did not, however, decrease prostatic weight or aggressive histology. These data, despite the limitations of the models studied, do not support BAG-1 being a validated target for prostate cancer therapy.

Finally, we developed and analytically validated a novel BAG-1L specific IHC assay. We demonstrated that BAG-1L protein expression was higher in CRPC metastasis compared to unmatched, untreated, castration sensitive prostatectomies, consistent with previous studies (23, 24). We next investigated changes in BAG-1L protein expression as patients developed CRPC, using matched, same patient, CSPC and CRPC samples. Interestingly, BAG-1L protein expression did not significantly change as patients progressed from CSPC to CRPC. This may not be unexpected, as all patients studied with CSPC developed CRPC, and genomic studies have shown these patients to have similar genomics at diagnosis to when CRPC develops, and this is different to localized CSPC in which the majority of patients do not develop CRPC (62). These results are different to the study of an antibody to all BAG-1 isoforms, where nuclear BAG-1 protein expression increased as patients progressed from CSPC to CRPC (14). This, in part, may be due to shorter BAG-1 isoforms (BAG-1M and BAG-1S) localizing to the nucleus under conditions of cellular stress (19). Multiple studies have demonstrated nuclear BAG-1 protein expression to associate with clinical benefit from AR-targeting therapies, and cytoplasmic BAG-1 protein expression to associate with benefit from radiotherapy in localized disease (14, 23, 24). Our current study demonstrates no association between BAG-1L protein expression at diagnosis and time to CRPC or overall survival, and no association between BAG-1L protein expression at CRPC and clinical benefit from AR-targeting therapies. These differences are unsurprising as analytical validation and clinical qualification of predictive and prognostic biomarkers for prostate and other cancers is challenging, and these studies have multiple variables including preanalytical variables, different antibodies, heterogenous patient cohorts and different quantification strategies (63, 64). The clinical significance of BAG-1L in prostate cancer remains uncertain and will require further interrogation to determine whether it should be further considered as a predictive or prognostic biomarker in prostate cancer.

Overall, these studies demonstrate that therapies targeting the AR N-terminus to suppress persistent AR signaling remain an attractive therapeutic strategy in CRPC. In contrast to studies focused on BAG-1L and the activated AR, however, targeting the BAG domain of BAG-1 to inhibit BAG-1L and other BAG-1 isoforms was not validated as a therapeutic strategy in CRPC by our current efforts.

## Acknowledgments

The authors gratefully acknowledge the patients and the families of patients who contributed to this study. This work was supported by the Claudia Adam Barr Foundation (Funding to L.C.), Prostate Cancer UK (Travelling Prize Fellowship to J.M.J.V.; Research Funding to J.S.dB. and A.Sharp), the Asociación Española contra el Cáncer (Funding to B.M.), Medical Research Council (Clinical Research Training Fellowship to A.Sharp; Research Funding to J.S.dB.), The Academy of Medical Sciences (Starter Grant for Clinical Lecturers to A.Sharp), the Prostate Cancer Foundation (Young Investigator Awards to L.C, A.Sharp; Challenge Awards to L.C., S.R.P., M.B., A.C.B.C., J.S.dB. and A.Sharp), Cancer Prevention and Research Institute of Texas (Funding to B.A-L.), the Movember Foundation through the London Movember Centre of Excellence (CEO13 2-002 to J.S.dB.), the Wellcome Trust (Clinical Research Career Development Fellowship to A.Sharp), the National Institute for Health and Care Research Biomedical Research Centre (Funding to A.Sharp), the Veterans Affairs Research and Development Service (to S.R.P.), and Cancer Research UK (Drug Discovery Committee Strategic Award to B.A-L.; Centre Programme and Experimental Cancer Medicine Centre grants to J.S.dB.).

