## Supplemental Figure Legends and Tables for "Targeting the BAG-1 family of co-chaperones in lethal prostate cancer"

### Supplementary Figure Legends

#### Supplementary Figure 1: Development of castration resistant prostate cancer patient derived xenograft organoids

**(A)** Patient derived xenografts (PDX) CP50, CP89 and CP142 were derived from metastatic lymph node biopsies of patients with castration resistant prostate cancer (CRPC) treated with multiple standard treatments for castration sensitive prostate cancer (CSPC) and CRPC. Staging (metastasis (M) yes (1) or no (0)) and histology (Gleason score (G) with or without neuroendocrine differentiation (NE) at diagnosis, treatments received for CSPC and CRPC, and timing of metastatic biopsy are shown. Once PDX were established (> 3 passages), PDX-organoids (PDX-O) were derived from fresh PDX tumors and plated in 96 well plates for drug treatment experiments presented. **(B)** CP50, CP89 and CP142 PDX tumors and derived PDX-Os were formalin-fixed paraffin-embedded and androgen receptor (AR-FL) and pan BAG-1 (panBAG-1) immunohistochemistry (IHC) performed. Representative micrographs of AR-FL and pan BAG-1 detection by IHC are shown. Scale bar, 50 µm.

#### Supplementary Figure 2: Validation of pan BAG-1 immunohistochemistry

Representative western blot and immunohistochemistry (IHC) of pan BAG-1 (panBAG-1) detection using a pan BAG-1 antibody in HeLa cells treated with control (siCnt) or BAG-1 (siBAG-1) siRNA. Scale bar, 50 µm.

#### Supplementary Figure 3: Thio-2 inhibits the growth of castration resistant prostate cancer patient derived xenograft organoids with different molecular subtypes

**(A)** CP89 and **(B)** CP142 PDOs were treated with vehicle (DMSO 0.1 %) or various concentrations Thio-2 (0.1, 1, 5, 10, 25 and 50 µM) or enzalutamide (1 and 10 µM) and growth determined after 5 days by CellTiter-Glo® 3D Cell Viability Assay. Mean fold change in growth (compared to day 0) with standard deviation from a single experiment with six replicates is shown. P values were calculated for each condition compared to vehicle using unpaired Student t-test. P values $\leq 0.05$ are shown.

**Supplementary Figure 4: *In-vivo* treatment of patient derived model of lethal prostate cancer demonstrates Thio-2 to be well tolerated with associated impact on AR signaling and growth**

**(A-C)** The impact of vehicle (5 % DMSO in 10 % (w/v) HBC in 0.9 % saline) and 15 mg/kg Thio-2 once daily (OD) intraperitoneal (IP) on NSG mice (n=3 per arm; A) on heart, kidney, testes, seminal vesicles and prostate weights (B), and total body weight (C) was determined. P values were calculated for vehicle compared to 15 mg/kg Thio-2 OD IP using unpaired Student t-test. **(D)** Schematic overview of experimental design using patient derived xenograft (PDX) CP50 that was developed from a lymph node biopsy from a patient who had progressed through all standard of care treatments for castration resistant prostate cancer. Once CP50 PDX tumor volume reached 300 $mm^3$ treatment, 15 mg/kg Thio-2 or vehicle OD IP was commenced for 14 days. **(E)** Mean growth (normalized to day 0; defined as 1) with standard deviation was determined for each tumor. P values were calculated comparing to 15 mg/kg Thio-2 IP OD treatment arm to vehicle control at 11 to 15 days using unpaired Student t-test. **(F)** The effect of 15 mg/kg Thio-2 OD IP compared to vehicle on serum PSA was determined at 14 days. Mean serum PSA with standard deviation was determined for each mouse. P values were calculated for vehicle compared to 15 mg/kg Thio-2 OD IP using unpaired Student t-test. **(G)** The effect of 15 mg/kg Thio-2 OD IP compared to vehicle on AR-FL, PSA and GAPDH protein expression was determined at 14 days. Singel western blot is shown. **(H)** The effect of 15 mg/kg Thio-2 OD IP compared to vehicle on BAG-1, AR-FL, AR-V7, PSA, TMPRSS2 and FKBP5 mRNA expression was determined. Mean mRNA expression (normalized to average of GAPDH/B2M/HRPT1/RPLP0 and vehicle treated; defined as 1) with standard deviation for five mice per arm with two replicates is shown. P values were calculated for 15 mg/kg Thio-2 OD IP compared to vehicle using unpaired Student t-test.

**Supplementary Figure 5: Thio-2 reduces genome-wide AR binding in LNCaP prostate cancer cells**

LNCaP cells were grown in starved media (10 % charcoal stripped serum) for 72 hours, following 1 hour pre-treatment with vehicle or 5 µM Thio-2. Cells were subsequently treated with vehicle (Ethanol 0.1%) or 10 nM dihydrotestosterone (DHT) for 16 hours (17 hours total treatment). Chromatin immunoprecipitation sequencing (ChIPseq) was performed on a single experiment in triplicate. Peak density histograms and heatmaps showing AR binding regions from ChIPseq analysis for vehicle (DMSO 0.1 %) and 5 µM Thio-2 treated samples are shown. For starved (Ethanol 0.1 %) and stimulated (10 nM DHT) samples, AR binding within a 4 kilobases window is shown for pooled peak summits from both vehicle (DMSO 0.1 %) and 5 µM Thio-2 treated samples. Colour bars show relative signal intensity.

**Supplementary Figure 6: Inhibition of growth and androgen receptor signaling by Thio-2 in the prostate cancer cell lines LNCaP95 is not exclusively dependent on BAG-1**

**(A)** LNCaP95 prostate cancer cells were transfected with 50 nM of either control (siCnt; clear bars) or BAG-1 (siBAG-1; red bars) siRNA for 72 hours prior to treatment with vehicle (0.1 % DMSO) or various concentrations (5, 10, 25 and 50 µM) of Thio-2 and growth was determined after 6 days by CellTiter-Glo® Luminescent Cell Viability Assay. Mean fold change in growth (compared to day 0) with standard deviation from a single experiment with six replicates is shown. P values were calculated for each condition compared to vehicle in siCnt and siBAG-1 cells, and between vehicle treated siCnt and siBAG-1 cells (grey shading), using unpaired Student t-test. P values $\leq$ 0.05 are shown and P values $>$ 0.05 are shown as non-significant (ns). **(B)** LNCaP95 prostate cancer cells were transfected with 50 nM of either siCnt or siBAG-1 siRNA for 55 hours prior to treatment with vehicle (0.1 % DMSO) or various concentrations (5 and 50 µM) of Thio-2 for 17 hours (total 72 hours) and AR-FL, AR-V7, PSA, BAG-1 and GAPDH protein expression was determined. Single western blot result is shown. **(C)** LNCaP95 prostate cancer cells were transfected with 50 nM of either siCnt or siBAG-1 siRNA for 55 hours prior to treatment with vehicle (DMSO 0.1 %) or various concentrations (5 and 50 µM) of Thio-2 for 17 hours (total 72 hours) and BAG-1, AR-FL, AR-V7, PSA, TMPRSS2 and FKBP5 mRNA expression was determined. Mean mRNA expression (normalized to average of GAPDH/B2M/HRPT1/RPLP0 and siCnt/vehicle; defined as 1) with standard deviation from a single experiment with six replicates is shown. P values were calculated for each condition compared to vehicle in siCnt and siBAG-1 cells, and between vehicle treated siCnt and siBAG-1 cells (grey shading), using unpaired Student t-test. P values ≤ 0.05 are shown and P values > 0.05 are shown as non-significant (ns).

**Supplementary Figure 7: Inhibition of growth and androgen receptor signaling by Thio-2 in the prostate cancer cell line LNCaP95 is not exclusively dependent on BAG-1**

**(A)** BAG-1 knockout CRISPR/Cas9 clones were developed in LNCaP95 prostate cancer cells. Control (Cas9) and two BAG-1 knock-out (guide 3, g3; guide 4, g4) were used for transfection and single cell derived LNCaP95 clones were selected. Clones were treated with vehicle (DMSO 0.1 %) or various concentrations (5 and 50 µM) of Thio-2 and growth was determined after 6 days by CellTiter-Glo® Luminescent Cell Viability Assay. Mean growth (normalized to vehicle treated control clone; defined as 1) with standard deviation from three separate experiments performed in duplicate is shown. P values were calculated for each condition compared to vehicle for each individual guide, and between vehicle treated control and BAG-1 guides (grey shading), using unpaired Student t-test. P values ≤ 0.05 are shown and P values > 0.05 are shown as non-significant (ns). **(B)** LNCaP95 clones were treated with vehicle (DMSO 0.1 %) or various concentrations (5 and 50 µM) of Thio-2 for 17 hours and AR-FL, AR-V7, BAG-1 and GAPDH protein expression was determined. Single western blot is shown. **(C)** LNCaP95 clones were treated with vehicle (DMSO 0.1%) or various concentrations (5 and 50 µM) of Thio-2 for 17 hours and BAG-1, AR-FL, AR-V7, PSA, TMPRSS2 and FKBP5 mRNA expression was determined. Mean mRNA expression (normalized to average of GAPDH/B2M/HRPT1/RPLP0 and vehicle treated control clone; defined as 1) with standard deviation from a single experiment with three replicates is shown. P values were calculated for each condition compared to vehicle for each individual guide, and between vehicle treated control and BAG-1 guides (grey shading), using unpaired Student t-test. P values ≤ 0.05 are shown and P values > 0.05 are shown as non-significant (ns).

**Supplementary Figure 8: Thio-2, despite limitations of solubility, may bind the N-terminus of the androgen receptor through a similar binding mode to EPI-001**

**(A-D)** Nuclear magnetic resonance (NMR) 1H-15N correlation spectra were recorded on 600 and 800 MHz Bruker Advance spectrometers equipped with cryoprobes. Experiments with 15N-labelled androgen receptor N-terminus (AR-NTD) constructs $NTD_{1-518}$ and $NTD_{330-447}$ at 25 µM were mixed with an excess amount (250 µM) of Thio-2 or EPI-001 (positive control) and measured at 5 ºC. This temperature was selected to prevent the protein to aggregate, and in consequence, affecting the resolution of the spectra. **(A and C)** Quantification of intensity changes in AR-NTD ($NTD_{1-518}$ construct) backbone amide signals in the presence of EPI-001 (A) or Thio-2 (C). **(B and D)** Quantification of chemical shift perturbations in AR-NTD ($NTD_{330-447}$ construct) backbone amide signals in the presence of EPI-001 (B) or Thio-2 (D). Colored shaded boxes indicate the most affected regions.

**Supplementary Figure 9: Thio-2 demonstrates low solubility**

Thio-2 solubility was measured by integration of Thio-2 1H aromatic signals (region 6.5-8 ppm) and the internal reference (10 µM DSS) 1H signal (at 0 ppm). A sample containing 5 µM Thio-2, 10 µM DSS, and 0.5 % DMSO (buffer 20 mM sodium phosphate (pH 7.4), 1 mM TCEP, 10% $D_20$) were recorded on 600 MHz Bruker Avance spectrometer equipped with a cryoprobe. 1H nuclear magnetic resonance (NMR) spectra of 5 µM Thio-2 in phosphate buffer containing 0.5 % DMSO measured at 37 ºC is shown.

**Supplementary Figure 10: Development and characterization of BAG-1 knockout mice**

**(A)** Microarray data from BAG-1 wildtype (WT) and exon 1 BAG-1 heterozygous knockout (Het KO) mouse prostates following up to 12-weeks castration demonstrating down-regulation of BAG-1 and CHMP5. $Log_2$ expression of those genes most altered is shown. P values were calculated for differences in gene expression between BAG-1 WT and BAG-1 Het KO using unpaired Student t-test. P values ≤ 0.05 are shown (*). **(B)** Representative western blot and immunohistochemistry (IHC) of BAG-1 detection using a panBAG-1 antibody in

NIH3T3 cells treated with control (siCnt) or BAG-1 (siBAG-1) siRNA. Scale bar, 50 μm. **(C)** Prostates from BAG-1 knockout (BAG-1 KO) mouse strain Bag1tm1a(EUCOMM)Hmgu and BAG-1 WT male mice were analyzed for mouse (mo) BAG-1 protein (immunohistochemistry; IHC) levels. Representative micrographs of BAG-1 detection in mouse prostates by pan-BAG-1 antibody IHC are shown. Scale bar, 200 µm. **(D)** Western blots on lysates of prostates from BAG-1 KO (n = 4) and BAG-1 wildtype (WT; n = 4) male mice were analyzed for BAG-1 and GAPDH protein levels. Single western blot is shown. **(E)** Mouse prostates from BAG-1 KO (red bars; n = 5) and BAG-1 WT (gray bars; n = 6) male mice at age 12 weeks were taken and samples prepared for RNA sequencing. Median BAG isoforms and CHMP5 mRNA levels (fragments per kilobase of transcript per million reads; FPKM) with interquartile range, and smallest and largest value, is shown. P values were calculated for BAG-1 KO compared with BAG-1 WT mice using unpaired Student t-test. **(F)** Overview of gene set enrichment analysis (GSEA) using MsigDB (v7.0) functional pathways: H, Hallmark; C2, Curated Gene Sets (including KEGG, Biocarta, Reactome). BAG-1 KO and BAG-1 WT RNA sequencing analysis was compared, change in gene expression was ranked by $\log_2$ fold change, then tested for enrichment against functional gene sets (H, C2), using GSEA. The distribution of resulting FDR values ($\log_{10}$) is shown in red, and the threshold for significance (FDR < 0.05) is shown by dotted grey line. None of the pathways tested reached the significance threshold.

**Supplementary Figure 11: BAG-1 knockout female mice are viable and fertile**

**(A)** Kaplan-Meier curves of overall survival (OS) of BAG-1 KO (red line; n=9) and BAG-1 WT (gray line; n=10) female mice from birth. Median OS, hazard ratio (HR) with 95% confidence intervals and P values for univariate Cox survival model are shown. **(B)** The body weight of BAG-1 KO (red bars) and BAG-1 WT (gray bars) female mice at 12 weeks and 12 months was determined. Median body weight with interquartile range, and smallest and largest value, is shown. P values were calculated for BAG-1 KO compared with BAG-1 WT mice using unpaired Student t-test. **(C)** The weight of the genitourinary tract and uterus from BAG-1 KO (red bars) and BAG-1 WT (gray bars) female mice at 3 months and older was determined. Median weight with interquartile range, and smallest and largest value, is shown. P values were calculated for BAG-1 KO compared with BAG-1 WT mice using unpaired Student t-test. **(D)** The duration of pregnancy, litter size, neonatal weight (on day 1), neonatal deaths (per litter) and neonatal survival rate (on day 2) for female BAG-1 KO (red bars) and BAG-1 WT (gray bars) mice was determined. Median values with interquartile range, and smallest and largest value, is shown. P values were calculated for BAG-1 KO compared with BAG-1 WT mice using unpaired Student t-test.

**Supplementary Figure 12: Mouse organ hematoxylin and eosin, and BAG-1 immunohistochemistry, in BAG-1 knockout mice**

Organs (brain, heart, kidney, liver, and lung) from BAG-1 KO and BAG-1 WT mice at 4 months and older were analyzed by hematoxylin and eosin (H & E) and for mouse (mo) BAG-1 protein (IHC) levels. Representative micrographs of H & E, and BAG-1 detection, in mouse organs by pan-moBAG-1 antibody IHC are shown. Scale bar, 200 µm.

**Supplementary Figure 13: Mouse organ hematoxylin and eosin, and BAG-1 immunohistochemistry, in BAG-1 knockout mice**

Organs (spleen, seminal vesicles, testis, and thymus) from BAG-1 KO and BAG-1 WT mice at 3 months and older were analyzed by H & E and for mouse (mo) BAG-1 protein (IHC) levels. Representative micrographs of H & E, and BAG-1 detection, in mouse organs by pan-moBAG-1 antibody IHC are shown. Scale bar: 200 µm.

**Supplementary Figure 14: Validation of mouse androgen receptor immunohistochemistry**

Representative western blot and immunohistochemistry (IHC) of androgen receptor (AR) detection using an AR N-terminal specific antibody in TRAMPC2 cells treated with control (siCnt) or AR (siAR) siRNA. Scale bar: 50 µm.

**Supplementary Figure 15: Impact of BAG-1 knockout in the TRAMP transgenic mouse model**

**(A)** Prostates from TRAMP transgenic (TG)/BAG-1 KO mice and TRAMP TG/BAG-1 WT male mice were analyzed at 6 months of age by H & E, and for BAG-1 and AR protein (IHC) levels. Representative micrographs (* shown in figure 2) of H & E, and BAG-1 and AR detection, in prostates are shown. Scale bar, 100 µm. **(B)** Quantification (H-score) of nuclear and cytoplasmic BAG-1 IHC was performed. Median H-score with interquartile range, and smallest and largest value, is shown. P values were calculated for TRAMP TG/BAG-1 KO compared with TRAMP TG/BAG-1 WT mice using unpaired Student t-test. **(C)** Quantification (H-score) of nuclear and cytoplasmic AR IHC was performed. Median H-score with interquartile range, and smallest and largest value, is shown. P values were calculated for TRAMP TG/BAG-1 KO compared with TRAMP TG/BAG-1 WT mice using unpaired Student t-test.

**Supplementary Figure 16: Impact of BAG-1 knockout in the inducible prostate specific PTEN knockout mouse model**

**(A)** Prostates from PTEN KO/BAG-1 KO mice and PTEN KO/BAG-1 WT male mice were analyzed by H & E, and for BAG-1 and AR protein (IHC) levels. Representative micrographs (* shown in figure 2) of H & E, and BAG-1 and AR detection, in prostates of 14 months old mice are shown. Scale bar, 100 µm. **(B)** Quantification (H-score) of nuclear and cytoplasmic BAG-1 IHC was performed. Median H-score with interquartile range, and smallest and largest value, is shown. P values were calculated for PTEN KO/BAG-1 KO compared PTEN KO/BAG-1 WT mice using unpaired Student t-test. **(C)** Quantification (H-score) of nuclear and cytoplasmic AR IHC was performed. Median H-score with interquartile range, and smallest and largest value, is shown. P values were calculated for PTEN KO/BAG-1 KO compared PTEN KO/BAG-1 WT mice using unpaired Student t-test.

**Supplementary Figure 17: Analytical validation of a novel BAG-1L antibody (clone RM310) for immunohistochemistry**

**(A)** Western blot of VCaP and 22Rv1 cells, and BAG-1L specific transcription activator-like effector nuclease (TALEN) knockout LNCaP cells, BAG-1 shRNA treated LNCaP cells (clone 506) and BAG-1 siRNA treated HeLa cells, compared to control cells, using a novel recombinant rabbit monoclonal anti–BAG-1L antibody (clone

RM310) and pan BAG-1 antibody (clone 3.10G3E2). (**B**) Immunoprecipitation of BAG-1L from 22Rv. cells using the BAG-1L specific antibody (clone RM310) and western blot performed with pan BAG-1 antibody (clone 3.10G3E2). (**C**) Micrographs of BAG-1L detection by IHC using the BAG-1L specific antibody (clone RM310) in cell line pellets from VCaP and 22Rv1 cells, and BAG-1L specific TALEN knockout LNCaP cells, BAG-1 shRNA treated LNCaP cells (clone 506) and BAG-1 siRNA treated HeLa cells, compared to control cells. Scale bar: 50 µm.

**Supplementary Figure 18: Overview of Institute of Cancer Research/Royal Marsden Hospital and University of Washington/Fred Hutchinson Cancer Research Center prostate cancer immunohistochemistry cohorts.**

**(A)** Overview of the Institute of Cancer Research/Royal Marsden Hospital (ICR/RMH) immunohistochemistry (IHC) cohort. The ICR/RMH IHC cohort included 43 castration sensitive prostate cancer (CSPC) biopsies and 67 castration resistant prostate cancer (CRPC) biopsies stained for BAG-1L expression. Response data were available for abiraterone acetate (AA) or enzalutamide (E) after chemotherapy in 50 patients (compared to BAG-1L expression at CRPC) and for time to CRPC and overall survival (OS) from diagnosis for 43 patients (compared to BAG-1L expression at CSPC). **(B)** Overview of the University of Washington/Fred Hutchinson Cancer Research Center (UW/FHCRC) prostate cancer IHC cohort. The UW/FHCRC IHC cohort included 30 primary prostatectomies and 30 CRPC metastases stained for BAG-1L expression.

**Supplementary Figure 19: BAG-1L protein expression in prostate cancer.**

**(A)** Representative micrographs of BAG-1L detection by immunohistochemistry (IHC) in 4 Institute of Cancer Research/Royal Marsden Hospital (ICR/RMH) patients with matched castration sensitive prostate cancer (CSPC) and castration resistant prostate cancer (CRPC) biopsies. Scale bar: 50 µm. Prostate biopsies (Prostate Bx), transurethral resection of the prostate (TURP) and bone marrow trephine (BMT) biopsies are shown. **(B)** Nuclear BAG-1L expression (H-score, HS) in 43 same-patient matched CSPC (gray) and CRPC (red) biopsies from the ICR/RMH cohort is shown. Median HS with interquartile range, and smallest and largest value, is shown. P values were calculated for BAG-1L expression at CSPC compared to CRPC using paired

Student t-test. **(C)** Nuclear BAG-1L expression (HS) in 67 CRPC (red) biopsies and dichotomized (orange) by pre (25 biopsies) and post (42 biopsies) abiraterone acetate (AA) or enzalutamide (E) treatment. Median HS with interquartile range, and smallest and largest value, is shown. P values were calculated for BAG-1L expression pre and post AA or E using unpaired Student t-test. **(D)** Representative micrographs of BAG-1L detection by IHC in 2 University of Washington/Fred Hutchinson Cancer Research Center (UW/FHCRC) patients showing a primary prostatectomy and metastasis. Scale bar, 50 µm. Nuclear BAG-1L expression (HS) in 30 primary (gray) and 30 metastatic (red) prostate cancer biopsies from the UW/FHCRC cohort is shown. Median HS with interquartile range, and smallest and largest value, is shown. P values were calculated for BAG-1L expression in primary prostate cancer compared to metastatic prostate cancer using unpaired Student t-test. **(E-F)** The impact of BAG-1L expression (dichotomized by median HS) at CSPC on time to CRPC and overall survival was determined for 43 patients. Kaplan-Meier curves show time to CRPC progression (E) and overall survival (F) from diagnosis. Hazard ratios (HR) with 95% confidence intervals (CI) are shown. P value was calculated using univariate Cox proportional hazards model.

**Supplementary Figure 20: BAG-1L protein expression in castration resistant prostate cancer and response to androgen receptor targeting therapies post chemotherapy**

**(A)** Percentage 12-week 50% PSA response rate on AR-targeting therapies after chemotherapy for patients with low nuclear BAG-1L (≤ median, gray) and high nuclear BAG-1L (> median, red) CRPC biopsy protein expression is shown. Twelve-week 50% PSA response rate is shown. P value was calculated using Fisher’s exact test. Kaplan-Meier curves show time to PSA progression (PD) (**B**), time to clinical/radiological PD (**C**), and overall survival (**D**) from start of AR-targeting therapy. Hazard ratios (HR) with 95% confidence intervals (CI) are shown. P value was calculated using univariate Cox proportional hazards model.

### Supplementary Tables

| Protein target (species, clone, supplier) | Retrieval buffer (method) | Dilution (time) | Detection | Controls |
|---|---|---|---|---|
| BAG-1L (human, RM310, RevMAb) | pH6 citrate (microwave) | 1:1000 (1 hour) | DAKO EnVision Detection System | BAG-1L specific TALEN knockout (neg) and control (pos) LNCaP; VCaP (pos) |
| panBAG-1 (human, RM356, RevMAb) | pH8.1 Tris/EDTA (microwave) | 1:1500 (1 hour) | DAKO EnVision Detection System | siBAG-1 (neg) and siControl (pos) HeLa; SKMEL2 (pos) |
| panBAG-1 (mouse, AF815, R&D systems) | pH6 ER1 30min (Leica Biosystems Bond RX) | 1:500 (1 hour) Rabbit anti-goat IgG secondary (Abcam) 1:5000 (30min) | Leica Novolink Polymer Detection System | siBAG-1 (neg) and siControl (pos) NIH 3T3 |
| AR-FL (mouse/human, EPR1535, abcam) | pH6 citrate (pressure cooker) | 1:500 (1 hour) | DAKO EnVision Detection System | siAR (neg) and siControl (pos) TRAMPC2, mouse normal prostate |

**Supplementary Table 1: Protocols and antibodies for immunohistochemical analysis.**

pos – positive, neg – negative.

| Protein target | Species (clone) | Company | Catalogue ID |
|---|---|---|---|
| panBAG-1 | mouse (AF815) | R&D systems | AF815 |
| GAPDH | mouse/human (G9) | Santa Cruz | sc-365062 |
| AR-FL | mouse/human (EPR1535(2)) | abcam | ab133273 |
| BAG-1L | human (RM310) | RevMAb | 31-1196-00 |
| panBAG-1 | human (RM356) | RevMAb | 31-1242-00 |
| AR-FL | human (D6F11) | Cell Signaling | 5153 |
| AR-V7 | human (RM7) | RevMAb | 31-1109-00 |
| PSA | human (D11E1) | Cell Signaling | 2475 |
| Vinculin | mouse/human (hVIN-1) | Sigma-Aldrich | V9131 |
| Beta-actin | Mouse/human (C4) | Santa Cruz | sc-47778 |

**Supplementary Table 2: Antibodies used for western blot analysis.**

| Gene target | Company | Catalogue ID |
|---|---|---|
| mouse BAG-1 exon 1/2 | ThermoFisher | Mm01208597_g1 |
| mouse BAG-1 exon 5/6 | ThermoFisher | Mm01208593_m1 |
| mouse BAG-1 exon 6/7 | ThermoFisher | Mm00437765_m1 |
| mouse GAPDH | ThermoFisher | Mm99999915_m1 |
| human BAG-1 | ThermoFisher | Hs00185390_m1 |
| human AR-FL | ThermoFisher | Hs00171172_m1 |
| human AR-V7 | ThermoFisher | Hs04260217_m1 |
| human PSA | ThermoFisher | Hs02576345_m1 |
| human TMPRSS2 | ThermoFisher | Hs05024838_m1 |
| human FKBP5 | ThermoFisher | Hs01561006_m1 |
| human GAPDH | ThermoFisher | Hs02786624_g1 |
| human B2M | ThermoFisher | Hs00187842_m1 |
| human HRPT1 | ThermoFisher | Hs00363810_m1 |
| Human RPLP0 | ThermoFisher | Hs00420895_gH |

**Supplementary Table 3: TaqMan probes for qRT-PCR analysis**

| Cell line | Supplier | Catalogue ID | Media^ |
|---|---|---|---|
| 22Rv1 | ATCC | CRL-2505 | RMPI1640/10% FCS |
| VCaP | ATCC | CRL-2876 | DMEM/10% FCS |
| LNCaP | ATCC | CRL-1740 | RMPI1640/10% FCS |
| LNCaP95 | Dr Meeker/Dr Luo* | does not apply | RMPI 1640^^/10%CCS |
| NIH3T3 | ATCC | CRL-1658 | DMEM/10% FCS |
| TRAMPC2 | ATCC | CRL-2731 | DMEM/10% FCS |
| HeLa | ATCC | CCL-2 | DMEM/10% FCS |

**Supplementary Table 4: Cell lines**

ATCC – American Type Culture Collection, FBS – fetal bovine serum, CSS – charcoal stripped serum, * - LNCaP95 cells were a kind gift from Drs. Alan K Meeker and Jun Luo (Johns Hopkins University, Baltimore, Maryland, USA), ^ - supplementation as per supplier instructions, ^^ - phenol red free, FCS – fetal calf serum, CCS – charcoal stripped fetal calf serum.

| Gene target | Supplier | Catalogue ID |
|---|---|---|
| human/mouse control | Dharmacon (Horizon) | D-001810-10-05 |
| human BAG-1 | Dharmacon (Horizon) | L-003871-00-0005 |
| mouse BAG-1 | Dharmacon (Horizon) | L-042153-00-0005 |
| mouse AR-FL | Dharmacon (Horizon) | L-050296-00-0005 |

**Supplementary Table 5: ON-TARGETplus siRNA pools for gene expression knockdown.**

| Guide | sgRNA | Sequence |
|---|---|---|
| BAG-1 g1 | CRISPRevolution EZ RNA-1 (BAG1+33262829) | A*G*G*UCGUGCUUCUCAUUG*C*C* |
| BAG-1 g2 | CRISPRevolution EZ RNA-2 (BAG1+33262809) | C*C*U*GCUGGGAGGUAACAU*G*A* |
| BAG-1 g3 | CRISPRevolution EZ RNA-3 (BAG1+33262794) | C*U*G*GUUCACUGCUGCCCU*G*C* |
| BAG-1 g4 | CRISPRevolution EZRNA-4 (BAG1+33262768) | U*G*A*ACCAGUUGUCCAAGA*C*C* |

**Supplementary Table 6: Synthego BAG-1 sgRNA for BAG-1 CRISPR knockout clones.**

| Pathway | NES | FDR |
|---|---|---|
| HALLMARK_INTERFERON_ALPHA_RESPONSE | 1.85 | 0.002 |
| HALLMARK_INTERFERON_GAMMA_RESPONSE | 1.74 | <0.001 |
| HALLMARK_INFLAMMATORY_RESPONSE | 1.53 | 0.014 |
| HALLMARK_EPITHELIAL_MESENCHYMAL_TRANSITION | 1.50 | 0.016 |
| HALLMARK_TNFA_SIGNALING_VIA_NFKB | 1.44 | 0.024 |
| HALLMARK_APOPTOSIS | 1.42 | 0.045 |
| HALLMARK_MITOTIC_SPINDLE | -1.46 | 0.011 |
| HALLMARK_OXIDATIVE_PHOSPHORYLATION | -1.54 | 0.008 |
| HALLMARK_DNA_REPAIR | -1.59 | 0.008 |
| HALLMARK_MYC_TARGETS_V2 | -2.14 | <0.001 |
| HALLMARK_MYC_TARGETS_V1 | -2.38 | <0.001 |
| HALLMARK_ANDROGEN_RESPONSE | -2.43 | <0.001 |
| HALLMARK_G2M_CHECKPOINT | -2.59 | <0.001 |
| HALLMARK_E2F_TARGETS | -2.96 | <0.001 |

**Supplementary Table 7: Cellular pathways de-enriched and enriched by Thio-2 treatment in LNCaP cells.** Normalized enrichment score (NES) for cellular pathways de-enriched and enriched by Thio-2 treatment with a false discovery rate (FDR) < 0.05 are shown.

| Uniprot ID | PDB ID | Chain | Complex | Organism | Res | Method | Released date |
|---|---|---|---|---|---|---|---|
| Q99933 | 1HX1 | B | HSPA8 | HUMAN | 1.90000 | X-Ray | 07/03/2001 |
| Q99933 | 3FZF | B | HSPA8 | HUMAN | 2.00000 | X-Ray | 17/03/2009 |
| Q99933 | 3FZH | B | HSPA8 | HUMAN | 1.90000 | X-Ray | 17/03/2009 |
| Q99933 | 3FZK | B | HSPA8 | HUMAN | 2.00000 | X-Ray | 17/03/2009 |
| Q99933 | 3FZL | B | HSPA8 | HUMAN | 2.00000 | X-Ray | 17/03/2009 |
| Q99933 | 3FZM | B | HSPA8 | HUMAN | 2.10000 | X-Ray | 17/03/2009 |
| Q99933 | 3LDQ | B | HSPA8 | HUMAN | 1.90000 | X-Ray | 26/01/2011 |
| Q99933 | 3M3Z | B | HSPA8 | HUMAN | 2.00000 | X-Ray | 26/01/2011 |
| Q99933 | 5AQF | D | HSPA8 | HUMAN | 1.88000 | X-Ray | 05/10/2016 |
| Q99933 | 5AQF | B | HSPA8 | HUMAN | 1.88000 | X-Ray | 05/10/2016 |
| Q99933 | 5AQG | F | HSPA8 | HUMAN | 2.24000 | X-Ray | 05/10/2016 |
| Q99933 | 5AQG | D | HSPA8 | HUMAN | 2.24000 | X-Ray | 05/10/2016 |
| Q99933 | 5AQG | B | HSPA8 | HUMAN | 2.24000 | X-Ray | 05/10/2016 |
| Q99933 | 5AQH | B | HSPA8 | HUMAN | 2.00000 | X-Ray | 05/10/2016 |
| Q99933 | 5AQI | D | HSPA8 | HUMAN | 1.98000 | X-Ray | 05/10/2016 |
| Q99933 | 5AQI | B | HSPA8 | HUMAN | 1.98000 | X-Ray | 05/10/2016 |
| Q99933 | 5AQJ | F | HSPA8 | HUMAN | 1.96000 | X-Ray | 05/10/2016 |
| Q99933 | 5AQJ | D | HSPA8 | HUMAN | 1.96000 | X-Ray | 05/10/2016 |
| Q99933 | 5AQJ | B | HSPA8 | HUMAN | 1.96000 | X-Ray | 05/10/2016 |
| Q99933 | 5AQK | B | HSPA8 | HUMAN | 2.09000 | X-Ray | 05/10/2016 |
| Q99933 | 5AQL | D | HSPA8 | HUMAN | 1.69000 | X-Ray | 05/10/2016 |
| Q99933 | 5AQL | B | HSPA8 | HUMAN | 1.69000 | X-Ray | 05/10/2016 |
| Q99933 | 5AQM | D | HSPA8 | HUMAN | 1.63000 | X-Ray | 05/10/2016 |
| Q99933 | 5AQM | B | HSPA8 | HUMAN | 1.63000 | X-Ray | 05/10/2016 |
| Q99933 | 5AQN | B | HSPA8 | HUMAN | 2.45000 | X-Ray | 05/10/2016 |
| Q99933 | 5AQN | F | HSPA8 | HUMAN | 2.45000 | X-Ray | 05/10/2016 |
| Q99933 | 5AQN | D | HSPA8 | HUMAN | 2.45000 | X-Ray | 05/10/2016 |
| Q99933 | 5AQO | F | HSPA8 | HUMAN | 2.12000 | X-Ray | 05/10/2016 |
| Q99933 | 5AQO | D | HSPA8 | HUMAN | 2.12000 | X-Ray | 05/10/2016 |
| Q99933 | 5AQO | B | HSPA8 | HUMAN | 2.12000 | X-Ray | 05/10/2016 |
| Q99933 | 5AQP | F | HSPA8 | HUMAN | 2.08000 | X-Ray | 05/10/2016 |
| Q99933 | 5AQP | D | HSPA8 | HUMAN | 2.08000 | X-Ray | 05/10/2016 |
| Q99933 | 5AQP | B | HSPA8 | HUMAN | 2.08000 | X-Ray | 05/10/2016 |
| Q99933 | 5AQQ | F | HSPA8 | HUMAN | 2.72000 | X-Ray | 05/10/2016 |
| Q99933 | 5AQQ | D | HSPA8 | HUMAN | 2.72000 | X-Ray | 05/10/2016 |
| Q99933 | 5AQQ | B | HSPA8 | HUMAN | 2.72000 | X-Ray | 05/10/2016 |
| Q99933 | 5AQR | F | HSPA8 | HUMAN | 1.91000 | X-Ray | 05/10/2016 |
| Q99933 | 5AQR | D | HSPA8 | HUMAN | 1.91000 | X-Ray | 05/10/2016 |
| Q99933 | 5AQR | B | HSPA8 | HUMAN | 1.91000 | X-Ray | 05/10/2016 |
| Q99933 | 5AQS | D | HSPA8 | HUMAN | 2.00000 | X-Ray | 05/10/2016 |
| Q99933 | 5AQS | B | HSPA8 | HUMAN | 2.00000 | X-Ray | 05/10/2016 |
| Q99933 | 5AQT | B | HSPA8 | HUMAN | 1.90000 | X-Ray | 05/10/2016 |
| Q99933 | 5AQU | B | HSPA8 | HUMAN | 1.92000 | X-Ray | 05/10/2016 |
| Q99933 | 5AQV | B | HSPA8 | HUMAN | 1.75000 | X-Ray | 05/10/2016 |

**Supplementary Table 8: List of the 44 3D structures of HSC70 (HSPA8) in complex with the BAG domain of BAG-1 used in comparative structural analysis**
ID – Identifier, PDB – protein data bank, Res – resolution, X-Ray – X-Ray diffraction.

| Clinical characteristics | |
|---|---|
| **Archival (castration-sensitive) biopsy (N=43)** | |
| **Histology (N, %)** | |
| Adenocarcinoma | 43, 100% |
| **Gleason score (N, %)** | |
| <7 | 3, 7% |
| 7 | 2, 5% |
| >7 | 35, 81% |
| NR | 3, 7% |
| **Metastatic at diagnosis (N, %)** | |
| M0 | 18, 42% |
| M1 | 16, 37% |
| NR | 9, 21% |
| **Treatment intent (N, %)** | |
| Radical | 18, 42% |
| Palliative | 25, 58% |
| **Metastatic castration-resistant biopsy (N=67; 43 paired)** | |
| **Biopsy site (N, %)** | |
| Bone | 31, 46% |
| Lymph node | 18, 27% |
| Liver | 7, 10% |
| TURP | 5, 8% |
| Other | 6, 9% |
| **NHT prior to biopsy** | |
| No | 27, 40% |
| Yes | 40, 60% |

**Supplementary table 9: Clinical characteristics of ICR/RMH patient IHC cohort at time of diagnostic (archival) castration-sensitive and metastatic castration-resistant tissue biopsies.**

N – number, NR – not recorded, NHT – novel hormonal therapies

| Clinical characteristics | |
|---|---|
| **Radical prostatectomies (N=30)** | |
| **Histology (N, %)**<br>Adenocarcinoma | 30, 100% |
| **Mean Age at diagnosis** | 63.0 (7.6) |
| **Mean Gleason score (SD)** | 6.7 (1.8) |
| **Metastases (N=30)** | |
| **Mean Age at diagnosis** | 62.0 (9.0) |
| **Mean Gleason score (SD)** | 8.2 (1.01) |

**Supplementary table 10: Clinical characteristics of UW/FHCRC patient IHC cohort.**

N – number, SD – standard deviation

| **Clinical characteristics** | |
|---|---|
| **Prior to starting NHT (N=50)** | |
| **Age (years), Mean (SD)** | 67.3 (8.2) |
| **Metastatic (N, %)** | |
| No | 7, 14% |
| Yes | 43, 86% |
| **NHT (N, %)** | |
| Abiraterone | 43, 86% |
| Enzalutamide | 7, 14% |
| **Previous taxane chemotherapy (N, %)** | |
| No | 0, 0% |
| Yes | 50, 100% |

**Supplementary table 11: Baseline characteristics of ICR/RMH patient IHC cohort prior to starting novel hormonal therapy following taxane chemotherapy.**

NHT – novel hormonal therapies, N – number, SD – standard deviation,
