## Supplemental Figures for "Targeting the BAG-1 family of co-chaperones in lethal prostate cancer"

#### Supplementary Figure 1

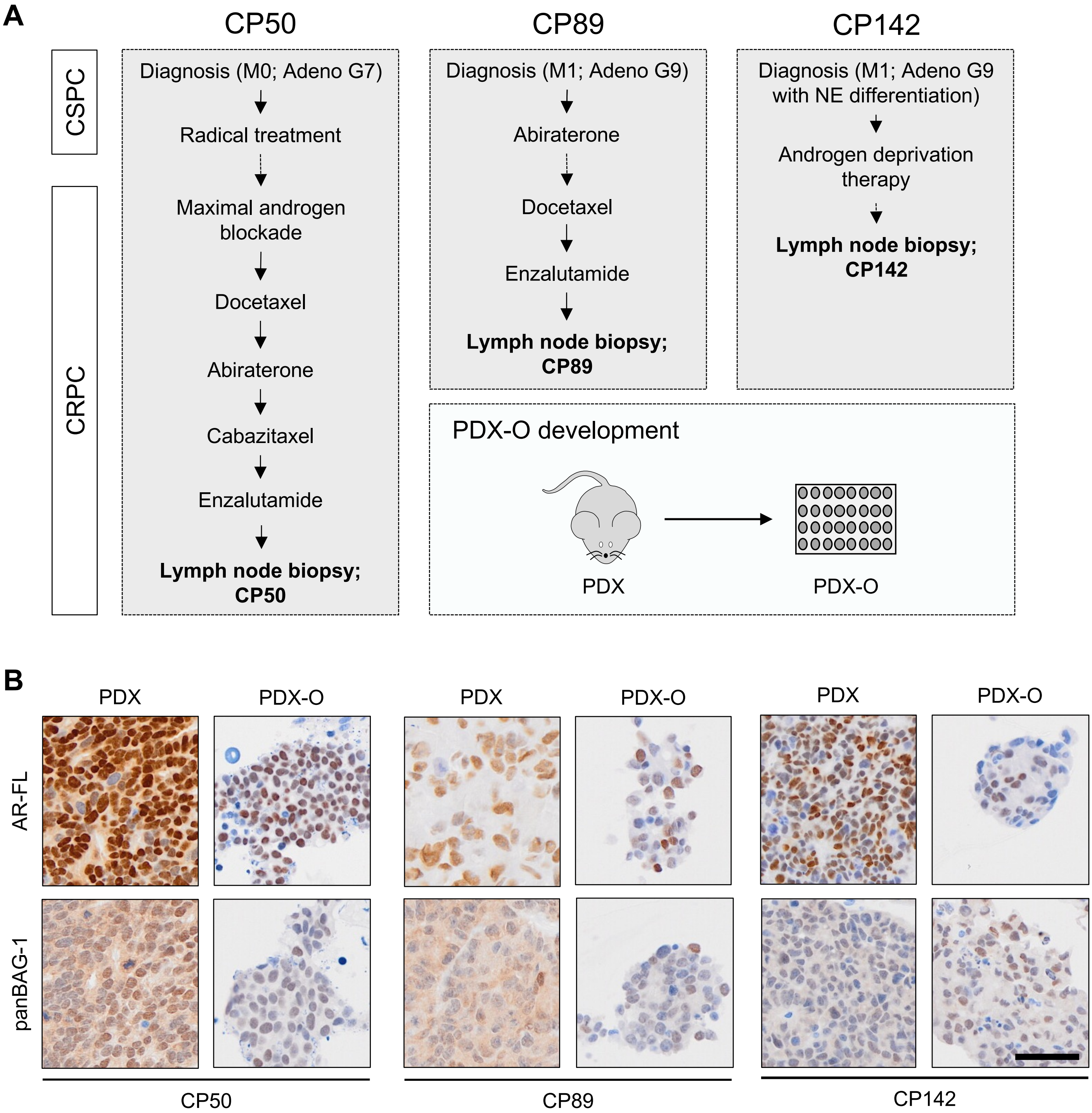

#### Supplementary Figure 2

**A**

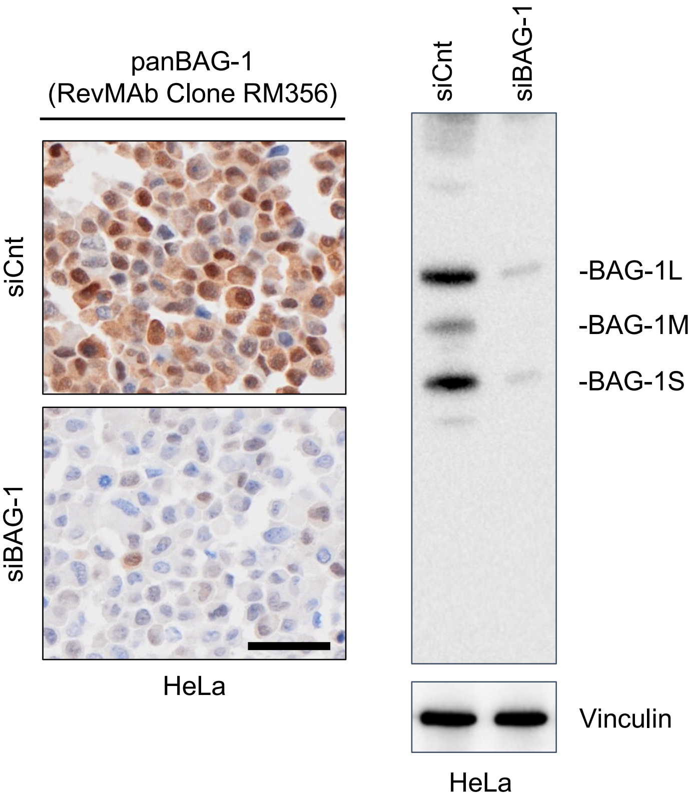

### Supplementary Figure 3

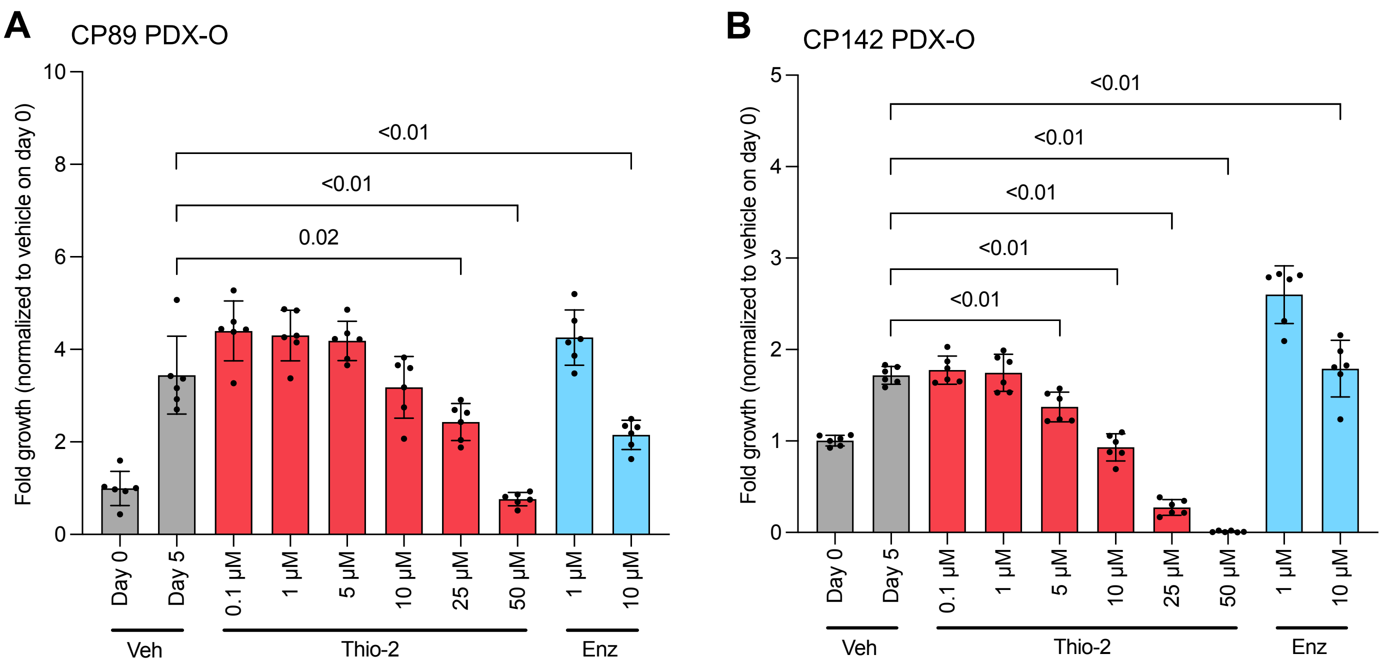

### Supplementary Figure 4

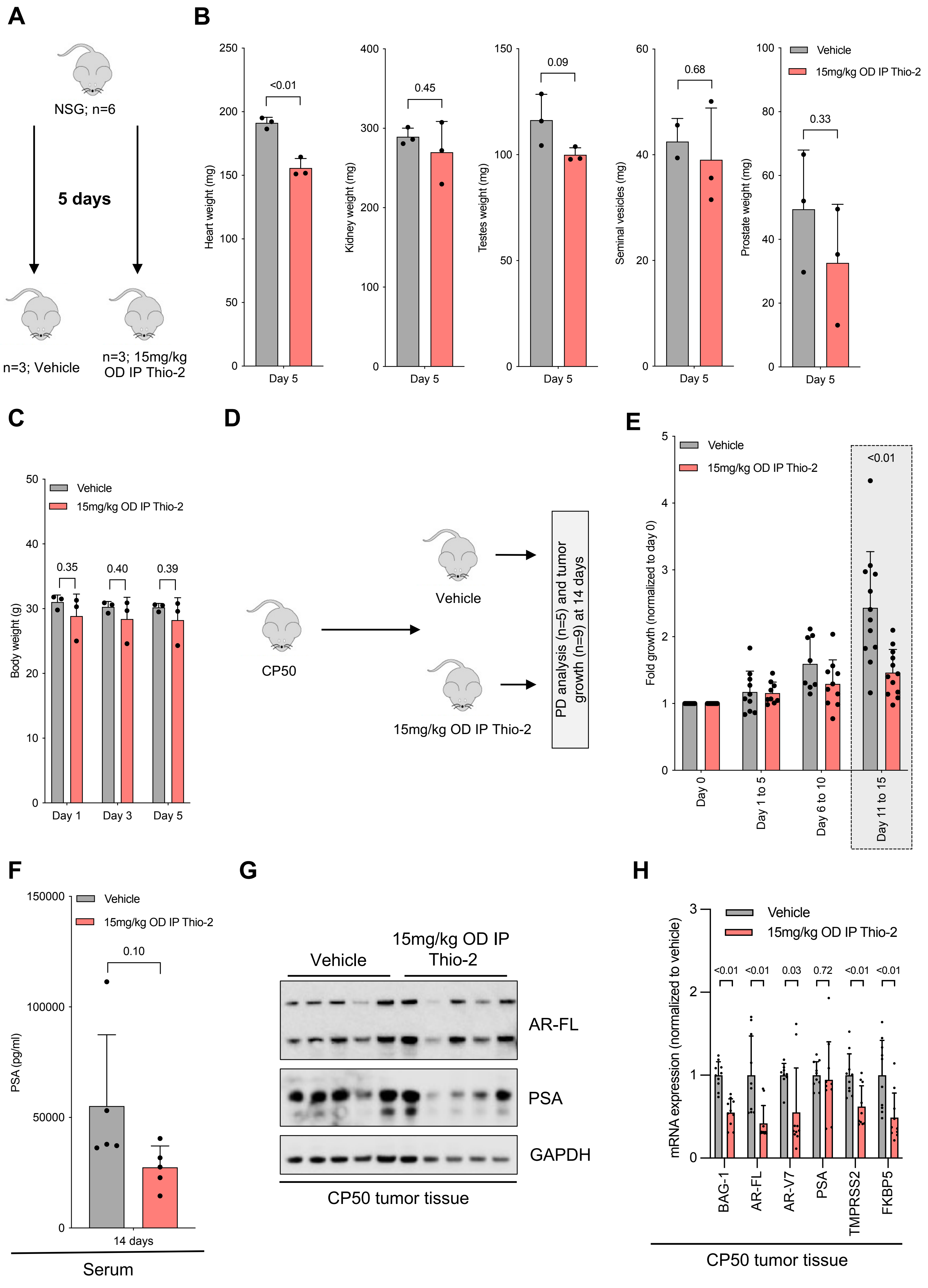

### Supplementary Figure 5

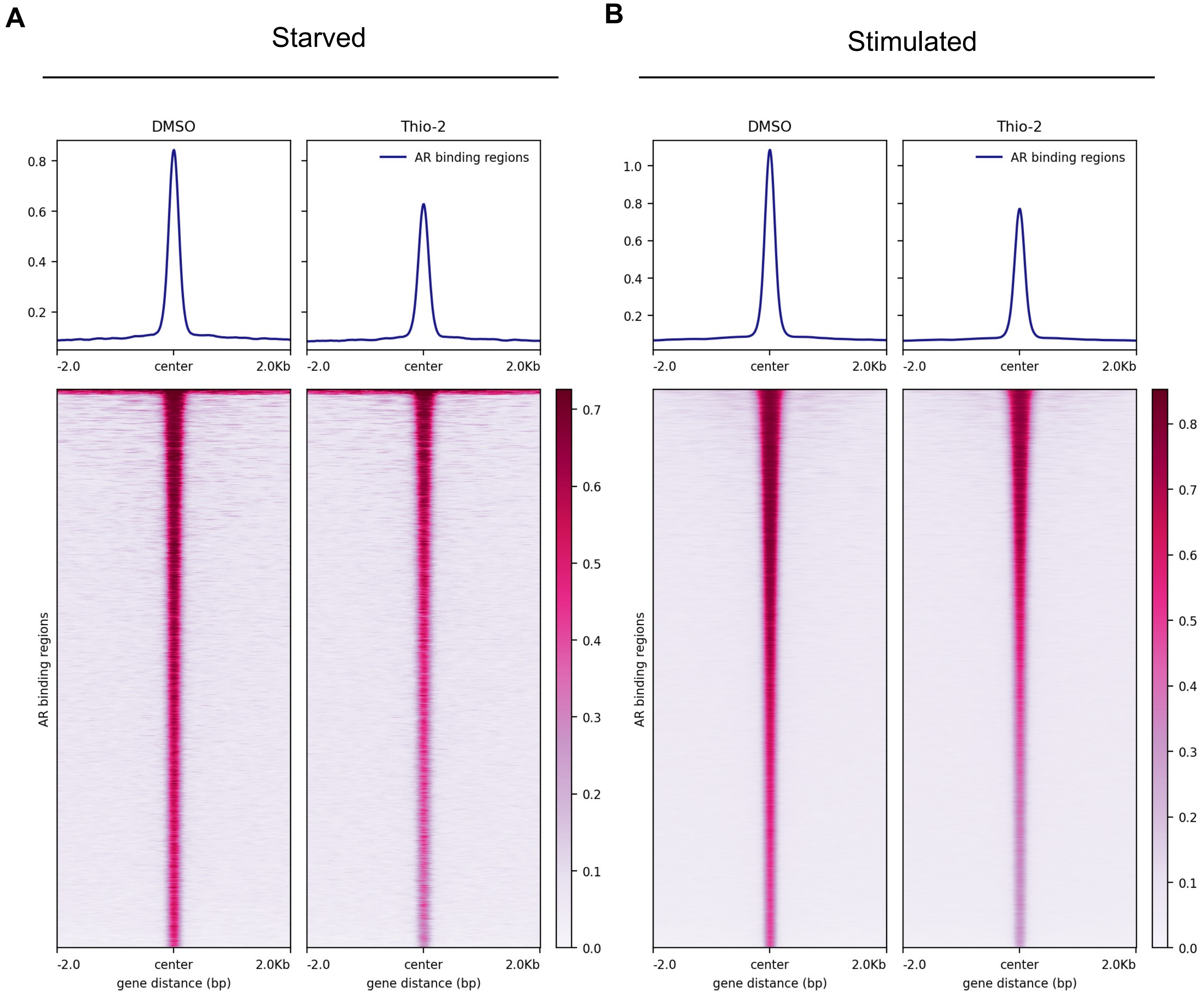

#### Supplementary Figure 6

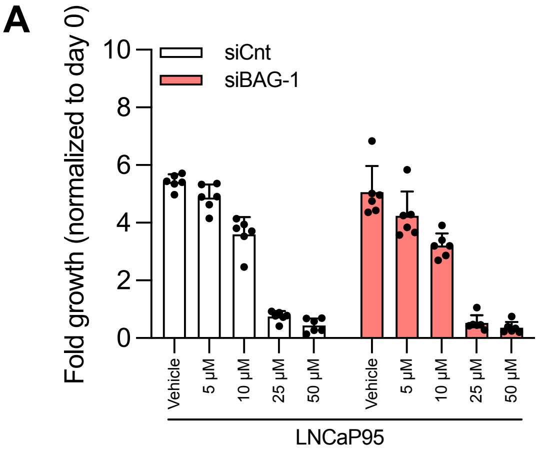


| | | P value siCnt | | P value siBAG-1 |
|---|---|---|---|---|
| Growth | 0 vs 5µM | 0.03 | ns | ns |
| | 0 vs 10µM | <0.01 | | <0.01 |
| | 0 vs 25µM | <0.01 | | <0.01 |
| | 0 vs 50µM | <0.01 | | <0.01 |

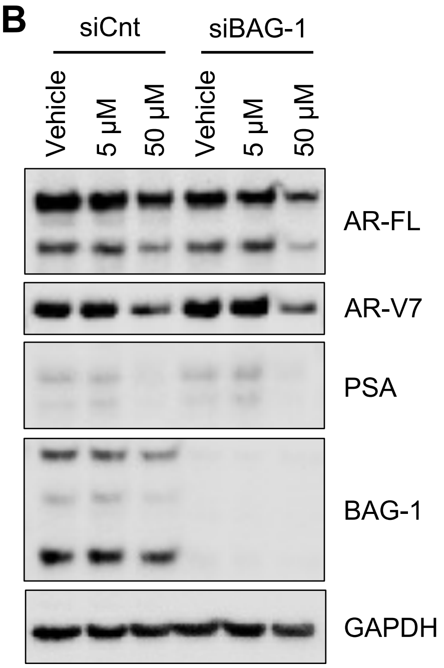


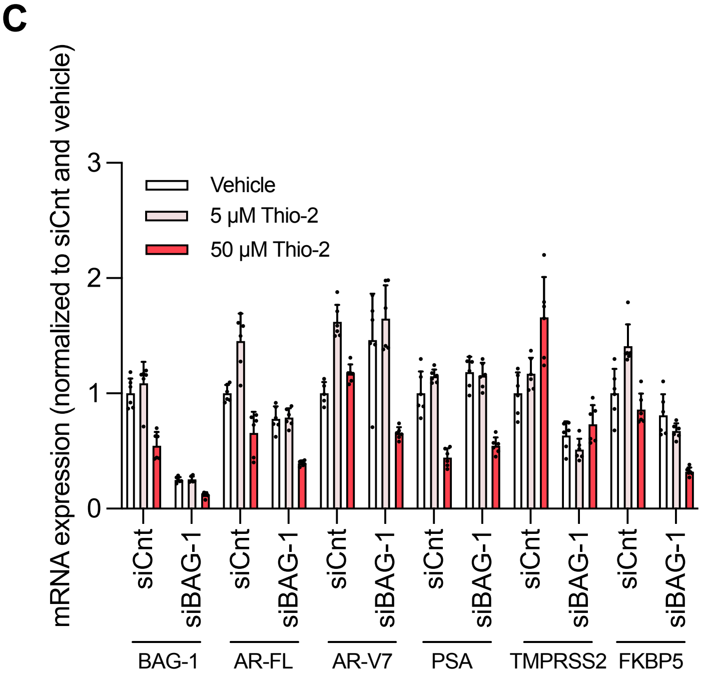


| | | P value siCnt | | P value siBAG-1 |
|---|---|---|---|---|
| BAG-1 | 0 vs 5µM | ns | <0.01 | ns |
| | 0 vs 50µM | <0.01 | | <0.01 |
| AR-FL | 0 vs 5µM | <0.01 | <0.01 | ns |
| | 0 vs 50µM | <0.01 | | <0.01 |
| AR-V7 | 0 vs 5µM | <0.01 | 0.04 | ns |
| | 0 vs 50µM | 0.01 | | <0.01 |
| PSA | 0 vs 5µM | ns | ns | ns |
| | 0 vs 50µM | <0.01 | | <0.01 |
| TMPRSS2 | 0 vs 5µM | ns | <0.01 | ns |
| | 0 vs 50µM | <0.01 | | ns |
| FKBP5 | 0 vs 5µM | <0.01 | ns | ns |
| | 0 vs 50µM | ns | | <0.01 |

### Supplementary Figure 7

**A**

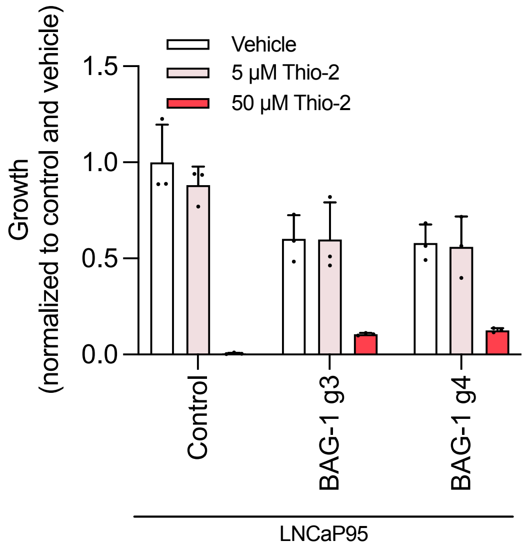


| | | P value | | | | |
|---|---|---|---|---|---|---|
| | | Control | | BAG-1 g3 | | BAG-1 g4 |
| Growth | 0 vs 5µM | ns | 0.04 | ns | 0.03 | ns |
| | 0 vs 50µM | <0.01 | | <0.01 | | <0.01 |

**B**

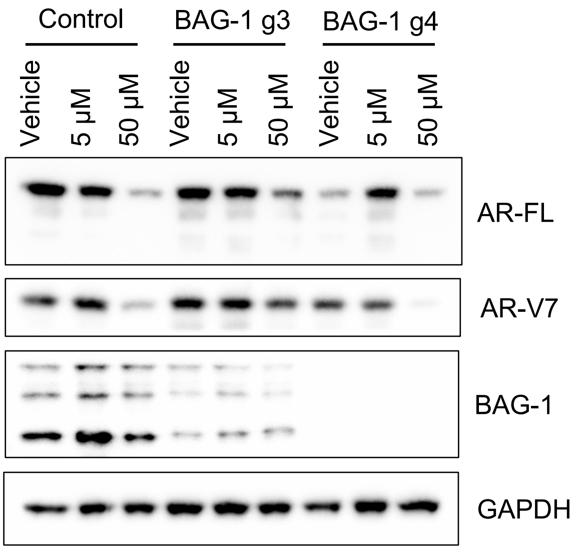


**C**

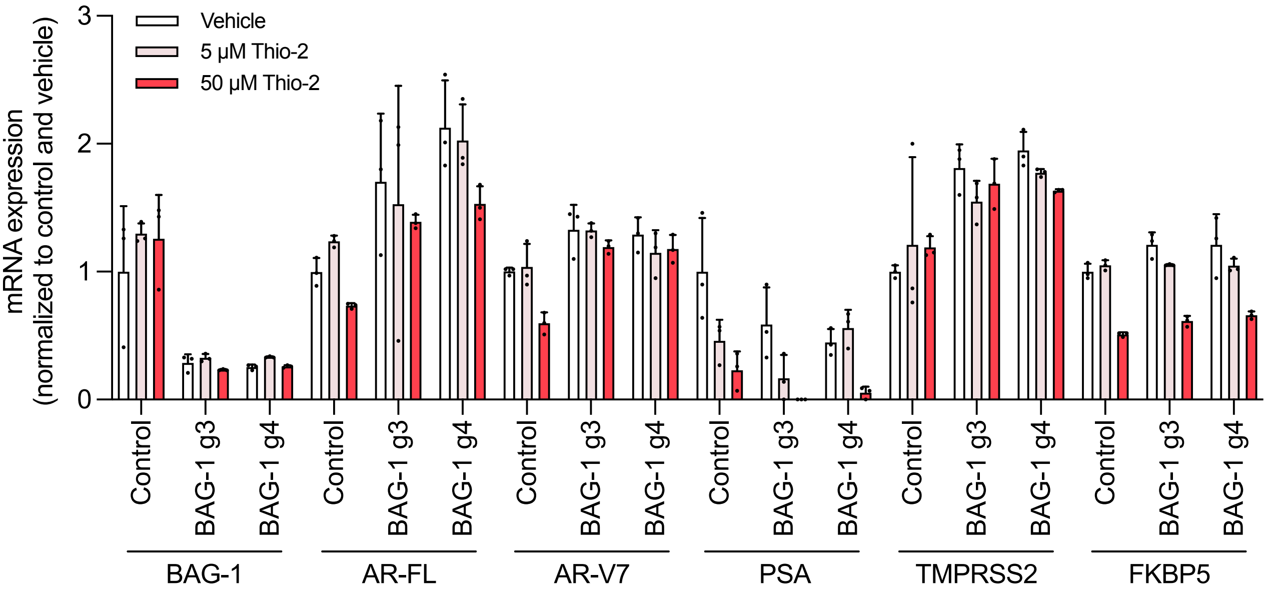


| | | P value | | | | |
|---|---|---|---|---|---|---|
| | | Control | | BAG-1 g3 | | BAG-1 g4 |
| BAG-1 | 0 vs 5µM | ns | ns | ns | ns | <0.01 |
| | 0 vs 50µM | ns | | ns | | ns |
| AR-FL | 0 vs 5µM | 0.03 | ns | ns | <0.01 | ns |
| | 0 vs 50µM | 0.02 | | ns | | ns |
| AR-V7 | 0 vs 5µM | ns | 0.05 | ns | 0.02 | ns |
| | 0 vs 50µM | <0.01 | | ns | | ns |

| | | P value | | | | |
|---|---|---|---|---|---|---|
| | | Control | | BAG-1 g3 | | BAG-1 g4 |
| PSA | 0 vs 5µM | ns | ns | ns | ns | ns |
| | 0 vs 50µM | 0.04 | | 0.03 | | <0.01 |
| TMPRSS2 | 0 vs 5µM | ns | <0.01 | ns | <0.01 | ns |
| | 0 vs 50µM | 0.03 | | ns | | 0.02 |
| FKBP5 | 0 vs 5µM | ns | 0.04 | ns | ns | ns |
| | 0 vs 50µM | <0.01 | | <0.01 | | 0.02 |

**Supplementary Figure 8**

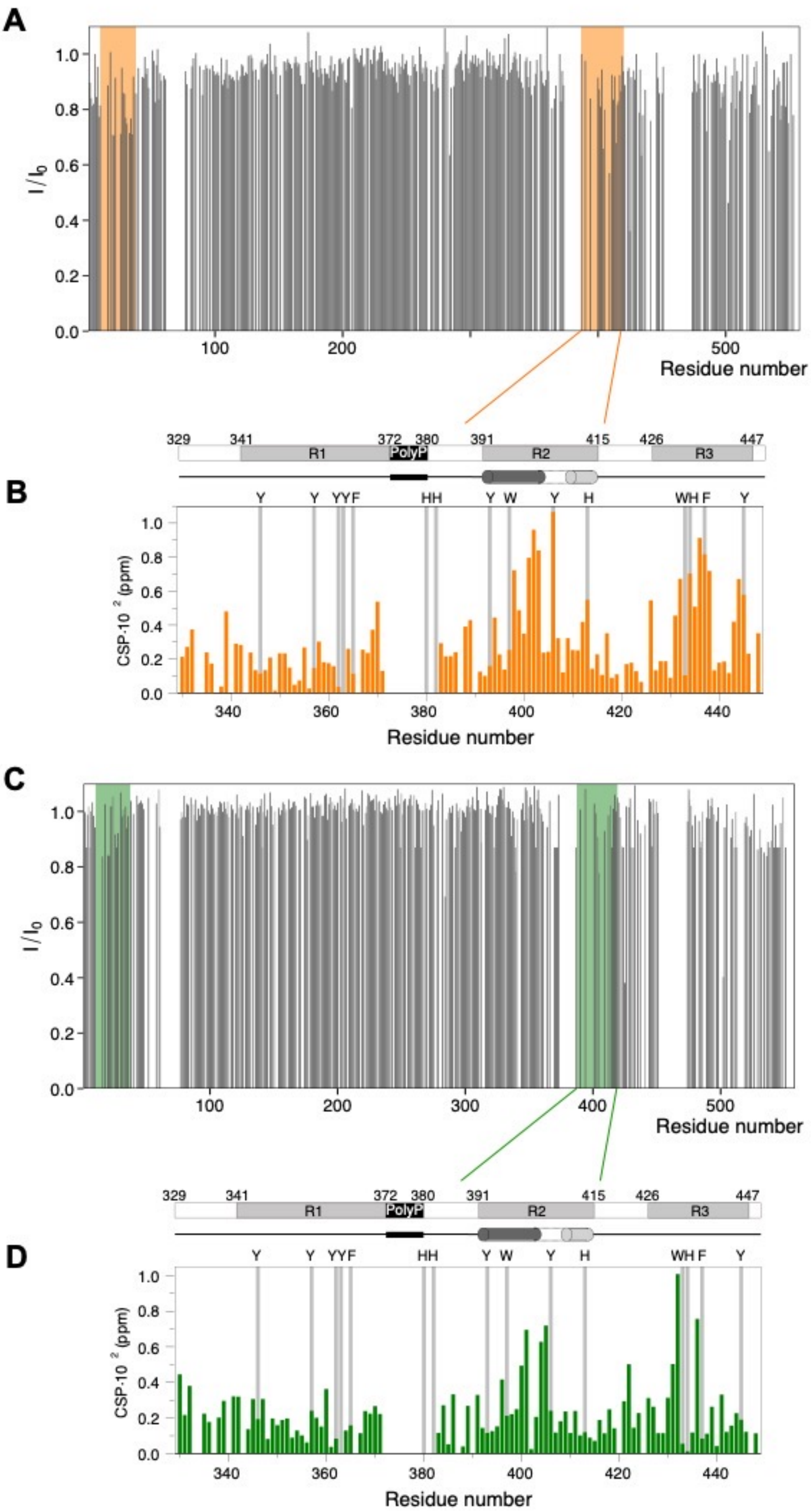

### Supplementary Figure 9

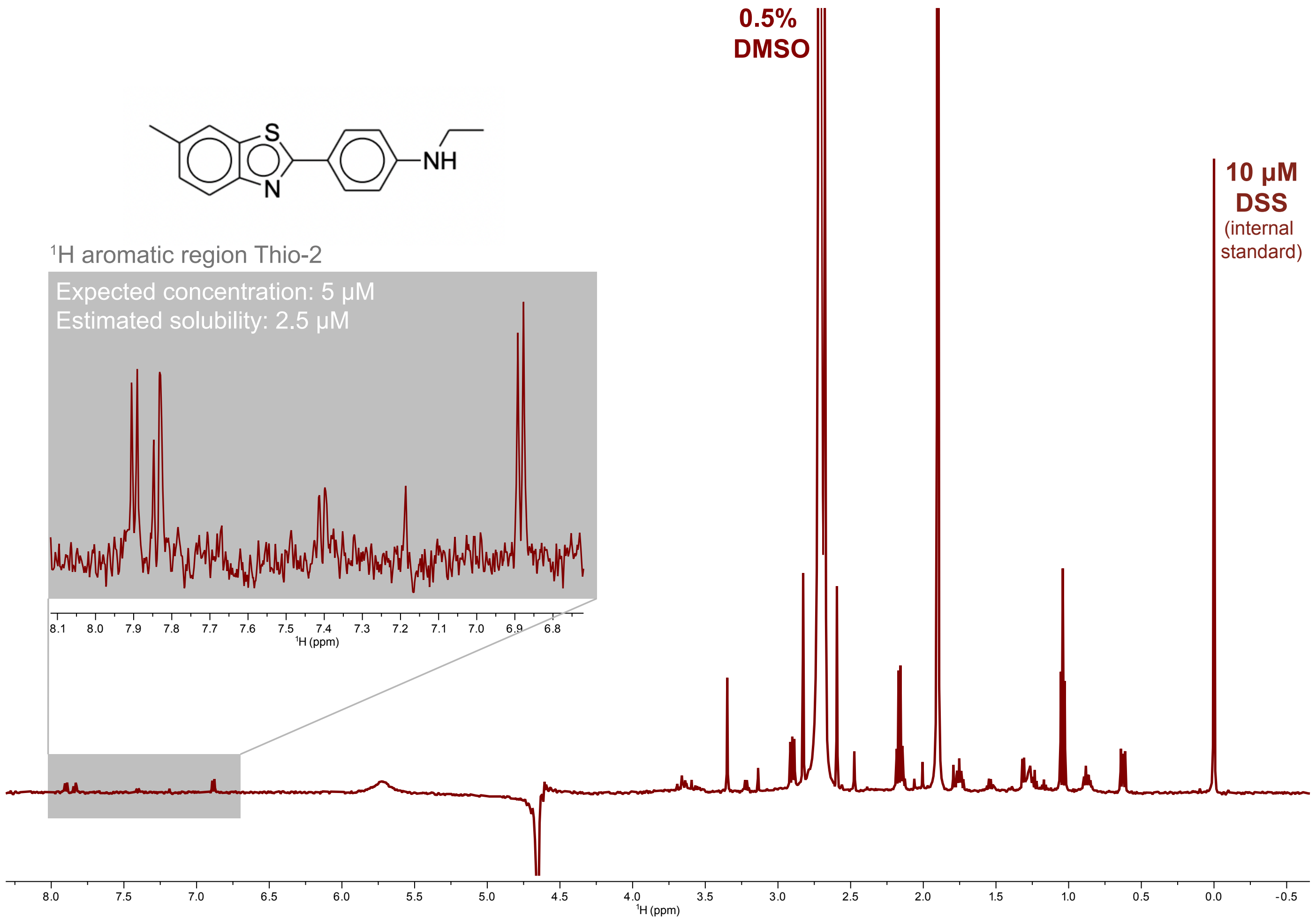

### Supplementary Figure 10

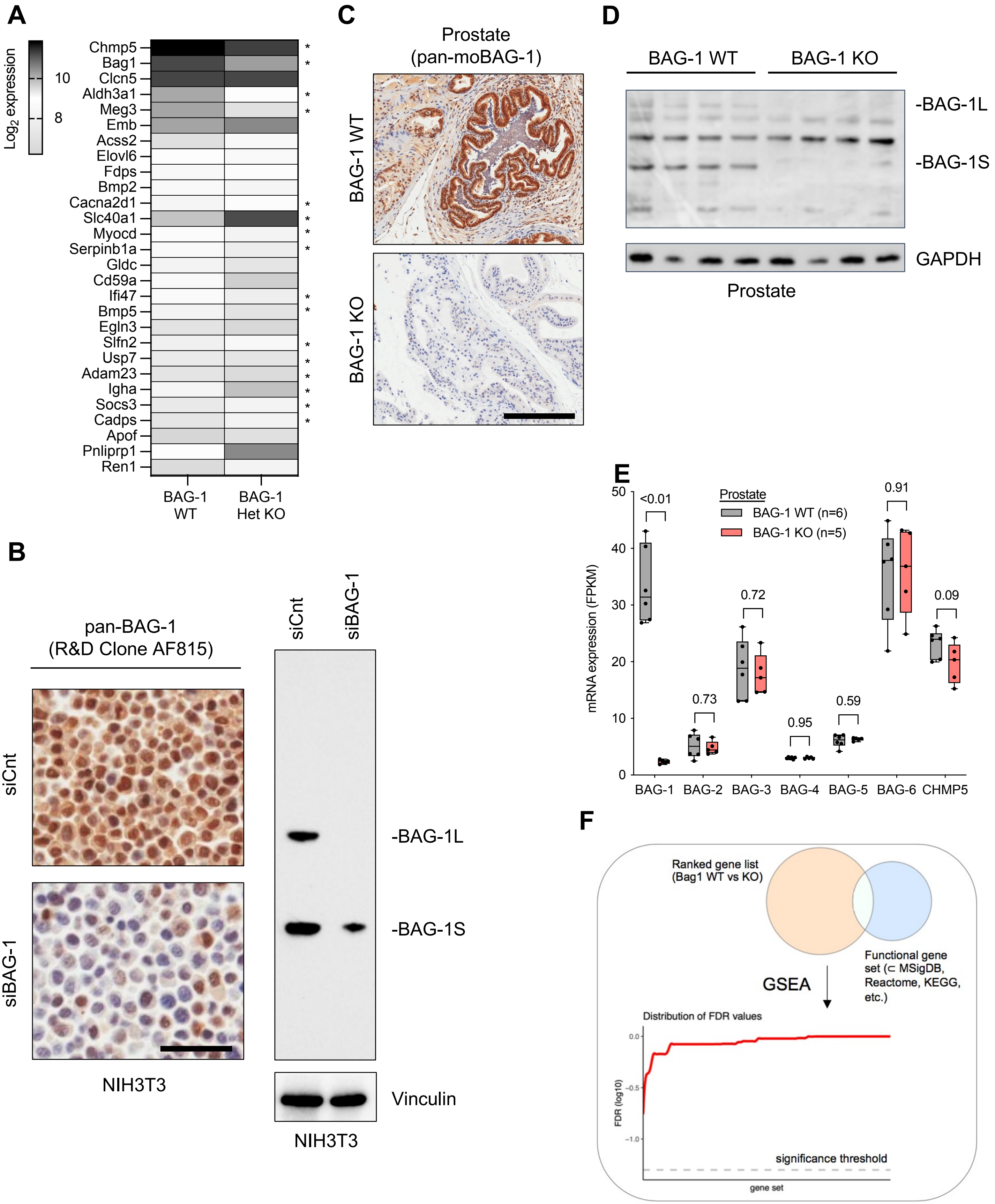

### Supplementary Figure 11

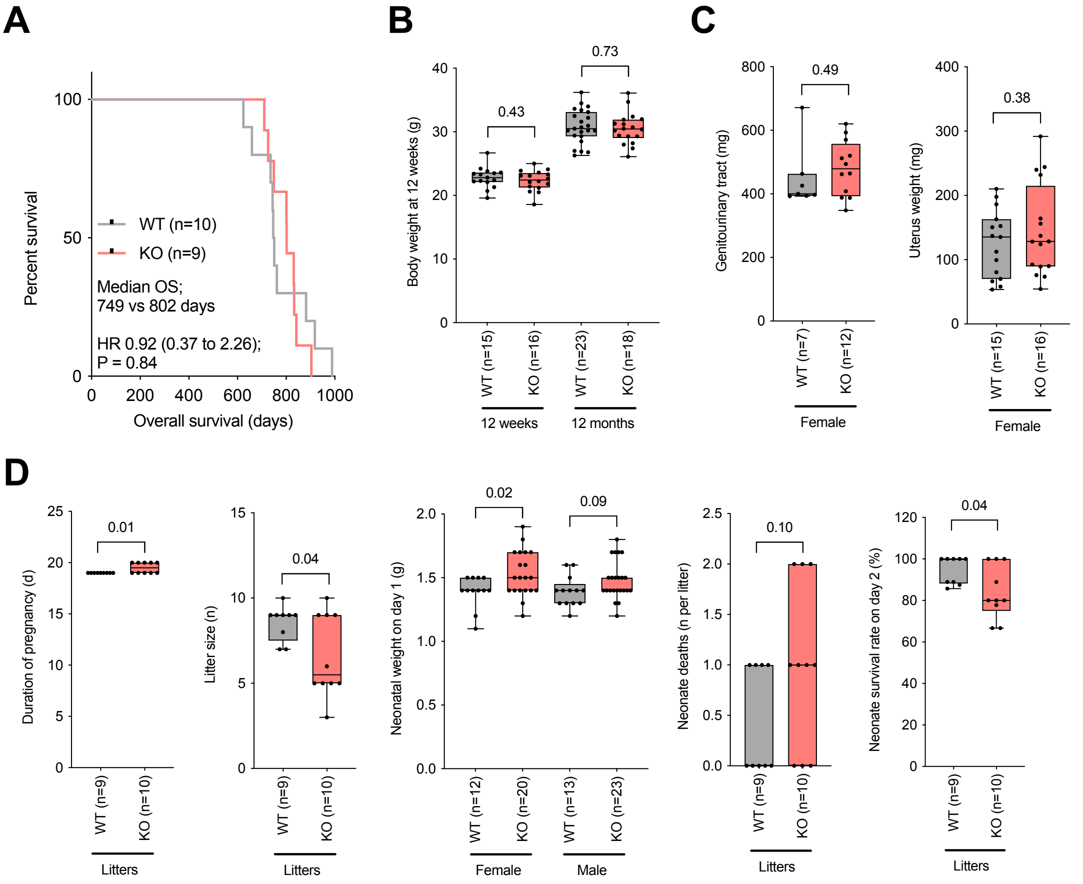

**Supplementary Figure 12**

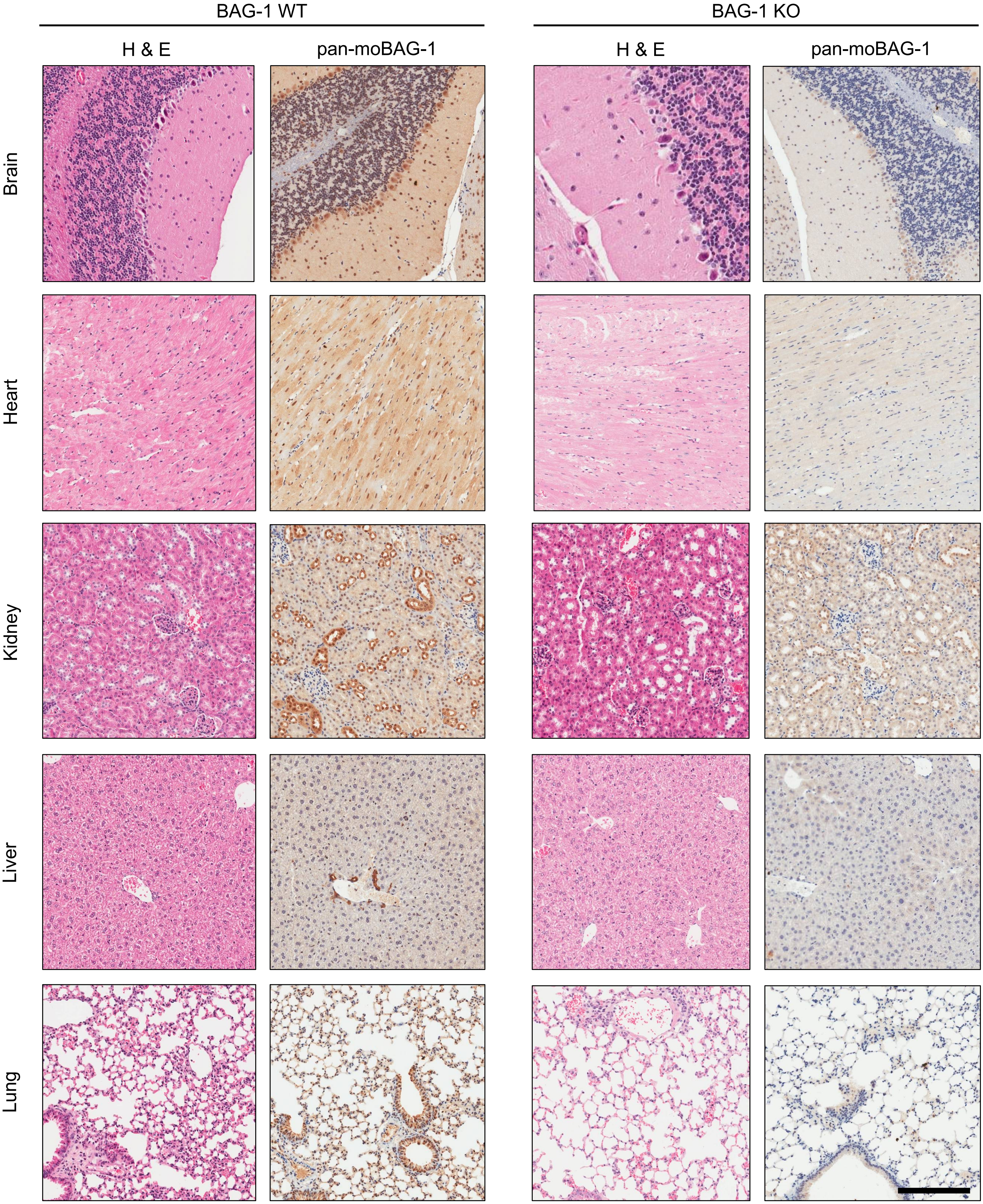

### Supplementary Figure 13

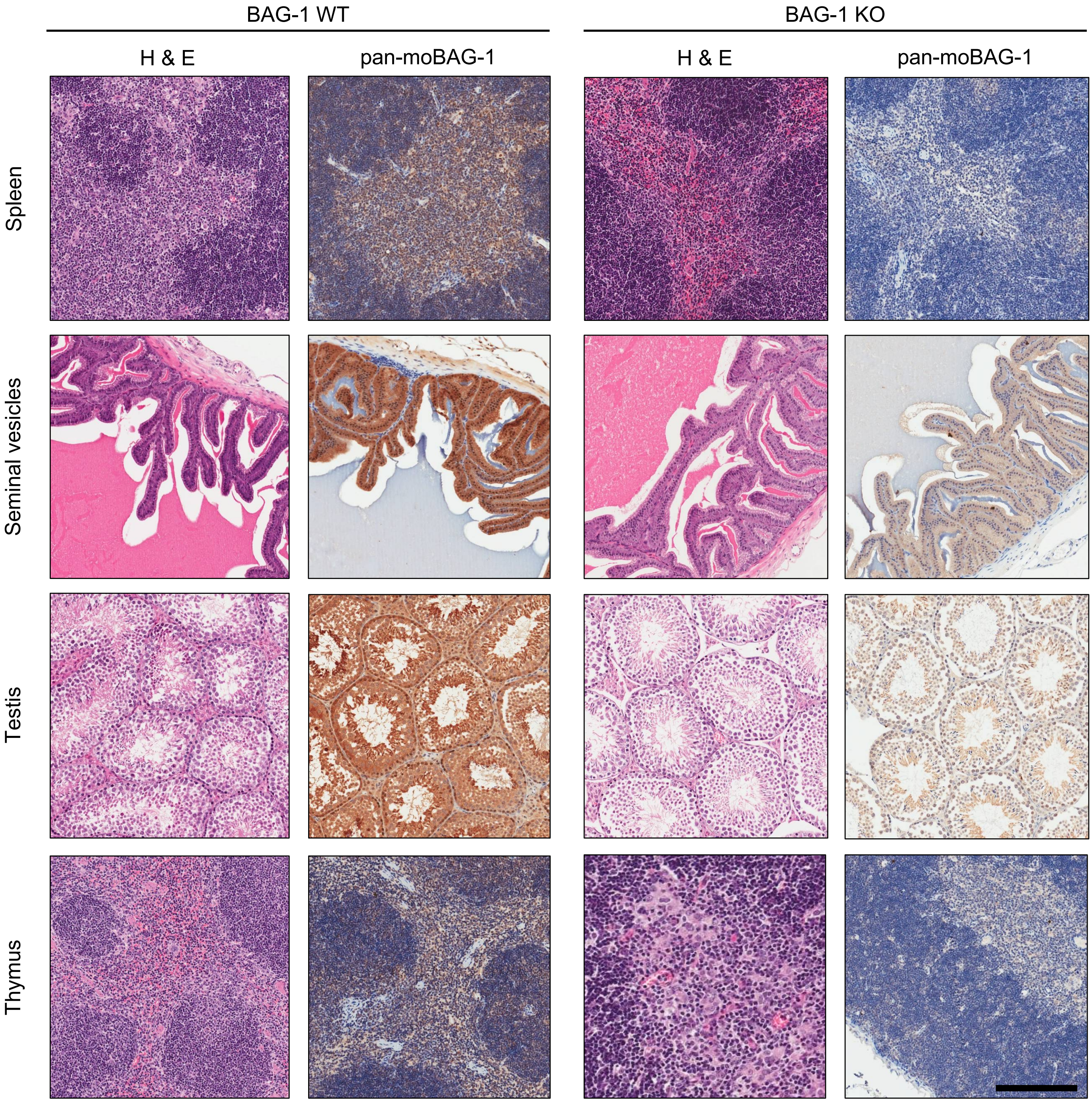

### Supplementary Figure 14

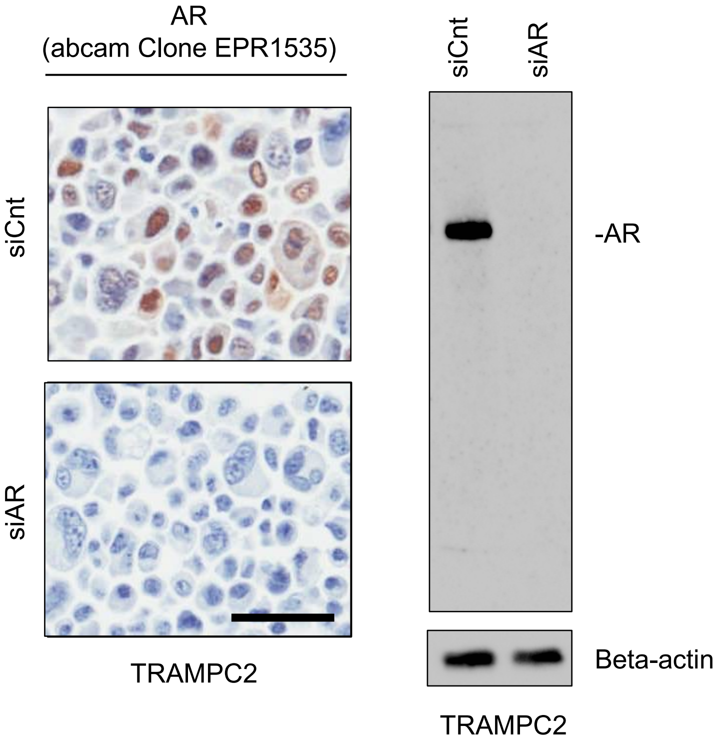

### Supplementary Figure 15

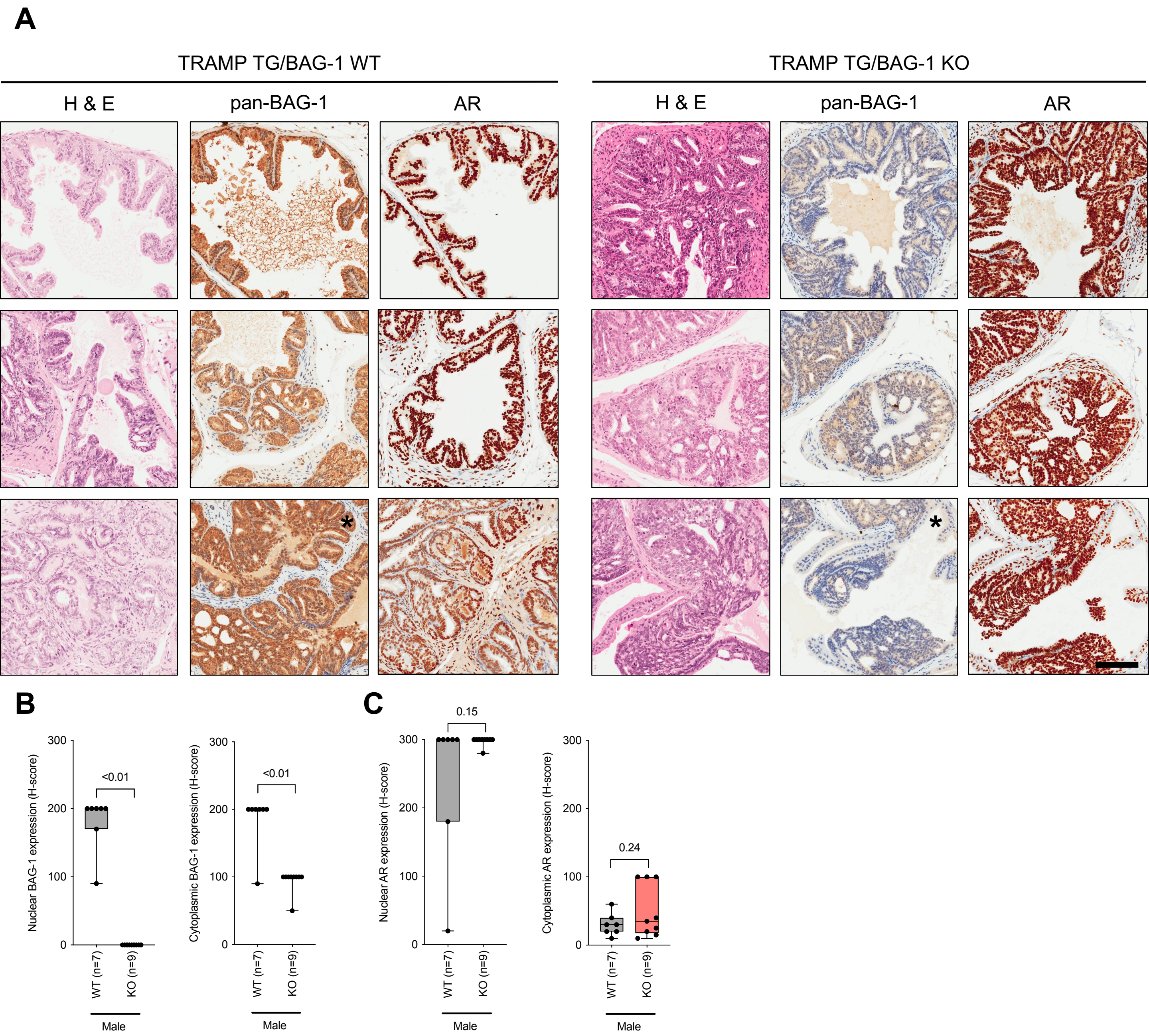

### Supplementary Figure 16

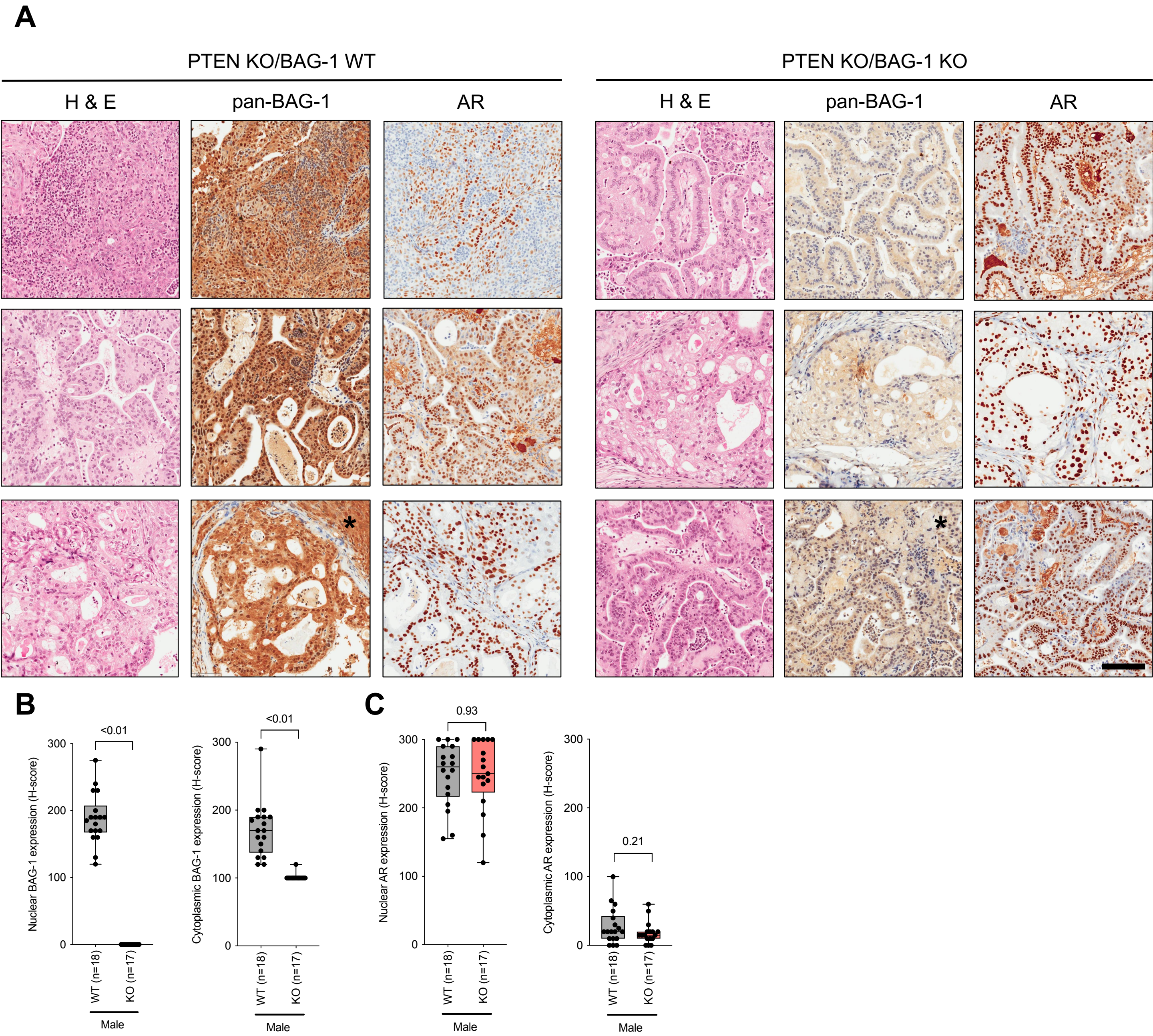

#### Supplementary Figure 17

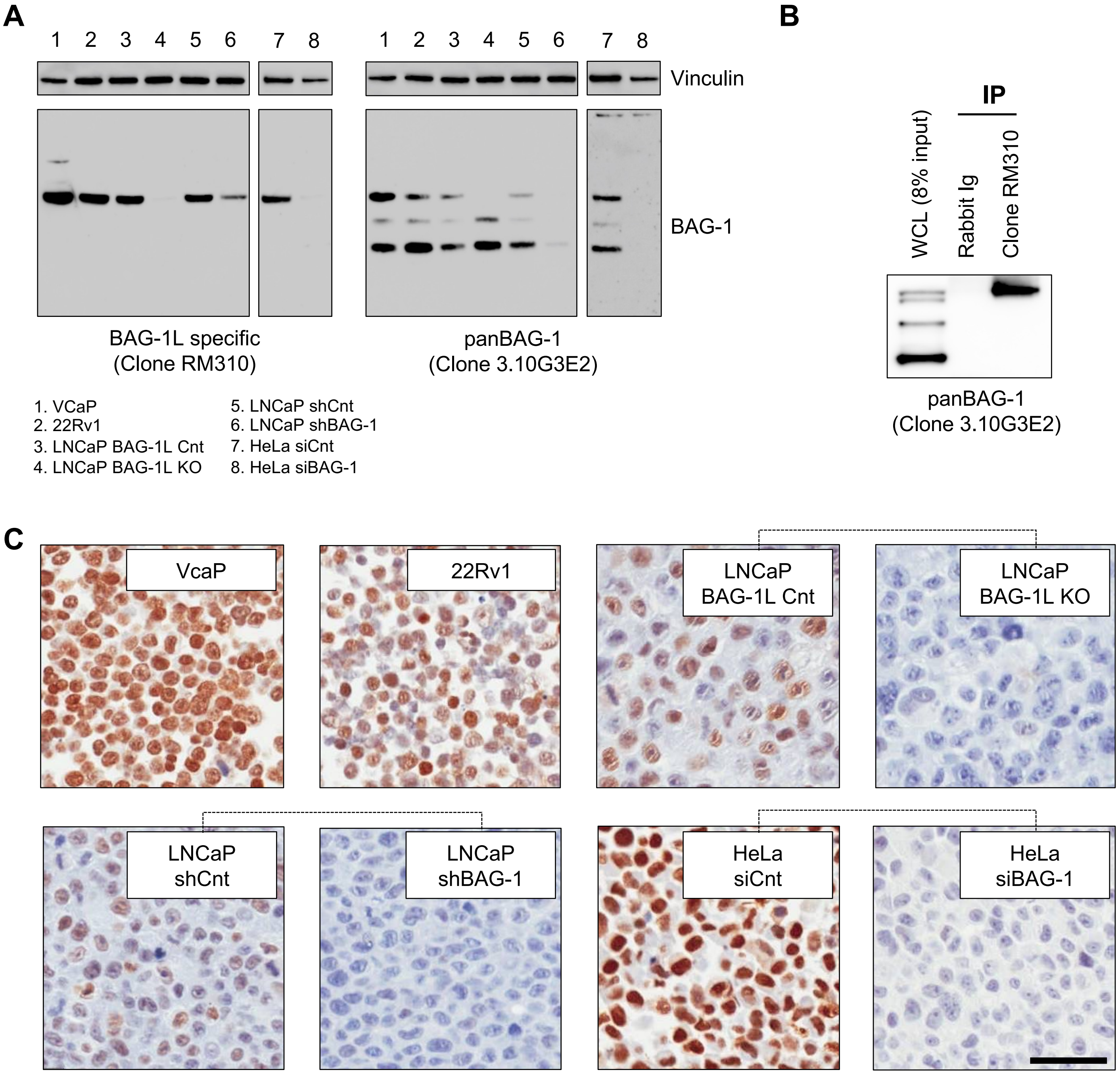

#### Supplementary Figure 18

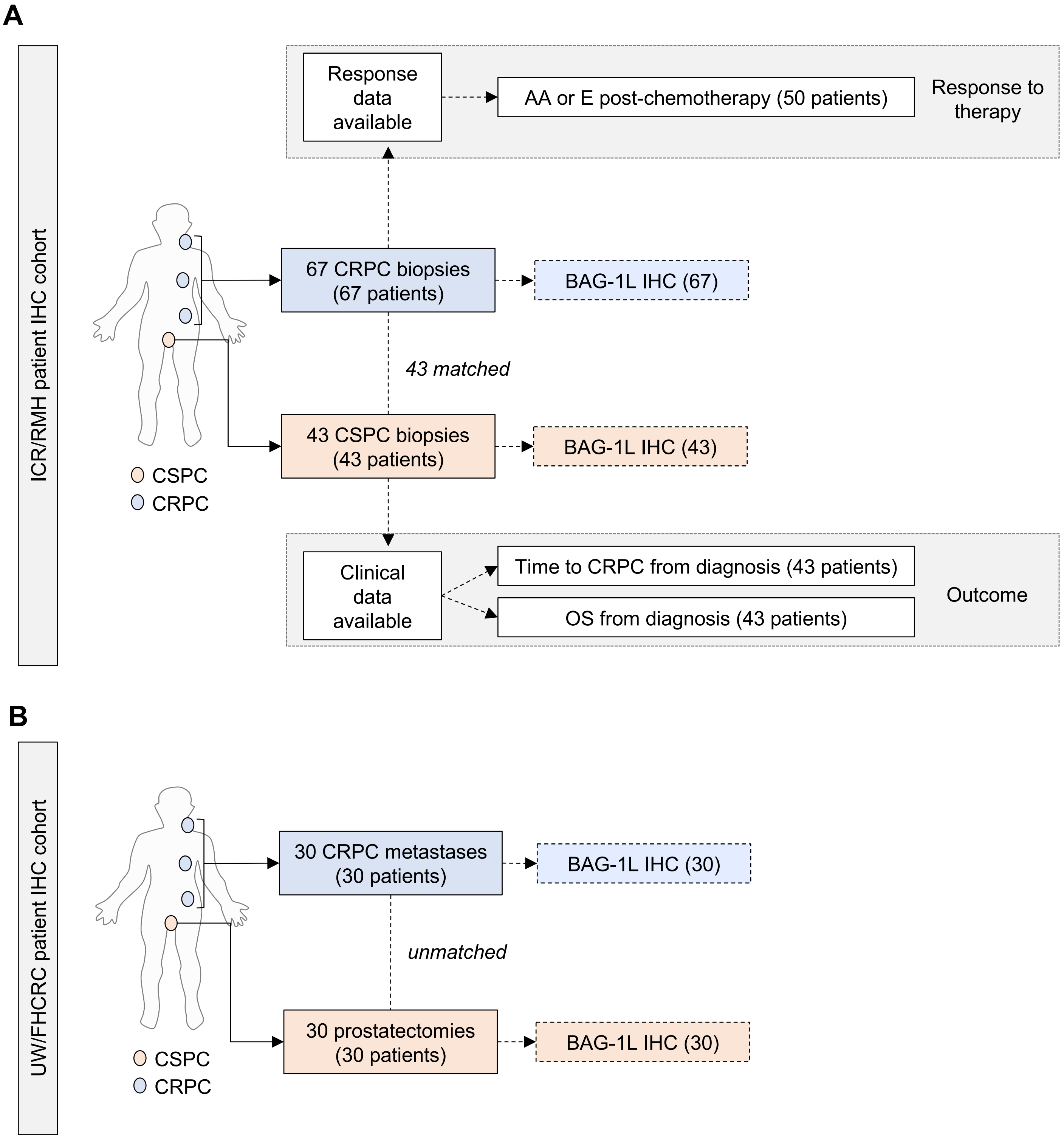

### Supplementary Figure 19

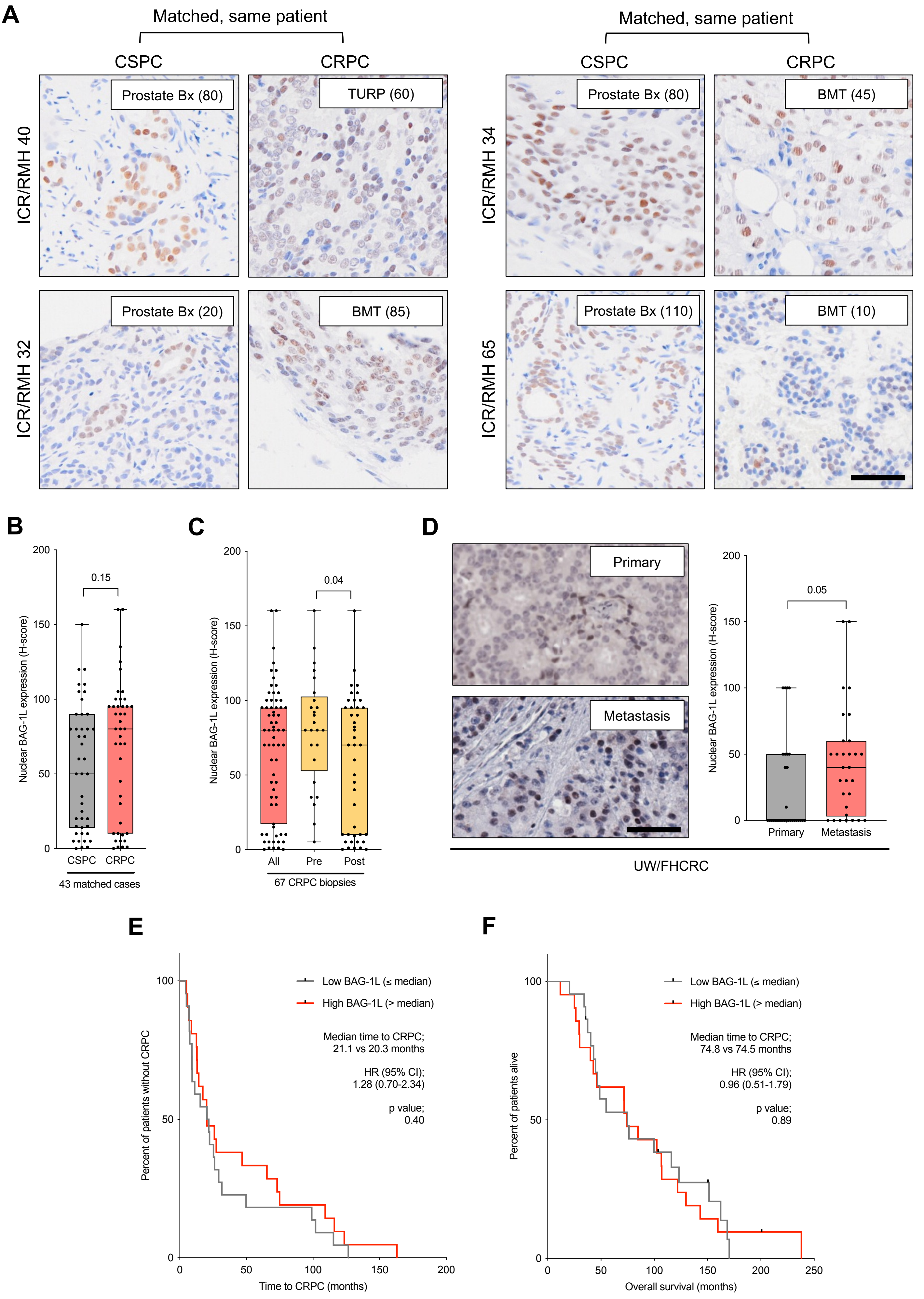

### Supplementary Figure 20

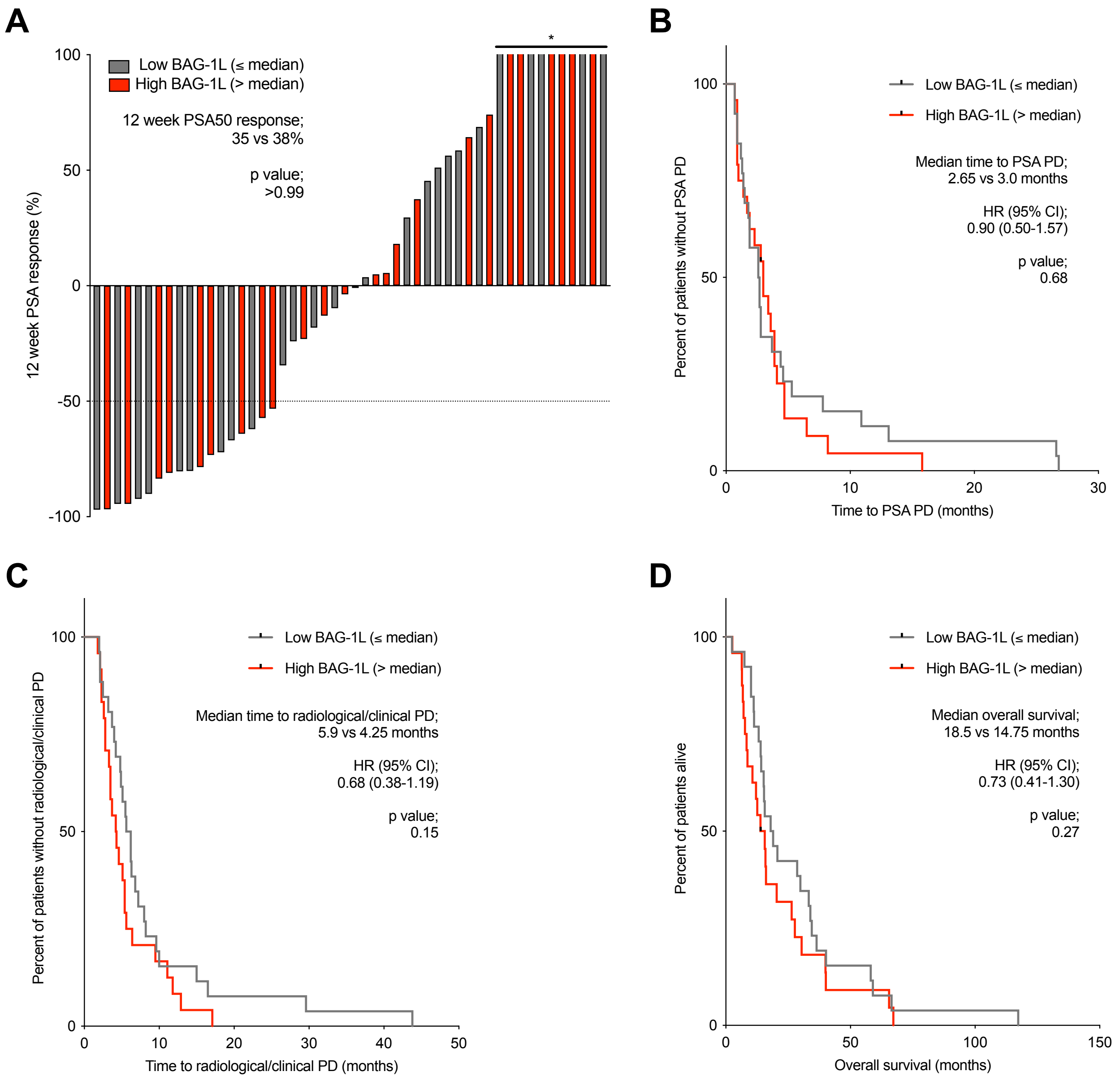
